# Long-fuse evolution of carnivoran skeletal phenomes through the Cenozoic

**DOI:** 10.1101/2025.07.14.664737

**Authors:** Chris Law, Leslea Hlusko, Jack Tseng

## Abstract

Climatic change is hypothesized to promote phenotypic diversification. While neontological analyses are often used to test this hypothesis, extant data only captures time-averaged signals of surviving lineages. More nuanced tests require paired and longitudinal climatic and organismal data. Here, we developed the most comprehensive phenomic dataset to-date of pan-carnivorans to test hypotheses that Cenozoic climatic change influenced the evolution of the cranial, appendicular, and axial skeleton. We found support for the hypothesis that a hierarchical progression of ecological diversification across the Cenozoic significantly influenced the establishment of modern carnivorans. Specifically, extinctions during the Eocene-Oligocene Transition released crown carnivorans from a constrained adaptive zone to interfamilial skeletal diversification. Intrafamilial skeletal diversification did not occur for another 20 million years until after the Mid-Miocene Climate Transition. Our work demonstrates the essential role of macroevolutionary data from the fossil record for revealing how major global climatic events steered the evolutionary trajectories of modern skeletal phenomes.

## Introduction

Ecological opportunities arising from climatic and environmental changes are often hypothesized to promote phenotypic diversification across macroevolutionary time scales [1–3]. After the Early Eocene Climate Optimum, trends towards decreased global temperatures [4] and consequent environmental transitions from closed forest habitats to open grassland habitats [5,6] have facilitated the increase in phenotypic diversity and evolution of novel phenotypes in several mammalian clades. For example, rodents, lagomorphs, and ungulates all independently evolved hypsodont dentition along with the lengthening of limbs for more efficient cursoriality in association with open habitats [7–10]. Similarly, phenotypic diversity in carnivores increased [11–13], enabling them to exploit diverse habitats through a range of hunting behaviors from ambush to cursorial or subterranean pursuit specialization [14–16]. However, the question of how these environmental changes influence the rate at which phenotypic disparity accumulates (i.e., tempo) and the pattern of phenotypic change across the phylogeny (i.e., mode) remains difficult to resolve due to the absence of time-sequence data (i.e., paleontological data) considered under a phylogenetic framework. The few studies that have successfully integrated the fossil record into neontological analyses demonstrated that fossils are essential for accurate model selection in phenotypic macroevolution analyses [17–21], thus providing a more informed understanding of phenotypic variation through phylogenetic time (e.g., [15,19,22–25]). To-date, these studies have primarily focused only on single phenotypic traits or structures like body size or the skull. No study has yet to simultaneously test how major environmental shifts may have facilitated the evolution of the skull, appendicular skeleton, and axial skeleton in both extant and extinct species.

In this study, we take advantage of the rich fossil record of pan-carnivorans (i.e., Hyaenodonta; Oxyaenodonta; and Carnivoramorpha, which includes Carnivora) to test hypotheses that climatic and environmental changes during the past 66 million years influenced the tempo and mode of the cranial, appendicular, and axial skeletal systems. Pan-carnivorans represent one of the best characterized crown mammal groups from the Cenozoic, with a crown-group fossil record going back more than 55 million years and stem taxa being traced back to the early Paleocene. With the extinction of non-avian dinosaurs, early pan-carnivorans became the first large carnivorous mammals [26]. Although crown carnivoran lineages co-existed with their now-extinct sister clades during much of the Eocene, most early carnivorans were small, generalized predators serving as secondary consumers. It was not until later in the Oligocene and Miocene that they expanded to large, specialized top predator niches [11–13,27,28]. Since their first appearance as small mongoose-like forms [11,13,17], carnivorans have diversified into the numerous body forms we see today, ranging from small, elongate weasels to big robust bears to fully aquatic, streamlined seals [29]. Here, we address the as-yet unanswered question of what drove the exceptional diversification of carnivoran body plans from a single blueprint.

The two leading hypotheses for the major influences on carnivoran morphological diversification are ecological opportunity facilitated by climate change and the extinction of coeval stem pan-carnivorans. Global cooling, increased aridity, and habitat shift from forest to grasslands have led to several rises and falls of carnivoran and stem pan-carnivoran clades [11]. Carnivorans survived during these environmental and climatic changes, as dental records suggest that crown carnivorans maintained their morphological diversity even during periods of extinction whereas morphological diversity in stem groups continued to decline [12]. Furthermore, stem pan-carnivorans exhibited steeper declines in species richness than carnivorans, and the overall percentage of stem pan-carnivoran taxa declined whereas those of carnivoran taxa increased [11–13]. These patterns have been attributed to competition between carnivorans and stem pan-carnivorans in which declining stem pan-carnivorans are replaced and excluded by phylogenetically distinct but functionally similar crown carnivorans [11–13,27]. Although analyses based on dental traits have partially supported this hypothesis [11,12], no quantitative evidence has been explicitly tested for this hypothesized link between paleoenvironmental changes and the diversification of the remaining skeletal phenome.

Here, we tested if increased carnivoran skeletal diversity was facilitated by rapid climate change, character displacement from stem ancestors, or independently from these factors. We predicted that the Eocene Oligocene Transition (EOT) and Mid-Miocene Climate Transition (MMCT) facilitated skeletal diversification within carnivorans, enabling species to exploit new resources during these periods of novel ecological opportunity. We leveraged the vast pan-carnivoran fossil record to quantify skeletal phenomes by incorporating traits from the skull, appendicular skeleton, and axial skeleton of extinct and extant species. We then used a phylogenetic comparative framework to examine the tempo and mode of skeletal evolution utilizing a total evidence phylogeny that contains almost all known extant and extinct carnivorans and their stem relatives [21].

## Results

### Skeletal trait imputation

We quantified the skeletal phenome of 118 extant species and 81 extinct species using 103 linear measurements obtained from 854 specimens held at 17 natural history museums (Table S1). Together, this 192 species dataset includes seven traits in the cranium; seven traits in the mandible; 13 traits in the forelimb; 13 traits in the hindlimb; and seven traits each in third cervical, fifth cervical, first thoracic, middle thoracic, diaphragmatic thoracic, last thoracic, first lumbar, middle lumbar, and last lumbar vertebrae (Fig. S1). We imputed missing trait measurements (25.2% of trait measurements in the full dataset; 0% in extant dataset; 61.9% in extinct dataset) using multiple imputation with PCA (MIPCA) under a Bayesian framework [30,31]. Our assessment of the quality of the trait imputations revealed one-to-one relationships (i.e., slopes are not significantly different from one) between imputed and empirical values of each trait regardless of the amount of missing data (Fig. S2A), indicating that our phenome dataset can reliably estimate missing trait data. Comparisons between log shape ratios calculated from imputed traits and empirical traits also show one-to-one relationships in most situations with 10–60% of randomly removed trait measurements (Fig. S2B).

We performed a principal component analysis (PCA) incorporating all trait measurements with size removed. Seven PC axes were found meaningful following procedures by [32]; these were subsequently used as our proxy of the skeletal phenome (76.3% of explained variance; Fig. 1; see Table S2 for trait loadings). The geometric mean of all trait measurements was used as our proxy for skeletal size. We similarly performed PCAs to reduce the dimensions of each skeletal component (i.e., cranium, mandible, forelimb, hindlimb, and each of the nine vertebrae).

**Fig. 1.**
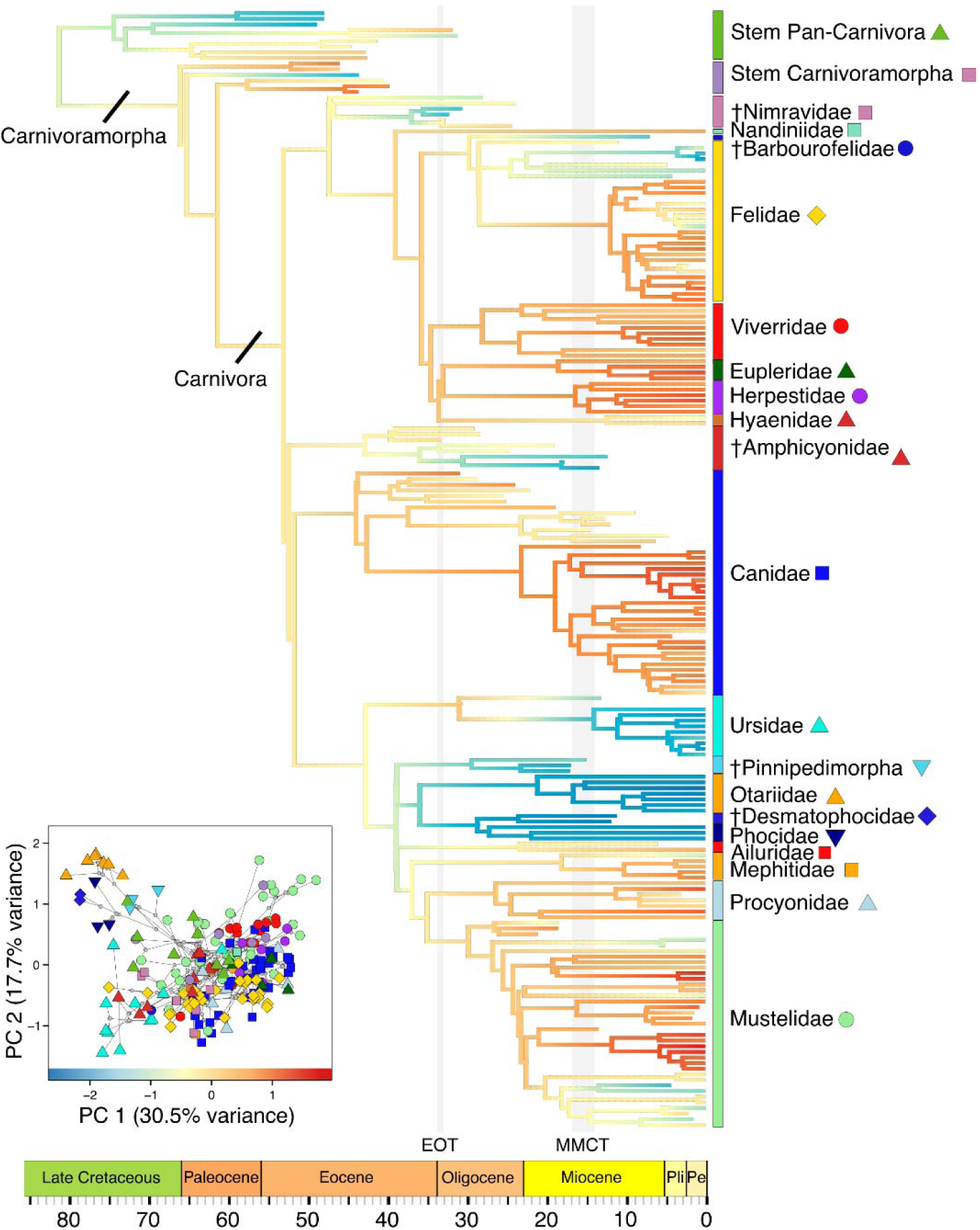
Pruned pan-carnivoran phylogeny with ancestral reconstruction of PC 1 of skeletal phenome, calculated from all 103 size-corrected skeletal traits.

### Evolution of carnivoran skeletal phenome and size

Phylogenetic signal in carnivoran skeletal phenome was moderate (Pagel’s λ = 0.60, p = 0.001; Blomberg’s K = 0.58, p = 0.001), suggesting that disparity in skeletal phenome primarily occurs within clades as opposed to among clades. Although disparity through time (DTT) analyses indicated that subclade disparity in skeletal phenome does not differ from what is expected under Brownian motion (BM; morphological disparity index [MDI] = 0.018, P = 0.632), further inspection revealed higher subclade disparity than expected under BM during two time intervals. The first occurs from ∼39 to ∼37 Ma (million years ago; MDI = 0.534, P = 0.012), right between the Middle Eocene Climatic Optimum 40 Ma and the Eocene-Oligocene Transition 34 Ma (Fig. 2A,B). The second occurs from ∼14 to ∼1 Ma, coinciding with the conclusion of the Middle Miocene Climatic Optimum and the Mid-Miocene Climate Transition ∼14 Ma (MDI = 0.238, P = 0.018) (Fig. 2C).

**Fig. 2.**
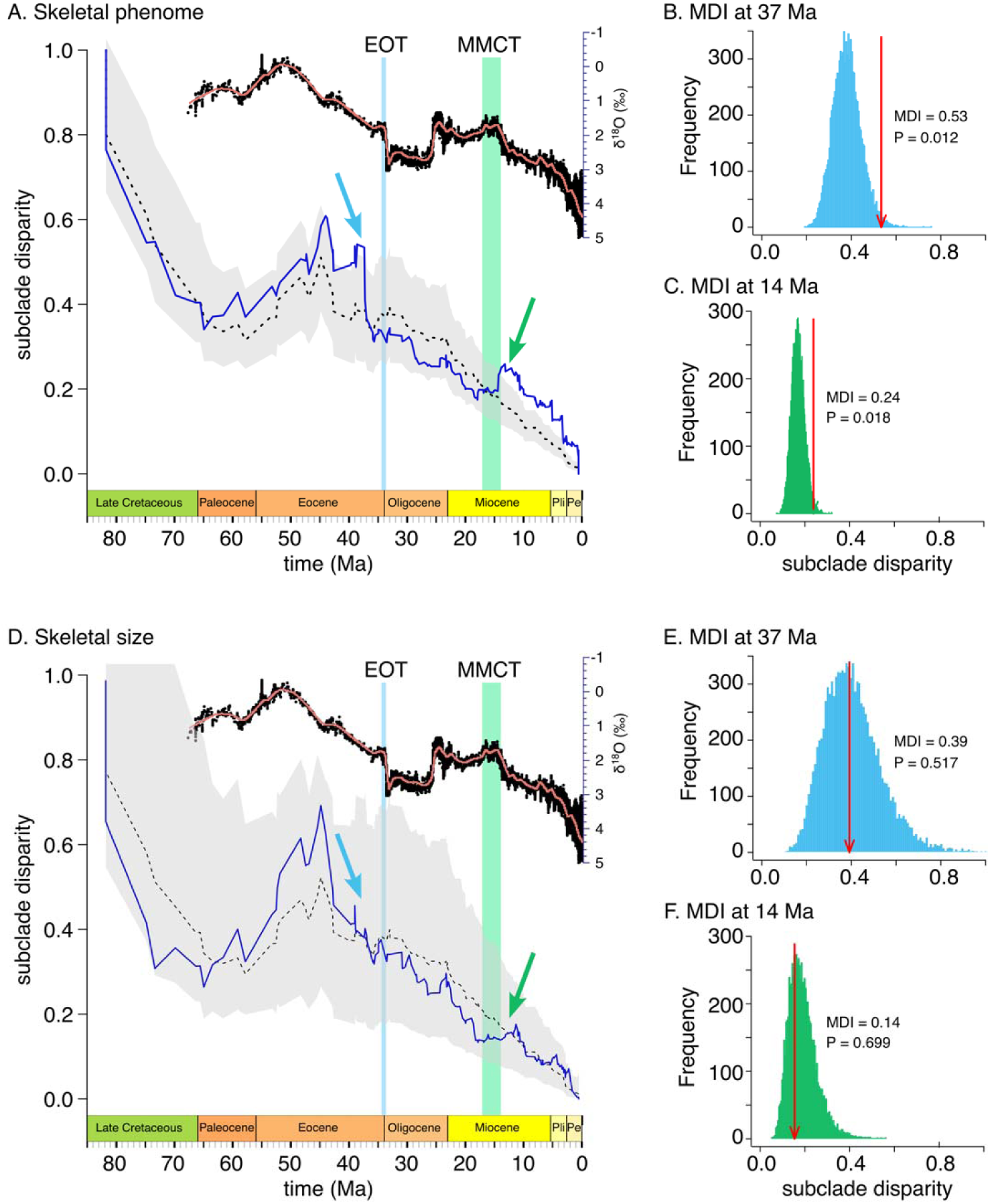
Disparity-through-time plots of carnivoran skeletal phenome and size. (A) Skeletal phenome deviates from Brownian motion 37 Ma (blue arrow) near the beginning of the Eocene-Oligocene Transition (EOT) and 14 Ma near the end of the Mid-Miocene Climate Transition (MMCT). Blue solid line indicates the empirical subclade DTT, and the dashed black line indicates the median DTT based on 10,000 simulations of trait evolution under Brownian motion. The grey shaded area indicates the 95% DTT range for the simulated data. Above th DTT is the global deep-sea oxygen isotope records as a proxy for global temperature change (modified from [4]). DTT suggests increased subclade disparity at two time points: (B) just prior to the EOT (∼39-37 Ma; MDI = 0.534, P = 0.012) and (C) just after the MMCT (∼14 Ma; MDI = 0.238, P = 0.018). For B. and C., histograms show the distribution of simulated subclad disparities and red arrows signify the empirical subclade disparities at time 37 Ma and 14 Ma, respectively. (D) Skeletal size does not deviate from Brownian motion across time and at (E) ∼37 Ma and (F) ∼14 Ma.

We tested 16 alternative hypotheses that could explain the disparity of carnivoran skeletal phenome by fitting a series of macroevolutionary models (Fig. 3). We found nearly equal support for the two release and radiate models (Table 1), where the evolution of skeletal phenomes transitions from an Ornstein–Uhlenbeck (OU) process to BM at the EOT 34 Ma (OUBMi_EOT_: AICcW = 0.66) and at the MMCT 14 Ma (OUBMi_MMCT_: ΔAICc = 1.33, AICcW = 0.34). These models indicated that evolutionary rates slowed after both the EOT (mean σ^2^ = 0.148; mean σ^2^ = 0.013) and the MMCT (mean σ^2^ = 0.048; mean σ^2^ = 0.013). Mean stationary variances (σ^2^/2α) for the OU portion of the OUBMi_EOT_ and OUBMi_MMCT_ models were 0.148 (α = 0.50) and 0.285 (α = 0.08), respectively. In contrast, models with distinct optima among pan-carnivoran groups (i.e., sOU and cOU models) were poorer fits (ΔAICc = 89.80– 107.95), even compared to single regime models based on BM or OU. Simulations under the parameters of the OUBMi_EOT_ model indicated that there was substantial power to distinguish this model from BM and alterative models (AICcW > 0.99; Table S3; Fig. 4A). Simulations under the parameters of the OUBMi_MMCT_ model indicated that there was substantial power to distinguish this model from all alterative models (ΔAICc = 1.96; AICcW = 0.27; Table S3; Fig. 4B) except the BMBM_MMCT_ (AICcW = 0.73; Table S3). These simulations thus suggest that the EOT rather than the MMCT may have a stronger effect on shifts of evolutionary mode of skeletal phenome.

**Fig. 3.**
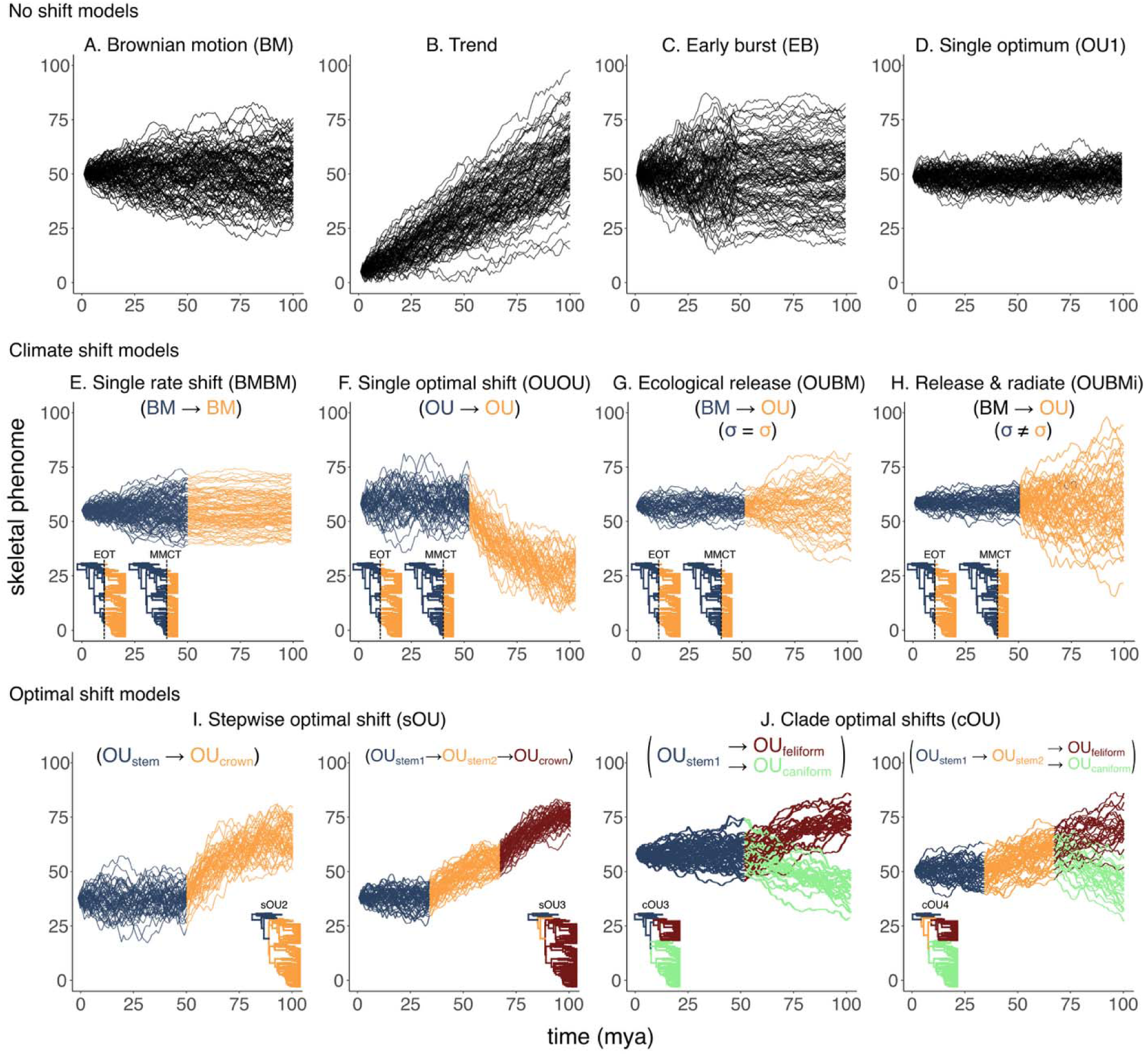
Alternative hypotheses for the evolution of carnivoran skeletal phenomes. (A) Brownian motion. (B) Brownian motion with a directional drift. (C) Brownian motion with an evolutionary rate that increases or decreases exponentially through time. (D) Single optimum under an Ornstein-Uhlenbeck process. (E) Single rate shift after the Eocene-Oligocene Transition (EOT) or Mid-Miocene Climate Transition (MMCT). (F) Single optimal shift after the EOT or MMCT. (G) Constrained evolution followed by unconstrained evolution after the EOT or MMCT. (H) Constrained evolution followed by unconstrained evolution after the EOT or MMCT with distinct evolutionary rates. (I) Stepwise optimal shifts between carnivorans and its stem relative (sOU2) or among carnivorans, stem carnivoramorphs, and stem pan-carnivorans (sOU3). (J) Stepwise optimal shifts followed by clade optimal shifts among caniforms, feliforms, and its stem relatives (cOU3) or among caniforms, feliforms, stem carnivoramorphs, and stem pan-carnivorans (cOU4).

**Fig. 4.**
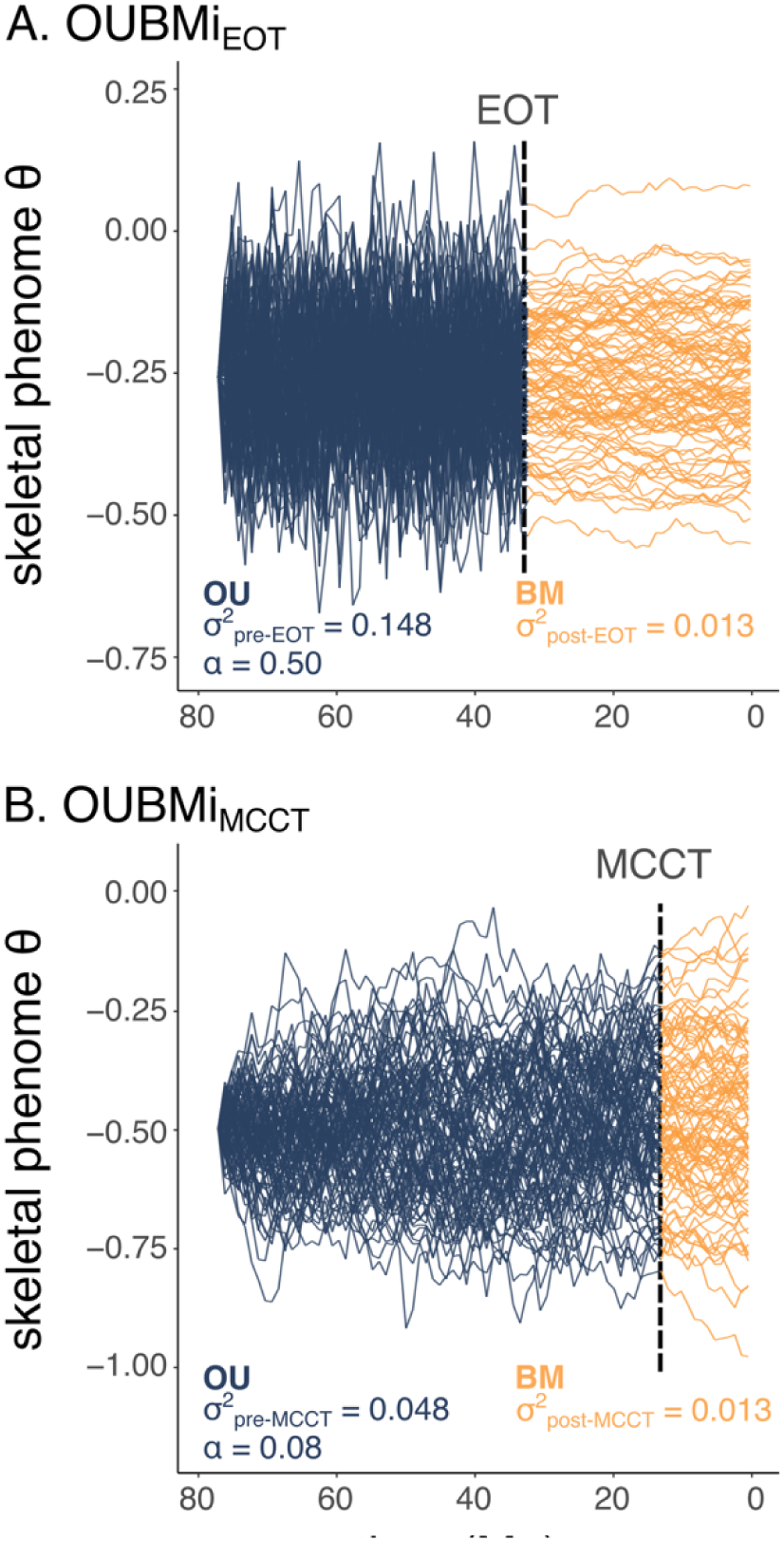
Simulations of the parameter estimates of the best supported models for skeletal phenome. 100 simulations were used for visualization.

**Table 1.** Comparisons of evolutionary models for skeletal phenome and skeletal size. Rows in boldface type represent the best supported model(s) as indicated by the lowest ΔAICc score. Models with ΔA<u>ICc < 2 were considered as equally well supported. AICcW = A</u>ICc weights.

| Trait | Model | AICc | $\Delta\text{AICc}$ | AICcW |
| --- | --- | --- | --- | --- |
| skeletal phenome | BM1 | 883.53 | 51.98 | 0.00 |
|  | trend | 880.84 | 49.30 | 0.00 |
|  | EB | 891.17 | 59.62 | 0.00 |
|  | OU1 | 907.73 | 76.18 | 0.00 |
|  | BMBMi<sub>EOT</sub> | 849.15 | 17.60 | 0.00 |
|  | OUOU<sub>EOT</sub> | 890.45 | 58.91 | 0.00 |
|  | OUBM<sub>EOT</sub> | 883.35 | 51.81 | 0.00 |
|  | **OUBMi<sub>EOT</sub>** | **831.55** | **0.00** | **0.66** |
|  | BMBMi<sub>MMCT</sub> | 859.67 | 28.12 | 0.00 |
|  | OUOU<sub>MMCT</sub> | 898.95 | 67.40 | 0.00 |
|  | OUBM<sub>MMCT</sub> | 879.37 | 47.82 | 0.00 |
|  | **OUBMi<sub>MMCT</sub>** | **832.87** | **1.33** | **0.34** |
|  | sOU2 | 921.34 | 89.80 | 0.00 |
|  | sOU3 | 931.93 | 100.38 | 0.00 |
|  | cOU3 | 939.50 | 107.95 | 0.00 |
|  | cOU4 | 926.50 | 94.95 | 0.00 |
| skeletal size | BM1 | 168.80 | 6.09 | 0.03 |
|  | trend | 167.63 | 4.93 | 0.05 |
|  | EB | 170.86 | 8.15 | 0.01 |
|  | OU1 | 172.89 | 10.19 | 0.00 |
|  | BMBMi<sub>EOT</sub> | 168.31 | 5.61 | 0.04 |
|  | OUOU<sub>EOT</sub> | 168.48 | 5.78 | 0.03 |
|  | OUBM<sub>EOT</sub> | 165.88 | 3.18 | 0.12 |
|  | OUBMi<sub>EOT</sub> | 167.92 | 5.22 | 0.04 |
|  | BMBMi<sub>MMCT</sub> | 171.14 | 8.44 | 0.01 |
|  | OUOU<sub>MMCT</sub> | 171.89 | 9.19 | 0.01 |
|  | OUBM<sub>MMCT</sub> | 166.69 | 3.98 | 0.08 |
|  | **OUBMi<sub>MMCT</sub>** | **162.70** | **0.00** | **0.58** |
|  | sOU2 | 173.31 | 10.61 | 0.00 |
|  | sOU3 | 175.23 | 12.52 | 0.00 |
|  | cOU3 | 175.31 | 12.61 | 0.00 |
|  | cOU4 | 177.37 | 14.66 | 0.00 |

In contrast, high phylogenetic signal (Pagel’s λ = 0.91, p = 0.001; Blomberg’s K = 0.56, p = 0.001) and DTT analyses were united in showing that disparity in skeletal size did not significantly deviate from BM and tends to occur within clades (Fig. 2D–F). The OUBMi_MMCT_ model was the best supported model (AICcW = 0.58) for skeletal size, suggesting that the evolution of skeletal size transitions from an OU process (α = 0.05; σ^2^ = 0.019) to BM (σ^2^_post-MMCT_ = 0.008) during the MMCT (Table 1). However, simulations could not distinguish this best supported model from OU1 (AICcW = 0.21), cOU3 (AICcW = 0.16), OUOU_EOT_ (AICcW 0.14), sOU2 (AICcW = 0.09) and the OUOU_MMCT_ (AICcW = 0.09) models (Table S3). Thus, climatic changes may have had less of an impact on the evolution of skeletal size than suggested by macroevolutionary models.

### Evolution of carnivoran skeletal components

Phylogenetic signal of the skeletal components ranged from moderate to high (Pagel’s λ = 0.44–0.94, all p = 0.001; Blomberg’s K = 0.38–0.45, all p = 0.001; Table S4), suggesting that variance in skeletal components tended to follow BM and primarily occurred within clades as opposed to among clades. DTT analyses revealed three general patterns in the evolution of these skeletal components (Fig. S3). First, all vertebrae largely followed BM expectations except during last 10–20 million years (My). Second, the cranium and limbs exhibited elevated subclade disparity compared to BM during the Middle Eocene Climatic Optimum 40 Ma and from the Early Miocene to present. Third, the mandible exhibited elevated subclade disparity throughout the majority of pan-carnivoran history.

Climate models with shifts in evolutionary rate, mode, and/or optima were the best fitting models for all skeletal components (Table S5; see Table S6 for parameter estimates from best model). The OUBMi_MMCT_ model was best supported for the hindlimb and most vertebrae (AICcW = 0.51–1.00; Table S5). Simulations under the parameters of this model indicated that there was substantial power to distinguish from alterative models for these components except for the last thoracic and first and middle lumbar vertebrae (Table S7). For the cranium, the BMBM_EOT_ (AICcW = 0.37), OUBMi_MMCT_ (ΔAICc = 0.59, AICcW = 0.28), and OUBM_EOT_ (ΔAICc = 1.43, AICcW = 0.18) models were equally supported; however, simulations could not distinguish these models from BM (Table S7). The OUOU_EOT_ (AICcW = 0.54) and OUBMi_MMCT_ (ΔAICc = 1.32; AICcW = 0.28) models were equally supported for the mandible, and simulations indicated there was substantial power to distinguish them from BM and alterative models (Table S7). The OUOU_MMCT_ model was best supported for the forelimb (AICcW = 0.84), and the OUBM_EOT_ model was best supported for the third cervical vertebrae (AICcW = 0.69); simulations indicated substantial power to distinguish these models from BM in both scenarios. Lastly, the BMBM_EOT_ and OUBMi_EOT_ models were equally supported for the first thoracic and middle thoracic vertebrae (AICcW = 0.30–0.81), but simulations could not distinguish these models from BM and EB models (Table S7).

## Discussion

The global cooling, increased aridity, and habitat shift from forests to grasslands during the past 56 My has spurred phenotypic diversification in many mammalian clades [7–10,14,15]. Here, we found that climatic and environmental changes from the Eocene-Oligocene Transition (EOT) and Mid-Miocene Climate Transition (MMCT) significantly facilitated the diversification of crown carnivoran skeletal phenomes across two distinct phases.

In the initial phase, EOT-induced climatic and environmental changes along with the decline of coeval stem pan-carnivorans released crown carnivorans from a constrained adaptive zone to radiation under Brownian motion, albeit at a slower evolutionary rate (Fig. 4A). With the onset of the EOT, global temperatures plummeted and the first Antarctic ice sheets appeared, transitioning Earth’s climate from a warm, equable “greenhouse” to a cooler, temperate “icehouse” with increased seasonality [33–35]. This period of climate change led to habitat transitions from warm-humid forest to dry-temperate forest-steppe and grasslands, which in turn led to massive faunal turnover [34–39]. Among pan-carnivorans, the EOT led to tremendous taxonomic turnover, resulting in the decline of Hyaenodonta and Oxyaenodonta and giving opportunities for carnivoran to radiate [11,40] under a BM pattern (Fig. 4A). Our analyses suggesting EOT-induced interfamilial diversification is consistent with paleontological records indicating that early carnivorans exhibited generalized, small-bodied forms and did not exhibit major lineage and phenotypic diversification until after the EOT [11,12,27]. Niche competition from stem pan-carnivorans may have initially constrained early carnivoran diversification prior to the EOT [11–13,27]. However, the EOT marked the decline of stem pan-carnivorans [11,40], possibly because they had become too specialized (i.e., macroevolutionary ratchet) to adapt to the climatic and environmental changes brought by the EOT [11,12]. Thus, although both stem pan-carnivorans and early carnivorans exhibited a macroevolutionary ratchet towards large body size and hypercarnivory during this time, only carnivorans retained and increased their phenotypic diversity, giving them the opportunity to exploit habitats of the Oligocene [12,13]. This may have further excluded stem pan-carnivorans from novel ecological opportunities [11–13,27,28], facilitating stem pan-carnivoran decline but carnivoran diversification and the appearance of all modern carnivoran families from the Early Oligocene to the Mid-Miocene [11]. Poor AICc support for all the optima shift models supports this hypothesis (Table 1). The lack of distinct optima indicates pan-carnivoran groups exhibit similar skeletal phenomes and ecomorphologies, further supporting that carnivorans displaced and replaced stem pan-carnivorans.

Further intrafamilial diversification of carnivoran skeletal phenomes did not occur for another 20 My until after the MMCT. During this second phase, another major period of rapid temperature decline, increased aridity, and enhanced seasonality facilitated further trends towards grasslands from forest habitats [6,41–43]. Our results suggested that these rapid climatic and environmental changes along with few competing clades of carnivorous mammals enabled crown carnivorans to exploit the new ecological niches from the late Miocene to Pleistocene [11–13,28]. Specifically, intrafamilial diversification of skeletal phenomes may have facilitated resource partitioning among ecologically similar taxa by increasing ecomorphological diversity or innovations in locomotor mode, hunting behavior, and dietary ecology [15,44–46]. Consistently, the appendicular skeleton and posterior region of the vertebrae column, two skeletal components that exhibit strong relationships with locomotor and hunting behavior evolution [45], were characterized by models that incorporate shifts in evolutionary rate and/or mode at the MMCT (Table S5). This suggests that evolutionary shifts or radiation of novel traits in the appendicular and axial skeleton coincided with environmental changes of the Mid-Miocene. For example, elbow-joint morphologies indicative of pounce-pursuit behaviors evolved by the late Miocene within canids [15], presumably in response to the Mid-Miocene diversification of grazing ungulates in open habitats [47,48]. Similarly, small, elongate body plans with short legs evolved within mustelids [16,46], presumably to exploit the Mid-Miocene diversification of rodents in open habitats, both above and subterranean [7,49]. More recent work with extant carnivorans indicated that specific skeletal components exhibited strong selection towards these distinct ecological peaks, and functional trade-offs and covariance among and within skeletal components further explain their distribution across the adaptive landscape [45,50,51]. Although multi-optima OU models with distinct ecological peaks may better capture the evolutionary mode of these skeletal components, we are unable to test these models without ecological data from extinct species. Unfortunately, ecological classifications are typically inferred using the same data (i.e., skeletal traits) used in our analyses, which may lead to biased results if the inferred ecologies of extinct species are used to test for ecological peaks.

Multi-phasic evolution driven by climatic and environmental changes and coupled with taxonomic turnover appears to characterize the evolution of placental mammals. The delayed intrafamilial diversification in carnivorans is reminiscent of the long fuse model used to describe the diversification of placental mammals, which posits that the initial interordinal diversification of placental mammals occurred in the Cretaceous whereas the majority of intraordinal diversification took place after the K-Pg mass extinction event in response to newly available niche [52–54]. The Paleocene–Eocene Thermal Maximum (PETM) may have also facilitated distinct phases of mammalian evolution, albeit this has yet to be quantitatively tested. The rise of forested environments and warmer climates after the PETM resulted in the replacement of stem lineages of the early Paleocene such as multituberculates, pantodonts, and condylarths by crown mammals such as primates, perissodactyls, and artiodactyls [55–57]. This turnover led to the evolution of crown mammalian orders that dominated the Cenozoic and today’s ecosystems [56]. Together, this body of work suggests that sequential evolutionary phases triggered by climatic and environment changes were largely hierarchical. For carnivorans, the hierarchical progression of ecological diversification across the Cenozoic led to the establishment of modern species.

Specifically, climatic and environmental changes from the K-Pg, PETM, EOT, and MMCT facilitated diversification of infraclass Placentalia, mirorder Pan-Carnivora, families within Carnivora, and genera within carnivoran families, respectively. Thus, our results suggest that the long-fuse EOT/MMCT pattern we report here is as consequential for establishing modern-day carnivore guilds as the PETM were for establishing the modern orders of mammals. Overall, this work highlights the importance of analyzing data from extant and extinct species under a phylogenetic comparative framework to disentangle the multiple climatic events that contributed to the tiered evolution and diversity of carnivoran skeletal phenomes. Genetic analyses that improve our assessment methods of skeletal variation will bring even more insight [58–60]. We posit that quantification of skeletal phenomes in other crown mammals and their extinct relatives will similarly reveal the effect of Cenozoic climatic change on the multi-phasal, hierarchical establishment of modern-day mammalian biodiversity.

## Methods

### Morphological data

We quantified the skeletal phenome of 118 extant species and 81 extinct species using 103 linear trait measurements obtained from 208 extant specimens and 646 fossils (see Table S1 for list of specimens and museums). Together, this dataset includes seven traits in the cranium; seven traits in the mandible; 13 traits in the forelimb (scapula, humerus, ulna, radius, third metacarpal); 13 traits in the hindlimb (pelvis, femur, tibia, fibula, calcarean, and third metatarsal); and seven traits in third cervical, fifth cervical, first thoracic, middle thoracic (the sixth thoracic if the species has 13 or 14 thoracic vertebrae or the seventh thoracic if the species has 15 or 16 thoracic vertebrae), diaphragmatic thoracic, last thoracic, first lumbar, middle lumbar (the third lumbar if the species has four, five, or six lumbar vertebrae or the fourth lumbar if the species has seven lumbar vertebrae), and last lumbar vertebrae (Fig. S1). These measurements were chosen based on previous investigations of distinct ecomorphology among carnivorans [23,61–64]. All measurements were taken using Mitutoyo digital calipers or a tape measure. All specimens were fully mature, determined by the closure of exoccipital-basioccipital and basisphenoid-basioccipital sutures on the cranium, full tooth eruption, or closure of sutures on the limb bones or vertebrae. In our extant dataset, we obtained representatives from 14 of the 16 carnivoran families. We were unable to collect data from Prionodontidae, which contains two extant species of Asiatic linsangs, and Odobenidae, which contains a single extant species, the walrus. We only used male specimens to limit the differing degrees of sexual dimorphism that carnivorans exhibit [65,66]. In our extinct dataset, we obtained representatives from seven extant carnivoran families, seven extinct carnivoran families, stem carnivoramorphs, and stem pan-carnivorans (Hyaenodontidae and Oxyaenidae). We were limited by the availability of fossils that were formally described, placed in a phylogenetic context, and free from taphonomic or other deformation.

Due to the fragmented nature of the fossil record, 73 extinct species were missing trait measurements, resulting in 25.2% missing data in the full dataset (61.9% in extinct dataset). We, therefore, imputed missing trait measurements using MIPCA (multiple imputation with PCA) in the R package missMDA v1.19 [30]. MIPCA predicts missing trait measurements using an iterative PCA algorithm with a predefined number of dimensions. The number of dimensions was estimated by generalized cross-validation criterion, and the iterative algorithm was performed under a Bayesian framework that alternates imputation of the data set and draw of the PCA parameters in a posterior distribution [31].

We assessed the quality of the trait imputations by comparing imputed and empirical data using a subset of species (n = 124) that contain all 103 measured traits. Specifically, we generated four datasets with different percentages (10%, 30%, 60%, and 80%) of randomly removed trait measurements and used MIPCA to impute these removed traits. We then assessed the accuracy of MIPCA by plotting the imputed traits against the empirical average traits and conducting reduced major axis regressions to test if the slope significantly differs from one. We found strong relationships between imputed traits and empirical traits where the slope is not significantly different from one for all datasets with different percentages of removed traits (see Fig. S2 for two trait examples).

Using the imputed dataset, we removed the effects of size by dividing each skeletal trait by the geometric mean of all 103 traits (i.e., log shape ratio: ln(trait/geometric mean) [67,68]. We then conducted a principal component analysis (PCA) on the entire dataset and retained the first seven PC axes (76.3% of explained variance) as our proxy of skeletal phenome. The number of PC axes to retain was determined using the getMeaningfulPCs function in the R package Morpho v2.11 [69]. We also used individual PCAs to reduce the dimension of each skeletal component (i.e., log shape ratios from the cranium, mandible, forelimb, hindlimb, and each of the nine vertebrae) and retained meaningful PC axes for subsequent analyses.

### Phylogenetic data

We used a modified species-level phylogeny containing almost all known extant and extinct carnivorans and stem relatives for our phylogenetic comparative methods [21]. After pruning, the phylogeny contained all 118 extant species and 73 of the 81 extinct species in our dataset. We manually added the following extinct species: *Crassidia intermedia* was placed as basal caniform and sister to the genus *Daphoenus* [70]; *Corumictis wolsani* was placed as stem to Mustelidae and sister to the genus *Promartes* [71]; *Cyonasua brevirostris* was placed as sister to genus *Nasua* [72]; *Eoarctos vorax* was placed as sister to Ursidae [73]; *Puma pardoides* was placed as sister to *Puma concolor* [74]; *Satherium ingens* was placed as sister to sister to genus *Pteronura* [75]; *Neovulpavus mccarrolli* was placed as a stem carnivoramorphan and sister to *Miacis washakius* [76]; and *Promartes olcotti* was placed as a stem to Mustelidae and sister to the genus *Corumictis* [77] (Table S8).

### Phylogenetic Comparative Methods

We first measured phylogenetic signal in skeletal phenome, skeletal size, and each skeletal component using the physignal.z function in R package geomorph v4.0.6 [78]. We then examined subclade disparity in each dataset of retained PC axes by performing disparity through time (DTT) analyses [79] using fdtt [22], a modified version of the dtt function in the R package geiger v2.0.11 [80]. We compared carnivoran DTT to a null, constant-rates distribution derived from 10,000 Brownian motion simulations, and computed the morphological disparity index (MDI) as the area between the empirical DTT curve and the median of these simulations.

We then tested 16 alternative hypotheses that could explain carnivoran skeletal evolution by fitting a series of macroevolutionary models (Fig. 3). Models 1–4 are based on single rate Brownian motion or single optimum Ornstein-Uhlenbeck process:

1. Brownian motion (BM): Suggests that disparity in skeletal phenome accumulates proportionally to evolutionary time under a random walk.
2. Trend: Suggests that disparity in skeletal phenome accumulates under BM with a directional drift.
3. Early burst (EB): Suggests that disparity in skeletal phenome initially accumulates rapidly but declines over time.
4. Single optimum Ornstein-Uhlenbeck process (OU1): Suggests that disparity in skeletal phenome is constrained or under selection to a single optimum over time. Models 5–12 are climate shift models to test if evolutionary rates, skeletal optima, and/or evolutionary mode differs before and after the Eocene Oligocene Transition (EOT) and Mid Miocene Climate Transition (MMCT):
5. Rate shift after the EOT (BMBM_EOT_). Suggests that skeletal phenome exhibits two distinct evolutionary rates before and after the EOT.
6. Rate shift after the MMCT (BMBM_MMCT_). Similar to the BMBM_EOT_ model, but the shift occurs at the MMCT.
7. Optima shift after the EOT (OUOU_EOT_). Suggests that skeletal phenome exhibits two distinct evolutionary optima before and after the EOT with the same evolutionary rate.
8. Optima shift after the MMCT (OUOU_MMCT_). Similar to the OUOU_EOT_ model, but the shift occurs at the MMCT.
9. Ecological release after the EOT (OUBM_EOT_). Suggests that skeletal phenomes are constricted in ancestral carnivorans via an OU model due to extrinsic factors associated with the EOT, followed by the increased skeletal disparity after the release from these factors via BM.
10. Ecological release after the MMCT (OUBMi_MMCT_). Similar to the OUBM_EOT_ model, but the shift occurs at the MMCT.
11. Release and radiate after the EOT (OUBMi_EOT_). Similar to the OUBM_EOT_ model, except the evolutionary rate also shifts after the EOT.
12. Release and radiate after the MMCT (OUBMi_MMCT_). Similar to the OUBMi_MMCT_ model, except the evolutionary rate also shifts after the MMCT. Models 13–16 are optima shift models to test if carnivorans and stem relatives exhibit separate optima. Because ecological factors influence several aspects of the carnivoran skeleton [44,45,64,81], distinct skeletal phenomes may potentially indicate a reduction of ecological competition through character displacement:
13. Stepwise optimal shift (sOU2): Suggests that carnivorans and its stem relatives exhibit separate skeletal phenome optima via a two-peak OU model.
14. Stepwise optimal shift (sOU3): Suggests that carnivorans, stem carnivoramorph relatives, and stem pan-carnivoran relatives exhibit separate skeletal phenome optima via a three-peak OU model.
15. Clade optimal shift (cOU3): Suggests that feliforms and caniforms exhibit separate skeletal phenome optima after an initial optimal shift from stem relatives via a three-peak OU model.
16. Clade optimal shift (cOU4): Suggests that feliforms and caniforms exhibit separate skeletal phenome optima after initial optimal shifts from stem carnivoramorph relatives and stem pan-carnivoran via a four-peak OU model.

All models were fit to skeletal phenome, skeletal size, and each skeletal component datasets with measurement error incorporated using the R package mvMORPH v1.1.7 [82]. We did not have complete intraspecific data to calculate standard errors for each species. Instead, we estimated standard errors of each species using the following procedure. First, we estimated the standard deviation of each PC axis of each species by multiplying the range of each PC axis with 5%. This approximation assumes that on average each species would have an estimated standard deviation of 5% of the maximum range of the full species range on that axis. We then estimated the standard error by dividing the estimated standard deviation with the square root of the total number of species. Optima shifts in the sOU and cOU models were specified on the phylogeny using the paintSubTree function in the R package phytools v2.4.4 [83]. Relative support for each of the 16 models was assessed through computation of small sample-corrected Akaike (AICc). Models with ΔAICc < 2 were considered as equally well supported. We assessed whether we had adequate power to accurately distinguish between complex models from Brownian motion using simulations [84]. We generated 500 simulations for all datasets using the parameter estimates of the best-supported model(s) from the empirical datasets. We then ran each set of simulated data through all 16 models to determine whether the best model(s) could be accurately recovered.

As a sensitivity analysis, we also fit all 16 models to the first seven axes from a phylogenetic PCA of skeletal phenome (56.6% of explained variance). We found nearly equal support for the two release and radiate models (OUBMi_EOT_: AICcW = 0.74; OUBMi_MMCT_: ΔAICc = 2.06, AICcW = 0.26; Table S9), a result similar to model selection using a standard PCA. We chose not to continue with the phylogenetic PCA because pPCA is more difficult to interpret as it is a mixture of major axes that describe non-phylogenetic variation and scores that contain phylogenetic components of variation [85]. Furthermore, pPCA axes are not orthogonal to each other, so the first few axes we used in this study may include less variance explained than PCA by containing correlated variance components rather than independent ones.

## Acknowledgments

On title page

## Data accessibility

Dataset and R script are uploaded on Dryad Digital Repository: doi.org/10.5061/dryad.z8w9ghxrz.

