## Supplementary Materials for "Long-fuse evolution of carnivoran skeletal phenomes through the Cenozoic"

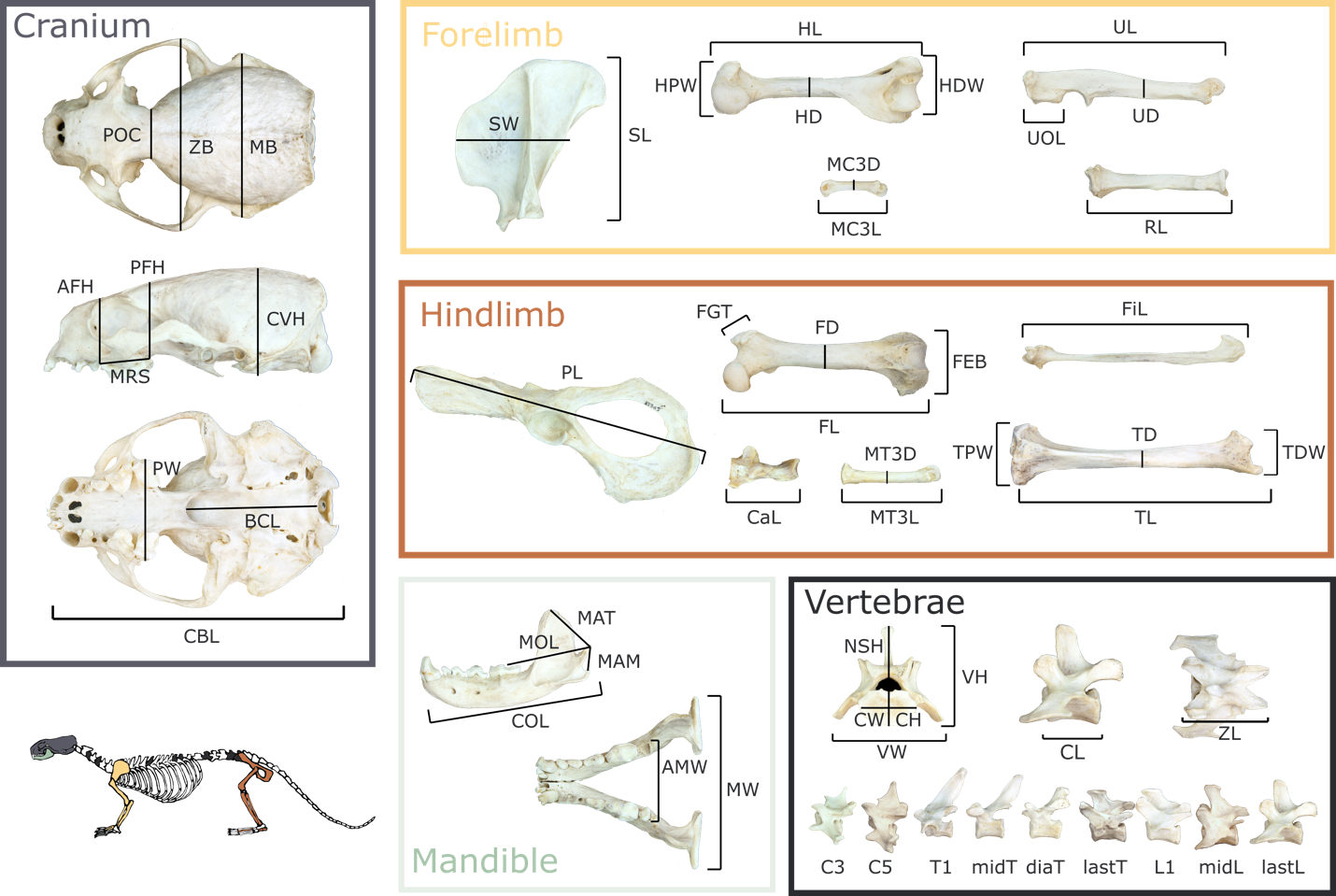

**Fig. S1. Skeletal trait measurements used in this study**. *Cranium*: CBL = condylobasal length, POC = postorbital constriction breadth, ZB = zygomatic breadth, MB = mastoid breadth, AFH = anterior facial height, PFH = posterior facial height, CVH = cranial vault height, PW = palate width, BCL = basicranial length. *Mandible*: MAT = moment arm of temporalis, MAM = moment arm of masseter, MAM2 = moment arm of masseter; MOL = molar out-lever; COL = canine out-lever; MW = mandibular width; AMW = anterior mandibular width. *Forelimb*: SL = scapula length; SW = scapula width; HL = humerus length; HD = humerus mid-shaft width; HPW = humerus proximal width; HDW = humerus distal width; UL = ulna length; UD = ulna mid-shaft width; UOL = ulnar olecranon length; RL = radius length; RD = radius mid-shaft width; MC3L = third metacarpel length; MC3W = third metacarpel width. *Hindlimb*: PL = pelvis length; FL = femur length; FD = femur mid-shaft width; FEB = femur distal width; FGT = height of the greater trochanter of the femur; TL = tibia length; TD = tibia mid-shaft width; TPW = tibia proximal width; TDW = tibia distal width; FiL = fibula length; CaL = calcaneus length; MT3L = third metatarsal length; MT3D = third metatarsal width. *Vertebrae*: VW = vertebrae width; VH = vertebrae height; CL = centrum length; CW = centrum width (posterior); CH = centrum height (posterior); ZL = inter-zygapophyseal length; NSH = neural spine height.

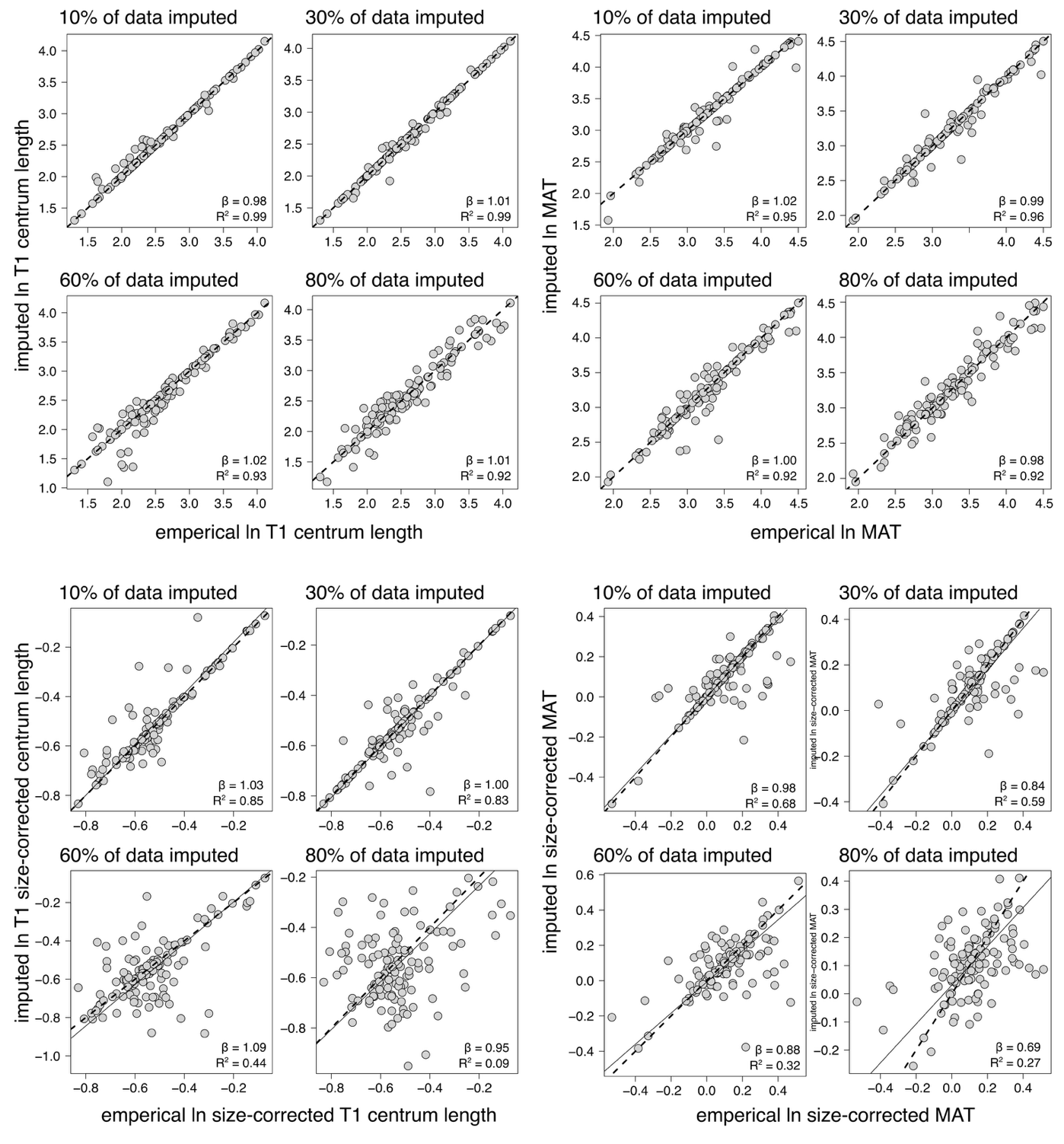

**Fig. S2. Relationships between imputed and empirical trait values to assess the quality of trait MIPCA imputation.** Assessments were performed using a subset of species (n = 192) that contain all 103 measured traits and randomly removing different percentages (10%, 30%, 60%, and 80%) of trait measurements. MIPCA was used to impute these removed traits, and reduced major axis regression models were used to test the relationship between imputed traits and empirical traits. Black dashed line indicates the 1:1 line. β = slope obtained from RMA models. T1 = first thoracic vertebrae. MAT = moment arm of the temporalis muscle.

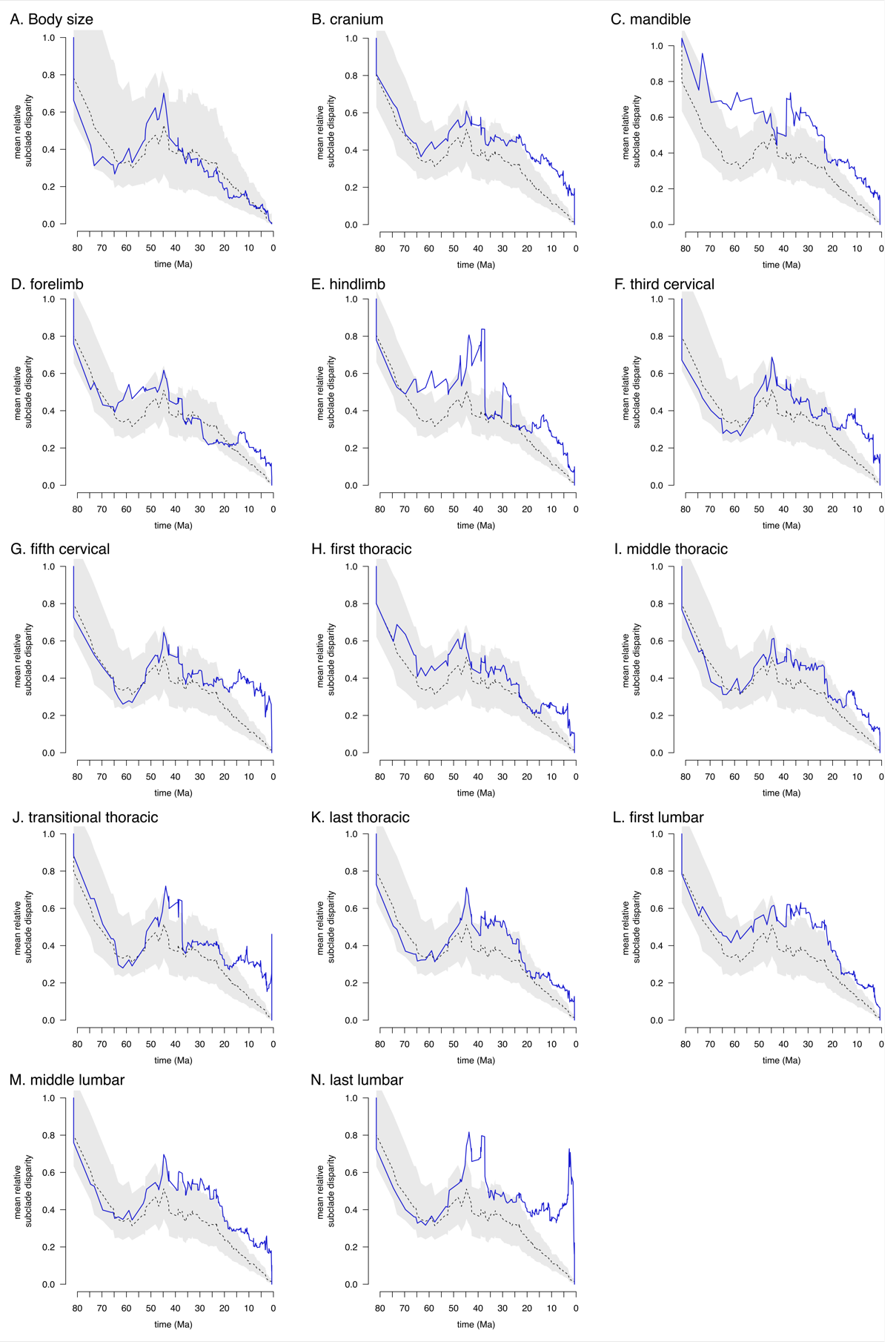

**Fig. S3. Disparity through time plots of skeletal components.** Blue solid line indicates the empirical subclade DTT, and the dashed black line indicates the median DTT based on 10,000 simulations of trait evolution under Brownian motion. The grey shaded area indicates the 95% DTT range for the simulated data.

**Table S1. Specimens and museum catalog numbers used in this study.** AMNH = American Museum of Natural History; CAS = California Academy of Sciences; FMNH = Field Museum of Natural History; JODA = John Day Fossil Beds National Monument; LACM = Natural History Museum of Los Angeles County; MVZ = Museum of Vertebrate Zoology; NHMUK = Natural History Museum, London; SDNHM = San Diego Natural History Museum; TxVP = Texas Vertebrate Paleontology Collection; UCMP = UC Berkeley Museum of Palenotology; UNSM = University of Nebraska State Museum; USNM = National Museum of Natural History; UWBM = Burke Museum of Natural History and Culture; YPM = Yale Peabody Museum

| clade | family | species | status | catalog |
| --- | --- | --- | --- | --- |
| Carnivora | Nandiniidae | *Nandinia_binotata* | extant | AMNH239582 |
| Carnivora | Nandiniidae | *Nandinia_binotata* | extant | FMNH161266 |
| Carnivora | Nandiniidae | *Nandinia_binotata* | extant | FMNH224514 |
| Carnivora | Felidae | *Acinonyx_jubatus* | extant | AMNH119654 |
| Carnivora | Felidae | *Caracal_aurata* | extant | AMNH51994 |
| Carnivora | Felidae | *Caracal_caracal* | extant | USNM520686 |
| Carnivora | Felidae | *Felis_chaus* | extant | AMNH238649 |
| Carnivora | Felidae | *Felis_chaus* | extant | FMNH97867 |
| Carnivora | Felidae | *Felis_silvestris* | extant | NHMUK1953.6.11.1 |
| Carnivora | Felidae | *Herpailurus_yagouaroundi* | extant | AMNH215137 |
| Carnivora | Felidae | *Herpailurus_yagouaroundi* | extant | FMNH69647 |
| Carnivora | Felidae | *Leopardus_geoffroyi* | extant | AMNH205910 |
| Carnivora | Felidae | *Leopardus_geoffroyi* | extant | AMNH205911 |
| Carnivora | Felidae | *Leopardus_pardalis* | extant | AMNH133959 |
| Carnivora | Felidae | *Leptailurus_serval* | extant | FMNH127843 |
| Carnivora | Felidae | *Lynx_canadensis* | extant | MVZ184071 |
| Carnivora | Felidae | *Lynx_canadensis* | extant | UWBM80612 |
| Carnivora | Felidae | *Lynx_canadensis* | extant | UWBM82238 |
| Carnivora | Felidae | *Lynx_rufus* | extant | UWBM32042 |
| Carnivora | Felidae | *Lynx_rufus* | extant | UWBM32046 |
| Carnivora | Felidae | *Lynx_rufus* | extant | UWBM75808 |
| Carnivora | Felidae | *Otocolobus_manul* | extant | UWBM35449 |
| Carnivora | Felidae | *Panthera_leo* | extant | AMNH52078 |
| Carnivora | Felidae | *Panthera_onca* | extant | AMNH139959 |
| Carnivora | Felidae | *Panthera_onca* | extant | FMNH70566 |
| Carnivora | Felidae | *Panthera_pardus* | extant | AMNH113745 |
| Carnivora | Felidae | *Panthera_tigris* | extant | AMNH54460 |
| Carnivora | Felidae | *Prionailurus_bengalensis* | extant | FMNH62888 |
| Carnivora | Felidae | *Prionailurus_bengalensis* | extant | USNM196601 |
| Carnivora | Felidae | *Prionailurus_viverrinus* | extant | AMNH70128 |
| Carnivora | Felidae | *Puma_concolor* | extant | UWBM32105 |
| Carnivora | Felidae | *Puma_concolor* | extant | UWBM35217 |
| Carnivora | Viverridae | *Arctictis_binturong* | extant | FMNH98270 |
| Carnivora | Viverridae | *Arctictis_binturong* | extant | UWBM81977 |
| Carnivora | Viverridae | *Arctogalidia_trivirgata* | extant | NHMUK1971.3072 |
| Carnivora | Viverridae | *Civettictis_civetta* | extant | AMNH51797 |
| Carnivora | Viverridae | *Genetta_genetta* | extant | AMNH187721 |
| Carnivora | Viverridae | *Genetta_genetta* | extant | AMNH187725 |
| Carnivora | Viverridae | *Genetta_genetta* | extant | AMNH187740 |
| Carnivora | Viverridae | *Genetta_maculata* | extant | AMNH216345 |
| Carnivora | Viverridae | *Genetta_maculata* | extant | FMNH17550 |
| Carnivora | Viverridae | *Genetta_servalina* | extant | AMNH150447 |
| Carnivora | Viverridae | *Genetta_servalina* | extant | FMNH145228 |
| Carnivora | Viverridae | *Genetta_servalina* | extant | FMNH145230 |
| Carnivora | Viverridae | *Genetta_victoriae* | extant | AMNH51406 |
| Carnivora | Viverridae | *Paguma_larvata* | extant | UWBM73281 |
| Carnivora | Viverridae | *Paguma_larvata* | extant | NHMUK1980.867 |
| Carnivora | Viverridae | *Paradoxurus_hermaphroditus* | extant | MVZ186573 |
| Carnivora | Viverridae | *Viverra_zibetha* | extant | AMNH113482 |
| Carnivora | Viverridae | *Viverra_zibetha* | extant | FMNH104395 |
| Carnivora | Hyaenidae | *Crocuta_crocuta* | extant | AMNH147880 |
| Carnivora | Hyaenidae | *Crocuta_crocuta* | extant | AMNH187769 |
| Carnivora | Hyaenidae | *Hyaena_hyaena* | extant | USNM329351 |
| Carnivora | Eupleridae | *Cryptoprocta_ferox* | extant | NHMU1938.11.16.1 |
| Carnivora | Eupleridae | *Galidia_elegans* | extant | FMNH151925 |
| Carnivora | Eupleridae | *Galidia_elegans* | extant | FMNH156651 |
| Carnivora | Eupleridae | *Galidictis_fasciata* | extant | FMNH162111 |
| Carnivora | Eupleridae | *Mungotictis_decemlineata* | extant | FMNH176128 |
| Carnivora | Herpestidae | *Atilax_paludinosus* | extant | LACM053753 |
| Carnivora | Herpestidae | *Galerella_pulverulenta* | extant | CAS28740 |
| Carnivora | Herpestidae | *Herpestes_edwardsii* | extant | USNM329348 |
| Carnivora | Herpestidae | *Herpestes_ichneumon* | extant | AMNH187746 |
| Carnivora | Herpestidae | *Herpestes_sanguineus* | extant | AMNH51019 |
| Carnivora | Herpestidae | *Ichneumia_albicauda* | extant | AMNH216353 |
| Carnivora | Herpestidae | *Ichneumia_albicauda* | extant | AMNH51594 |
| Carnivora | Herpestidae | *Ichneumia_albicauda* | extant | TxVP12616 |
| Carnivora | Canidae | *Canis_adustus* | extant | AMNH216344 |
| Carnivora | Canidae | *Canis_aureus* | extant | AMNH54516 |
| Carnivora | Canidae | *Canis_latrans* | extant | MVZ25553 |
| Carnivora | Canidae | *Canis_latrans* | extant | MVZ83446 |
| Carnivora | Canidae | *Canis_latrans* | extant | MVZ83447 |
| Carnivora | Canidae | *Canis_lupus* | extant | MVZ44166 |
| Carnivora | Canidae | *Canis_lupus* | extant | MVZ88225 |
| Carnivora | Canidae | *Canis_mesomelas* | extant | AMNH34734 |
| Carnivora | Canidae | *Canis_simensis* | extant | AMNH81001 |
| Carnivora | Canidae | *Cerdocyon_thous* | extant | AMNH205807 |
| Carnivora | Canidae | *Cerdocyon_thous* | extant | AMNH205811 |
| Carnivora | Canidae | *Chrysocyon_brachyurus* | extant | AMNH133941 |
| Carnivora | Canidae | *Cuon_alpinus* | extant | AMNH54984 |
| Carnivora | Canidae | *Lycaon_pictus* | extant | AMNH82083 |
| Carnivora | Canidae | *Lycaon_pictus* | extant | AMNH82085 |
| Carnivora | Canidae | *Lycaon_pictus* | extant | FMNH127812 |
| Carnivora | Canidae | *Otocyon_megalotis* | extant | LACM041790 |
| Carnivora | Canidae | *Otocyon_megalotis* | extant | LACM041792 |
| Carnivora | Canidae | *Pseudalopex_culpaeus* | extant | AMNH262663 |
| Carnivora | Canidae | *Pseudalopex_gymnocercus* | extant | AMNH205777 |
| Carnivora | Canidae | *Pseudalopex_gymnocercus* | extant | AMNH205778 |
| Carnivora | Canidae | *Speothos_venaticus* | extant | FMNH125402 |
| Carnivora | Canidae | *Urocyon_cinereoargenteus* | extant | MVZ225296 |
| Carnivora | Canidae | *Urocyon_cinereoargenteus* | extant | MVZ236230 |
| Carnivora | Canidae | *Urocyon_cinereoargenteus* | extant | UWBM13640 |
| Carnivora | Canidae | *Urocyon_littoralis* | extant | LACM6391 |
| Carnivora | Canidae | *Urocyon_littoralis* | extant | LACM6397 |
| Carnivora | Canidae | *Urocyon_littoralis* | extant | SDNHM233389 |
| Carnivora | Canidae | *Vulpes_chama* | extant | LACM041795 |
| Carnivora | Canidae | *Vulpes_lagopus* | extant | MVZ206554 |
| Carnivora | Canidae | *Vulpes_lagopus* | extant | UWBM82184 |
| Carnivora | Canidae | *Vulpes_lagopus* | extant | UWBM82638 |
| Carnivora | Canidae | *Vulpes_macrotis* | extant | MVZ206981 |
| Carnivora | Canidae | *Vulpes_macrotis* | extant | MVZ224411 |
| Carnivora | Canidae | *Vulpes_macrotis* | extant | MVZ224412 |
| Carnivora | Canidae | *Vulpes_velox* | extant | LACM86858 |
| Carnivora | Canidae | *Vulpes_velox* | extant | LACM86859 |
| Carnivora | Canidae | *Vulpes_velox* | extant | LACM86862 |
| Carnivora | Canidae | *Vulpes_vulpes* | extant | UWBM32532 |
| Carnivora | Canidae | *Vulpes_vulpes* | extant | UWBM32533 |
| Carnivora | Canidae | *Vulpes_vulpes* | extant | UWBM32534 |
| Carnivora | Canidae | *Vulpes_zerda* | extant | FMNH89716 |
| Carnivora | Ursidae | *Ailuropoda_melanoleuca* | extant | AMNH89028 |
| Carnivora | Ursidae | *Helarctos_malayanus* | extant | USNM198713 |
| Carnivora | Ursidae | *Melursus_ursinus* | extant | AMNH54467 |
| Carnivora | Ursidae | *Ursus_americanus* | extant | UWBM33259 |
| Carnivora | Ursidae | *Ursus_americanus* | extant | UWBM39060 |
| Carnivora | Ursidae | *Ursus_arctos* | extant | MVZ4385 |
| Carnivora | Ursidae | *Ursus_arctos* | extant | MVZ970 |
| Carnivora | Ursidae | *Ursus_maritimus* | extant | AMNH100039 |
| Carnivora | Otariidae | *Arctocephalus_galapagoensis* | extant | AMNH100341 |
| Carnivora | Otariidae | *Arctocephalus_pusillus* | extant | AMNH81701 |
| Carnivora | Otariidae | *Callorhinus_ursinus* | extant | USNM258588 |
| Carnivora | Otariidae | *Eumetopias_jubatus* | extant | AMNH38400 |
| Carnivora | Otariidae | *Otaria_byronia* | extant | USNM484912 |
| Carnivora | Otariidae | *Phocarctos_hookeri* | extant | USNM484526 |
| Carnivora | Otariidae | *Zalophus_californianus* | extant | UWBM32518 |
| Carnivora | Phocidae | *Erignathus_barbatus* | extant | USNM500251 |
| Carnivora | Phocidae | *Monachus_schauinslandi* | extant | USNM606271 |
| Carnivora | Phocidae | *Pusa_hispida* | extant | SDNHM12214 |
| Carnivora | Mephitidae | *Conepatus_leuconotus* | extant | AMNH136415 |
| Carnivora | Mephitidae | *Conepatus_leuconotus* | extant | MVZ85319 |
| Carnivora | Mephitidae | *Conepatus_leuconotus* | extant | TxVP10785 |
| Carnivora | Mephitidae | *Mephitis_macroura* | extant | MVZ41133 |
| Carnivora | Mephitidae | *Mephitis_macroura* | extant | MVZ85324 |
| Carnivora | Mephitidae | *Mephitis_mephitis* | extant | MVZ198469 |
| Carnivora | Mephitidae | *Mephitis_mephitis* | extant | MVZ38300 |
| Carnivora | Mephitidae | *Mephitis_mephitis* | extant | UWBM26340 |
| Carnivora | Mephitidae | *Spilogale_gracilis* | extant | MVZ39890 |
| Carnivora | Mephitidae | *Spilogale_gracilis* | extant | MVZ90009 |
| Carnivora | Mephitidae | *Spilogale_gracilis* | extant | UWBM38625 |
| Carnivora | Mephitidae | *Spilogale_putorius* | extant | MVZ41127 |
| Carnivora | Mephitidae | *Spilogale_putorius* | extant | MVZ41130 |
| Carnivora | Mephitidae | *Spilogale_putorius* | extant | MVZ99980 |
| Carnivora | Ailuridae | *Ailurus_fulgens* | extant | UWBM39029 |
| Carnivora | Ailuridae | *Ailurus_fulgens* | extant | UWBM76122 |
| Carnivora | Procyonidae | *Bassaricyon_neblina* | extant | FMNH70726 |
| Carnivora | Procyonidae | *Bassariscus_astutus* | extant | MVZ191007 |
| Carnivora | Procyonidae | *Bassariscus_astutus* | extant | MVZ44099 |
| Carnivora | Procyonidae | *Bassariscus_astutus* | extant | UWBM34131 |
| Carnivora | Procyonidae | *Nasua_narica* | extant | FMNH129310 |
| Carnivora | Procyonidae | *Nasua_nasua* | extant | AMNH214716 |
| Carnivora | Procyonidae | *Nasua_nasua* | extant | AMNH214718 |
| Carnivora | Procyonidae | *Nasua_nasua* | extant | AMNH255871 |
| Carnivora | Procyonidae | *Potos_flavus* | extant | USNM449468 |
| Carnivora | Procyonidae | *Procyon_lotor* | extant | CAS703 |
| Carnivora | Procyonidae | *Procyon_lotor* | extant | LACM052204 |
| Carnivora | Procyonidae | *Procyon_lotor* | extant | UWBM35531 |
| Carnivora | Mustelidae | *Aonyx_capensis* | extant | MVZ226937 |
| Carnivora | Mustelidae | *Aonyx_congicus* | extant | MVZ19210 |
| Carnivora | Mustelidae | *Eira_barbara* | extant | FMNH69585 |
| Carnivora | Mustelidae | *Eira_barbara* | extant | FMNH69586 |
| Carnivora | Mustelidae | *Eira_barbara* | extant | MVZ153647 |
| Carnivora | Mustelidae | *Enhydra_lutris* | extant | MVZ140630 |
| Carnivora | Mustelidae | *Enhydra_lutris* | extant | MVZ486320 |
| Carnivora | Mustelidae | *Enhydra_lutris* | extant | UWBM34543 |
| Carnivora | Mustelidae | *Galictis_vittata* | extant | USNM395079 |
| Carnivora | Mustelidae | *Gulo_gulo* | extant | FMNH57196 |
| Carnivora | Mustelidae | *Gulo_gulo* | extant | MVZ184099 |
| Carnivora | Mustelidae | *Gulo_gulo* | extant | MVZ22121 |
| Carnivora | Mustelidae | *Lontra_canadensis* | extant | UWBM32230 |
| Carnivora | Mustelidae | *Lontra_canadensis* | extant | UWBM32242 |
| Carnivora | Mustelidae | *Lontra_canadensis* | extant | UWBM81969 |
| Carnivora | Mustelidae | *Lontra_felina* | extant | MVZ141632 |
| Carnivora | Mustelidae | *Lutra_lutra* | extant | FMNH99384 |
| Carnivora | Mustelidae | *Martes_americana* | extant | UWBM33272 |
| Carnivora | Mustelidae | *Martes_americana* | extant | UWBM33278 |
| Carnivora | Mustelidae | *Martes_americana* | extant | UWBM33279 |
| Carnivora | Mustelidae | *Martes_martes* | extant | AMNH183359 |
| Carnivora | Mustelidae | *Meles_meles* | extant | AMNH70604 |
| Carnivora | Mustelidae | *Meles_meles* | extant | NHMUK2002.476 |
| Carnivora | Mustelidae | *Meles_meles* | extant | NHMUK2006.551 |
| Carnivora | Mustelidae | *Mellivora_capensis* | extant | USNM96107 |
| Carnivora | Mustelidae | *Melogale_moschata* | extant | NHMUK2004.20 |
| Carnivora | Mustelidae | *Melogale_personata* | extant | USNM357537 |
| Carnivora | Mustelidae | *Mustela_erminea* | extant | UWBM33273 |
| Carnivora | Mustelidae | *Mustela_erminea* | extant | UWBM56648 |
| Carnivora | Mustelidae | *Mustela_erminea* | extant | UWBM57518 |
| Carnivora | Mustelidae | *Mustela_frenata* | extant | UWBM31872 |
| Carnivora | Mustelidae | *Mustela_frenata* | extant | UWBM39154 |
| Carnivora | Mustelidae | *Mustela_frenata* | extant | UWBM82713 |
| Carnivora | Mustelidae | *Mustela_nigripes* | extant | FMNH88607 |
| Carnivora | Mustelidae | *Mustela_nigripes* | extant | MVZ78134 |
| Carnivora | Mustelidae | *Mustela_nivalis* | extant | MVZ143796 |
| Carnivora | Mustelidae | *Mustela_nivalis* | extant | UWBM30001 |
| Carnivora | Mustelidae | *Mustela_putorius* | extant | LACM31087 |
| Carnivora | Mustelidae | *Mustela_sibirica* | extant | USNM239584 |
| Carnivora | Mustelidae | *Mustela_vison* | extant | UWBM41783 |
| Carnivora | Mustelidae | *Mustela_vison* | extant | UWBM41784 |
| Carnivora | Mustelidae | *Mustela_vison* | extant | UWBM41791 |
| Carnivora | Mustelidae | *Pekania_pennanti* | extant | UWBM34568 |
| Carnivora | Mustelidae | *Pekania_pennanti* | extant | UWBM77855 |
| Carnivora | Mustelidae | *Pekania_pennanti* | extant | UWBM81034 |
| Carnivora | Mustelidae | *Poecilogale_albinucha* | extant | FMNH177235 |
| Carnivora | Mustelidae | *Poecilogale_albinucha* | extant | FMNH177236 |
| Carnivora | Mustelidae | *Pteronura_brasiliensis* | extant | NHMUK1939.3.4.1 |
| Carnivora | Mustelidae | *Pteronura_brasiliensis* | extant | AMNH30190 |
| Carnivora | Mustelidae | *Taxidea_taxus* | extant | MVZ218236 |
| Carnivora | Mustelidae | *Taxidea_taxus* | extant | MVZ41454 |
| Carnivora | Mustelidae | *Taxidea_taxus* | extant | UWBM34281 |
| Carnivora | Felidae | *Amphimachairodus_coloradensis* | extinct | TxVP41261-8 |
| Carnivora | Felidae | *Puma_pardoides* | extinct | YPM031526 |
| Carnivora | Felidae | *Homotherium_serum* | extinct | TxVP_unspec |
| Carnivora | Felidae | *Homotherium_serum* | extinct | TxVP_unspec |
| Carnivora | Felidae | *Homotherium_serum* | extinct | TxVP_unspec |
| Carnivora | Felidae | *Homotherium_serum* | extinct | TxVP_unspec |
| Carnivora | Felidae | *Homotherium_serum* | extinct | TxVP_unspec |
| Carnivora | Felidae | *Homotherium_serum* | extinct | TxVP_unspec |
| Carnivora | Felidae | *Homotherium_serum* | extinct | TxVP_unspec |
| Carnivora | Felidae | *Homotherium_serum* | extinct | TxVP_unspec |
| Carnivora | Felidae | *Homotherium_serum* | extinct | TxVP_unspec |
| Carnivora | Felidae | *Homotherium_serum* | extinct | TxVP_unspec |
| Carnivora | Felidae | *Homotherium_serum* | extinct | TxVP_unspec |
| Carnivora | Felidae | *Homotherium_serum* | extinct | TxVP_unspec |
| Carnivora | Felidae | *Homotherium_serum* | extinct | TxVP_unspec |
| Carnivora | Felidae | *Homotherium_serum* | extinct | TxVP_unspec |
| Carnivora | Felidae | *Homotherium_serum* | extinct | TxVP_unspec |
| Carnivora | Felidae | *Homotherium_serum* | extinct | TxVP_unspec |
| Carnivora | Felidae | *Homotherium_serum* | extinct | TxVP_unspec |
| Carnivora | Felidae | *Homotherium_serum* | extinct | TxVP_unspec |
| Carnivora | Felidae | *Homotherium_serum* | extinct | TxVP_unspec |
| Carnivora | Felidae | *Homotherium_serum* | extinct | TxVP_unspec |
| Carnivora | Felidae | *Homotherium_serum* | extinct | TxVP_unspec |
| Carnivora | Felidae | *Homotherium_serum* | extinct | TxVP_unspec |
| Carnivora | Felidae | *Homotherium_serum* | extinct | TxVP_unspec |
| Carnivora | Felidae | *Homotherium_serum* | extinct | TxVP933-1 |
| Carnivora | Felidae | *Homotherium_serum* | extinct | TxVP933-3444 |
| Carnivora | Felidae | *Homotherium_serum* | extinct | TxVP933-3582 |
| Carnivora | Felidae | *Hyperailurictis_marshi* | extinct | YPM012865 |
| Carnivora | Felidae | *Lynx_longignathus* | extinct | SDNHM68691 |
| Carnivora | Felidae | *Nimravides_catacopsis* | extinct | AMNH104044 |
| Carnivora | Felidae | *Panthera_atrox* | extinct | LACM_unspec |
| Carnivora | Felidae | *Panthera_atrox* | extinct | LACM_unspec |
| Carnivora | Felidae | *Panthera_atrox* | extinct | LACM_unspec |
| Carnivora | Felidae | *Panthera_atrox* | extinct | LACM_unspec |
| Carnivora | Felidae | *Panthera_atrox* | extinct | LACM_unspec |
| Carnivora | Felidae | *Panthera_atrox* | extinct | LACM_unspec |
| Carnivora | Felidae | *Panthera_atrox* | extinct | LACM_unspec |
| Carnivora | Felidae | *Panthera_atrox* | extinct | LACM_unspec |
| Carnivora | Felidae | *Panthera_atrox* | extinct | LACM_unspec |
| Carnivora | Felidae | *Panthera_atrox* | extinct | LACM_unspec |
| Carnivora | Felidae | *Panthera_atrox* | extinct | LACM_unspec |
| Carnivora | Felidae | *Panthera_atrox* | extinct | LACM_unspec |
| Carnivora | Felidae | *Panthera_atrox* | extinct | LACM_unspec |
| Carnivora | Felidae | *Panthera_atrox* | extinct | LACM_unspec |
| Carnivora | Felidae | *Panthera_atrox* | extinct | LACM_unspec |
| Carnivora | Felidae | *Panthera_atrox* | extinct | LACM_unspec |
| Carnivora | Felidae | *Panthera_atrox* | extinct | LACM_unspec |
| Carnivora | Felidae | *Panthera_atrox* | extinct | LACM_unspec |
| Carnivora | Felidae | *Panthera_atrox* | extinct | LACM_unspec |
| Carnivora | Felidae | *Panthera_atrox* | extinct | LACM_unspec |
| Carnivora | Felidae | *Panthera_atrox* | extinct | LACM_unspec |
| Carnivora | Felidae | *Panthera_atrox* | extinct | LACM_unspec |
| Carnivora | Felidae | *Panthera_atrox* | extinct | LACM_unspec |
| Carnivora | Felidae | *Panthera_atrox* | extinct | LACM_unspec |
| Carnivora | Felidae | *Panthera_atrox* | extinct | LACM_unspec |
| Carnivora | Felidae | *Panthera_atrox* | extinct | LACM_unspec |
| Carnivora | Felidae | *Panthera_atrox* | extinct | LACM_unspec |
| Carnivora | Felidae | *Panthera_atrox* | extinct | LACM_unspec |
| Carnivora | Felidae | *Panthera_atrox* | extinct | LACM_unspec |
| Carnivora | Felidae | *Panthera_atrox* | extinct | LACM_unspec |
| Carnivora | Felidae | *Panthera_atrox* | extinct | LACM_unspec |
| Carnivora | Felidae | *Panthera_atrox* | extinct | LACM_unspec |
| Carnivora | Felidae | *Panthera_atrox* | extinct | LACM_unspec |
| Carnivora | Felidae | *Panthera_atrox* | extinct | LACM_unspec |
| Carnivora | Felidae | *Panthera_atrox* | extinct | LACM_unspec |
| Carnivora | Felidae | *Panthera_atrox* | extinct | LACM_unspec |
| Carnivora | Felidae | *Panthera_atrox* | extinct | LACM_unspec |
| Carnivora | Felidae | *Panthera_atrox* | extinct | LACM_unspec |
| Carnivora | Felidae | *Panthera_atrox* | extinct | LACM_unspec |
| Carnivora | Felidae | *Panthera_atrox* | extinct | LACM_unspec |
| Carnivora | Felidae | *Panthera_atrox* | extinct | LACM_unspec |
| Carnivora | Felidae | *Panthera_atrox* | extinct | LACM_unspec |
| Carnivora | Felidae | *Panthera_atrox* | extinct | LACM_unspec |
| Carnivora | Felidae | *Panthera_atrox* | extinct | LACM_unspec |
| Carnivora | Felidae | *Panthera_atrox* | extinct | LACM_unspec |
| Carnivora | Felidae | *Panthera_atrox* | extinct | LACM_unspec |
| Carnivora | Felidae | *Panthera_atrox* | extinct | LACM_unspec |
| Carnivora | Felidae | *Panthera_atrox* | extinct | LACM_unspec |
| Carnivora | Felidae | *Panthera_atrox* | extinct | LACM_unspec |
| Carnivora | Felidae | *Panthera_atrox* | extinct | LACM_unspec |
| Carnivora | Felidae | *Panthera_atrox* | extinct | LACM_unspec |
| Carnivora | Felidae | *Panthera_atrox* | extinct | LACM_unspec |
| Carnivora | Felidae | *Panthera_atrox* | extinct | LACM_unspec |
| Carnivora | Felidae | *Panthera_atrox* | extinct | LACM_unspec |
| Carnivora | Felidae | *Panthera_atrox* | extinct | LACM_unspec |
| Carnivora | Felidae | *Panthera_atrox* | extinct | LACM_unspec |
| Carnivora | Felidae | *Panthera_atrox* | extinct | LACM_unspec |
| Carnivora | Felidae | *Panthera_atrox* | extinct | LACM_unspec |
| Carnivora | Felidae | *Panthera_atrox* | extinct | LACM_unspec |
| Carnivora | Felidae | *Panthera_atrox* | extinct | LACM_unspec |
| Carnivora | Felidae | *Panthera_atrox* | extinct | LACM_unspec |
| Carnivora | Felidae | *Panthera_atrox* | extinct | LACM_unspec |
| Carnivora | Felidae | *Panthera_atrox* | extinct | LACM_unspec |
| Carnivora | Felidae | *Panthera_atrox* | extinct | LACM_unspec |
| Carnivora | Felidae | *Panthera_atrox* | extinct | LACM_unspec |
| Carnivora | Felidae | *Panthera_atrox* | extinct | LACM_unspec |
| Carnivora | Felidae | *Panthera_atrox* | extinct | LACM_unspec |
| Carnivora | Felidae | *Panthera_atrox* | extinct | LACM_unspec |
| Carnivora | Felidae | *Panthera_atrox* | extinct | LACM_unspec |
| Carnivora | Felidae | *Panthera_atrox* | extinct | LACM_unspec |
| Carnivora | Felidae | *Panthera_atrox* | extinct | LACM_unspec |
| Carnivora | Felidae | *Panthera_atrox* | extinct | LACM_unspec |
| Carnivora | Felidae | *Panthera_atrox* | extinct | LACM_unspec |
| Carnivora | Felidae | *Panthera_atrox* | extinct | LACM_unspec |
| Carnivora | Felidae | *Panthera_atrox* | extinct | LACM_unspec |
| Carnivora | Felidae | *Panthera_atrox* | extinct | LACM_unspec |
| Carnivora | Felidae | *Panthera_atrox* | extinct | LACM_unspec |
| Carnivora | Felidae | *Panthera_atrox* | extinct | LACM_unspec |
| Carnivora | Felidae | *Panthera_atrox* | extinct | LACM_unspec |
| Carnivora | Felidae | *Panthera_atrox* | extinct | LACM_unspec |
| Carnivora | Felidae | *Panthera_atrox* | extinct | LACM_unspec |
| Carnivora | Felidae | *Panthera_atrox* | extinct | LACM_unspec |
| Carnivora | Felidae | *Panthera_atrox* | extinct | LACM_unspec |
| Carnivora | Felidae | *Panthera_atrox* | extinct | LACM_unspec |
| Carnivora | Felidae | *Panthera_atrox* | extinct | LACM2900-18 |
| Carnivora | Felidae | *Panthera_atrox* | extinct | LACM2900-8 |
| Carnivora | Felidae | *Panthera_atrox* | extinct | LACM2900-9 |
| Carnivora | Felidae | *Panthera_atrox* | extinct | LACM599 |
| Carnivora | Felidae | *Panthera_atrox* | extinct | LACM65 |
| Carnivora | Felidae | *Panthera_atrox* | extinct | LACMcompile |
| Carnivora | Felidae | *Smilodon_fatalis* | extinct | LACM_unspec |
| Carnivora | Felidae | *Smilodon_fatalis* | extinct | LACM_unspec |
| Carnivora | Felidae | *Smilodon_fatalis* | extinct | LACM_unspec |
| Carnivora | Felidae | *Smilodon_fatalis* | extinct | LACM_unspec |
| Carnivora | Felidae | *Smilodon_fatalis* | extinct | LACM_unspec |
| Carnivora | Felidae | *Smilodon_fatalis* | extinct | LACM_unspec |
| Carnivora | Felidae | *Smilodon_fatalis* | extinct | LACM_unspec |
| Carnivora | Felidae | *Smilodon_fatalis* | extinct | LACM_unspec |
| Carnivora | Felidae | *Smilodon_fatalis* | extinct | LACM_unspec |
| Carnivora | Felidae | *Smilodon_fatalis* | extinct | LACM_unspec |
| Carnivora | Felidae | *Smilodon_fatalis* | extinct | LACM_unspec |
| Carnivora | Felidae | *Smilodon_fatalis* | extinct | LACM_unspec |
| Carnivora | Felidae | *Smilodon_fatalis* | extinct | LACM_unspec |
| Carnivora | Felidae | *Smilodon_fatalis* | extinct | LACM_unspec |
| Carnivora | Felidae | *Smilodon_fatalis* | extinct | LACM_unspec |
| Carnivora | Felidae | *Smilodon_fatalis* | extinct | LACM_unspec |
| Carnivora | Felidae | *Smilodon_fatalis* | extinct | LACM_unspec |
| Carnivora | Felidae | *Smilodon_fatalis* | extinct | LACM_unspec |
| Carnivora | Felidae | *Smilodon_fatalis* | extinct | LACM_unspec |
| Carnivora | Felidae | *Smilodon_fatalis* | extinct | LACM_unspec |
| Carnivora | Felidae | *Smilodon_fatalis* | extinct | LACM_unspec |
| Carnivora | Felidae | *Smilodon_fatalis* | extinct | LACM_unspec |
| Carnivora | Felidae | *Smilodon_fatalis* | extinct | LACM_unspec |
| Carnivora | Felidae | *Smilodon_fatalis* | extinct | LACM_unspec |
| Carnivora | Felidae | *Smilodon_fatalis* | extinct | LACM_unspec |
| Carnivora | Felidae | *Smilodon_fatalis* | extinct | LACM_unspec |
| Carnivora | Felidae | *Smilodon_fatalis* | extinct | LACM_unspec |
| Carnivora | Felidae | *Smilodon_fatalis* | extinct | LACM_unspec |
| Carnivora | Felidae | *Smilodon_fatalis* | extinct | LACM_unspec |
| Carnivora | Felidae | *Smilodon_fatalis* | extinct | LACM_unspec |
| Carnivora | Felidae | *Smilodon_fatalis* | extinct | LACM_unspec |
| Carnivora | Felidae | *Smilodon_fatalis* | extinct | LACM_unspec |
| Carnivora | Felidae | *Smilodon_fatalis* | extinct | LACM_unspec |
| Carnivora | Felidae | *Smilodon_fatalis* | extinct | LACM_unspec |
| Carnivora | Felidae | *Smilodon_fatalis* | extinct | LACM_unspec |
| Carnivora | Felidae | *Smilodon_fatalis* | extinct | LACM_unspec |
| Carnivora | Felidae | *Smilodon_fatalis* | extinct | LACM_unspec |
| Carnivora | Felidae | *Smilodon_fatalis* | extinct | LACM_unspec |
| Carnivora | Felidae | *Smilodon_fatalis* | extinct | LACM_unspec |
| Carnivora | Felidae | *Smilodon_fatalis* | extinct | LACM_unspec |
| Carnivora | Felidae | *Smilodon_fatalis* | extinct | LACM_unspec |
| Carnivora | Felidae | *Smilodon_fatalis* | extinct | LACM_unspec |
| Carnivora | Felidae | *Smilodon_fatalis* | extinct | LACM_unspec |
| Carnivora | Felidae | *Smilodon_fatalis* | extinct | LACM_unspec |
| Carnivora | Felidae | *Smilodon_fatalis* | extinct | LACM_unspec |
| Carnivora | Felidae | *Smilodon_fatalis* | extinct | LACM_unspec |
| Carnivora | Felidae | *Smilodon_fatalis* | extinct | LACM_unspec |
| Carnivora | Felidae | *Smilodon_fatalis* | extinct | LACM_unspec |
| Carnivora | Felidae | *Smilodon_fatalis* | extinct | LACM_unspec |
| Carnivora | Felidae | *Smilodon_fatalis* | extinct | LACM_unspec |
| Carnivora | Felidae | *Smilodon_fatalis* | extinct | LACM_unspec |
| Carnivora | Felidae | *Smilodon_fatalis* | extinct | LACM_unspec |
| Carnivora | Felidae | *Smilodon_fatalis* | extinct | LACM_unspec |
| Carnivora | Felidae | *Smilodon_fatalis* | extinct | LACM_unspec |
| Carnivora | Felidae | *Smilodon_fatalis* | extinct | LACM_unspec |
| Carnivora | Felidae | *Smilodon_fatalis* | extinct | LACM_unspec |
| Carnivora | Felidae | *Smilodon_fatalis* | extinct | LACM_unspec |
| Carnivora | Felidae | *Smilodon_fatalis* | extinct | LACM_unspec |
| Carnivora | Felidae | *Smilodon_fatalis* | extinct | LACM_unspec |
| Carnivora | Felidae | *Smilodon_fatalis* | extinct | LACM_unspec |
| Carnivora | Felidae | *Smilodon_fatalis* | extinct | LACM_unspec |
| Carnivora | Felidae | *Smilodon_fatalis* | extinct | LACM_unspec |
| Carnivora | Felidae | *Smilodon_fatalis* | extinct | LACM_unspec |
| Carnivora | Felidae | *Smilodon_fatalis* | extinct | LACM_unspec |
| Carnivora | Felidae | *Smilodon_fatalis* | extinct | LACM_unspec |
| Carnivora | Felidae | *Smilodon_fatalis* | extinct | LACM_unspec |
| Carnivora | Felidae | *Smilodon_fatalis* | extinct | LACM_unspec |
| Carnivora | Felidae | *Smilodon_fatalis* | extinct | LACM_unspec |
| Carnivora | Felidae | *Smilodon_fatalis* | extinct | LACM_unspec |
| Carnivora | Felidae | *Smilodon_fatalis* | extinct | LACM_unspec |
| Carnivora | Felidae | *Smilodon_fatalis* | extinct | LACM_unspec |
| Carnivora | Felidae | *Smilodon_fatalis* | extinct | LACM_unspec |
| Carnivora | Felidae | *Smilodon_fatalis* | extinct | LACM_unspec |
| Carnivora | Felidae | *Smilodon_fatalis* | extinct | LACM_unspec |
| Carnivora | Felidae | *Smilodon_fatalis* | extinct | LACM_unspec |
| Carnivora | Felidae | *Smilodon_fatalis* | extinct | LACM_unspec |
| Carnivora | Felidae | *Smilodon_fatalis* | extinct | LACM_unspec |
| Carnivora | Felidae | *Smilodon_fatalis* | extinct | LACM_unspec |
| Carnivora | Felidae | *Smilodon_fatalis* | extinct | LACM_unspec |
| Carnivora | Felidae | *Smilodon_fatalis* | extinct | LACM_unspec |
| Carnivora | Felidae | *Smilodon_fatalis* | extinct | LACM_unspec |
| Carnivora | Felidae | *Smilodon_fatalis* | extinct | LACM_unspec |
| Carnivora | Felidae | *Smilodon_fatalis* | extinct | LACM_unspec |
| Carnivora | Felidae | *Smilodon_fatalis* | extinct | LACM_unspec |
| Carnivora | Felidae | *Smilodon_fatalis* | extinct | LACM_unspec |
| Carnivora | Felidae | *Smilodon_fatalis* | extinct | LACM_unspec |
| Carnivora | Felidae | *Smilodon_fatalis* | extinct | LACM_unspec |
| Carnivora | Felidae | *Smilodon_fatalis* | extinct | LACM_unspec |
| Carnivora | Felidae | *Smilodon_fatalis* | extinct | LACM_unspec |
| Carnivora | Felidae | *Smilodon_fatalis* | extinct | LACM_unspec |
| Carnivora | Felidae | *Smilodon_fatalis* | extinct | LACM_unspec |
| Carnivora | Felidae | *Smilodon_fatalis* | extinct | LACM_unspec |
| Carnivora | Felidae | *Smilodon_fatalis* | extinct | LACM_unspec |
| Carnivora | Felidae | *Smilodon_fatalis* | extinct | LACM_unspec |
| Carnivora | Felidae | *Smilodon_fatalis* | extinct | LACM_unspec |
| Carnivora | Felidae | *Smilodon_fatalis* | extinct | LACM_unspec |
| Carnivora | Felidae | *Smilodon_fatalis* | extinct | LACM_unspec |
| Carnivora | Felidae | *Smilodon_fatalis* | extinct | LACM_unspec |
| Carnivora | Felidae | *Smilodon_fatalis* | extinct | LACM_unspec |
| Carnivora | Felidae | *Smilodon_fatalis* | extinct | LACM_unspec |
| Carnivora | Felidae | *Smilodon_fatalis* | extinct | LACM_unspec |
| Carnivora | Felidae | *Smilodon_fatalis* | extinct | LACM_unspec |
| Carnivora | Felidae | *Smilodon_fatalis* | extinct | LACM_unspec |
| Carnivora | Felidae | *Smilodon_fatalis* | extinct | LACM_unspec |
| Carnivora | Felidae | *Smilodon_fatalis* | extinct | LACM_unspec |
| Carnivora | Felidae | *Smilodon_fatalis* | extinct | LACM_unspec |
| Carnivora | Felidae | *Smilodon_fatalis* | extinct | LACM_unspec |
| Carnivora | Felidae | *Smilodon_fatalis* | extinct | LACM_unspec |
| Carnivora | Felidae | *Smilodon_fatalis* | extinct | LACM_unspec |
| Carnivora | Felidae | *Smilodon_fatalis* | extinct | LACM_unspec |
| Carnivora | Felidae | *Smilodon_fatalis* | extinct | LACM_unspec |
| Carnivora | Felidae | *Smilodon_fatalis* | extinct | LACM_unspec |
| Carnivora | Felidae | *Smilodon_fatalis* | extinct | LACM_unspec |
| Carnivora | Felidae | *Smilodon_fatalis* | extinct | LACM_unspec |
| Carnivora | Felidae | *Smilodon_fatalis* | extinct | LACM_unspec |
| Carnivora | Felidae | *Smilodon_fatalis* | extinct | LACM_unspec |
| Carnivora | Felidae | *Smilodon_fatalis* | extinct | LACM_unspec |
| Carnivora | Felidae | *Smilodon_fatalis* | extinct | LACM_unspec |
| Carnivora | Felidae | *Smilodon_fatalis* | extinct | LACM_unspec |
| Carnivora | Felidae | *Smilodon_fatalis* | extinct | LACM_unspec |
| Carnivora | Felidae | *Smilodon_fatalis* | extinct | LACM_unspec |
| Carnivora | Felidae | *Smilodon_fatalis* | extinct | LACM_unspec |
| Carnivora | Felidae | *Smilodon_fatalis* | extinct | LACM_unspec |
| Carnivora | Felidae | *Smilodon_fatalis* | extinct | LACM_unspec |
| Carnivora | Felidae | *Smilodon_fatalis* | extinct | LACM_unspec |
| Carnivora | Felidae | *Smilodon_fatalis* | extinct | LACM_unspec |
| Carnivora | Felidae | *Smilodon_fatalis* | extinct | LACM_unspec |
| Carnivora | Felidae | *Smilodon_fatalis* | extinct | LACM_unspec |
| Carnivora | Felidae | *Smilodon_fatalis* | extinct | LACM_unspec |
| Carnivora | Felidae | *Smilodon_fatalis* | extinct | LACM_unspec |
| Carnivora | Felidae | *Smilodon_fatalis* | extinct | LACM_unspec |
| Carnivora | Felidae | *Smilodon_fatalis* | extinct | LACM_unspec |
| Carnivora | Felidae | *Smilodon_fatalis* | extinct | LACM_unspec |
| Carnivora | Felidae | *Smilodon_fatalis* | extinct | LACM_unspec |
| Carnivora | Felidae | *Smilodon_fatalis* | extinct | LACM_unspec |
| Carnivora | Felidae | *Smilodon_fatalis* | extinct | LACM_unspec |
| Carnivora | Felidae | *Smilodon_fatalis* | extinct | LACM_unspec |
| Carnivora | Felidae | *Smilodon_fatalis* | extinct | LACM_unspec |
| Carnivora | Felidae | *Smilodon_fatalis* | extinct | LACM_unspec |
| Carnivora | Felidae | *Smilodon_fatalis* | extinct | LACM_unspec |
| Carnivora | Felidae | *Smilodon_fatalis* | extinct | LACM_unspec |
| Carnivora | Felidae | *Smilodon_fatalis* | extinct | LACM_unspec |
| Carnivora | Felidae | *Smilodon_fatalis* | extinct | LACM_unspec |
| Carnivora | Felidae | *Smilodon_fatalis* | extinct | LACM_unspec |
| Carnivora | Felidae | *Smilodon_fatalis* | extinct | LACM_unspec |
| Carnivora | Felidae | *Smilodon_fatalis* | extinct | LACM_unspec |
| Carnivora | Felidae | *Smilodon_fatalis* | extinct | LACM_unspec |
| Carnivora | Felidae | *Smilodon_fatalis* | extinct | LACM_unspec |
| Carnivora | Felidae | *Smilodon_fatalis* | extinct | LACM_unspec |
| Carnivora | Felidae | *Smilodon_fatalis* | extinct | LACM_unspec |
| Carnivora | Felidae | *Smilodon_fatalis* | extinct | LACM1326 |
| Carnivora | Felidae | *Smilodon_fatalis* | extinct | LACM1339 |
| Carnivora | Felidae | *Smilodon_fatalis* | extinct | LACM1341 |
| Carnivora | Felidae | *Smilodon_fatalis* | extinct | LACM1359 |
| Carnivora | Felidae | *Smilodon_fatalis* | extinct | LACM186 |
| Carnivora | Felidae | *Smilodon_gracilis* | extinct | UF226962 |
| Carnivora | Felidae | *Smilodon_gracilis* | extinct | UF226963 |
| Carnivora | Felidae | *Smilodon_gracilis* | extinct | UF226991 |
| Carnivora | Felidae | *Smilodon_gracilis* | extinct | UF226994 |
| Carnivora | Felidae | *Smilodon_gracilis* | extinct | UF80088 |
| Carnivora | Felidae | *Smilodon_gracilis* | extinct | UF80162 |
| Carnivora | Felidae | *Smilodon_gracilis* | extinct | UF81154 |
| Carnivora | Felidae | *Smilodon_gracilis* | extinct | UF81294 |
| Carnivora | Felidae | *Smilodon_gracilis* | extinct | UF81724 |
| Carnivora | Felidae | *Smilodon_gracilis* | extinct | UF84411 |
| Carnivora | Felidae | *Smilodon_gracilis* | extinct | UF86330 |
| Carnivora | Felidae | *Smilodon_gracilis* | extinct | UF87239 |
| Carnivora | Felidae | *Smilodon_gracilis* | extinct | UF_unspecified |
| Carnivora | Felidae | *Smilodon_populator* | extinct | FMNH-P14271 |
| Carnivora | Felidae | *Smilodon_populator* | extinct | FMNH-P14294 |
| Carnivora | Canidae | *Aelurodon_ferox* | extinct | AMNH_unspec |
| Carnivora | Canidae | *Aelurodon_ferox* | extinct | UNSM46630 |
| Carnivora | Canidae | *Aelurodon_ferox* | extinct | UNSM76624 |
| Carnivora | Canidae | *Aelurodon_ferox* | extinct | UNSM76625 |
| Carnivora | Canidae | *Aelurodon_ferox* | extinct | UNSM76631 |
| Carnivora | Canidae | *Aelurodon_ferox* | extinct | UNSM83898 |
| Carnivora | Canidae | *Aelurodon_ferox* | extinct | UNSM83910 |
| Carnivora | Canidae | *Aelurodon_ferox* | extinct | UNSM83916 |
| Carnivora | Canidae | *Aelurodon_ferox* | extinct | UNSM83917 |
| Carnivora | Canidae | *Aelurodon_ferox* | extinct | UNSM83923 |
| Carnivora | Canidae | *Aelurodon_ferox* | extinct | UNSM83929 |
| Carnivora | Canidae | *Aelurodon_ferox* | extinct | USNM539 |
| Carnivora | Canidae | *Aelurodon_ferox* | extinct | YPM010057 |
| Carnivora | Canidae | *Aelurodon_mcgrewi* | extinct | AMNH22410 |
| Carnivora | Canidae | *Aelurodon_stirtoni* | extinct | UNSM25789 |
| Carnivora | Canidae | *Aenocyon_dirus* | extinct | LACM_unspec |
| Carnivora | Canidae | *Aenocyon_dirus* | extinct | LACM_unspec |
| Carnivora | Canidae | *Aenocyon_dirus* | extinct | LACM_unspec |
| Carnivora | Canidae | *Aenocyon_dirus* | extinct | LACM_unspec |
| Carnivora | Canidae | *Aenocyon_dirus* | extinct | LACM_unspec |
| Carnivora | Canidae | *Aenocyon_dirus* | extinct | LACM_unspec |
| Carnivora | Canidae | *Aenocyon_dirus* | extinct | LACM_unspec |
| Carnivora | Canidae | *Aenocyon_dirus* | extinct | LACM_unspec |
| Carnivora | Canidae | *Aenocyon_dirus* | extinct | LACM_unspec |
| Carnivora | Canidae | *Aenocyon_dirus* | extinct | LACM_unspec |
| Carnivora | Canidae | *Aenocyon_dirus* | extinct | LACM_unspec |
| Carnivora | Canidae | *Aenocyon_dirus* | extinct | LACM_unspec |
| Carnivora | Canidae | *Aenocyon_dirus* | extinct | LACM_unspec |
| Carnivora | Canidae | *Aenocyon_dirus* | extinct | LACM_unspec |
| Carnivora | Canidae | *Aenocyon_dirus* | extinct | LACM_unspec |
| Carnivora | Canidae | *Aenocyon_dirus* | extinct | LACM_unspec |
| Carnivora | Canidae | *Aenocyon_dirus* | extinct | LACM_unspec |
| Carnivora | Canidae | *Aenocyon_dirus* | extinct | LACM_unspec |
| Carnivora | Canidae | *Aenocyon_dirus* | extinct | LACM_unspec |
| Carnivora | Canidae | *Aenocyon_dirus* | extinct | LACM_unspec |
| Carnivora | Canidae | *Aenocyon_dirus* | extinct | LACM_unspec |
| Carnivora | Canidae | *Aenocyon_dirus* | extinct | LACM_unspec |
| Carnivora | Canidae | *Aenocyon_dirus* | extinct | LACM_unspec |
| Carnivora | Canidae | *Aenocyon_dirus* | extinct | LACM_unspec |
| Carnivora | Canidae | *Aenocyon_dirus* | extinct | LACM_unspec |
| Carnivora | Canidae | *Aenocyon_dirus* | extinct | LACM_unspec |
| Carnivora | Canidae | *Aenocyon_dirus* | extinct | LACM_unspec |
| Carnivora | Canidae | *Aenocyon_dirus* | extinct | LACM_unspec |
| Carnivora | Canidae | *Aenocyon_dirus* | extinct | LACM_unspec |
| Carnivora | Canidae | *Aenocyon_dirus* | extinct | LACM_unspec |
| Carnivora | Canidae | *Aenocyon_dirus* | extinct | LACM_unspec |
| Carnivora | Canidae | *Aenocyon_dirus* | extinct | LACM_unspec |
| Carnivora | Canidae | *Aenocyon_dirus* | extinct | LACM_unspec |
| Carnivora | Canidae | *Aenocyon_dirus* | extinct | LACM_unspec |
| Carnivora | Canidae | *Aenocyon_dirus* | extinct | LACM_unspec |
| Carnivora | Canidae | *Aenocyon_dirus* | extinct | LACM_unspec |
| Carnivora | Canidae | *Aenocyon_dirus* | extinct | LACM_unspec |
| Carnivora | Canidae | *Aenocyon_dirus* | extinct | LACM_unspec |
| Carnivora | Canidae | *Aenocyon_dirus* | extinct | LACM_unspec |
| Carnivora | Canidae | *Aenocyon_dirus* | extinct | LACM_unspec |
| Carnivora | Canidae | *Aenocyon_dirus* | extinct | LACM_unspec |
| Carnivora | Canidae | *Aenocyon_dirus* | extinct | LACM_unspec |
| Carnivora | Canidae | *Aenocyon_dirus* | extinct | LACM_unspec |
| Carnivora | Canidae | *Aenocyon_dirus* | extinct | LACM_unspec |
| Carnivora | Canidae | *Aenocyon_dirus* | extinct | LACM_unspec |
| Carnivora | Canidae | *Aenocyon_dirus* | extinct | LACM_unspec |
| Carnivora | Canidae | *Aenocyon_dirus* | extinct | LACM_unspec |
| Carnivora | Canidae | *Aenocyon_dirus* | extinct | LACM_unspec |
| Carnivora | Canidae | *Aenocyon_dirus* | extinct | LACM_unspec |
| Carnivora | Canidae | *Aenocyon_dirus* | extinct | LACM_unspec |
| Carnivora | Canidae | *Aenocyon_dirus* | extinct | LACM_unspec |
| Carnivora | Canidae | *Aenocyon_dirus* | extinct | LACM_unspec |
| Carnivora | Canidae | *Aenocyon_dirus* | extinct | LACM_unspec |
| Carnivora | Canidae | *Aenocyon_dirus* | extinct | LACM_unspec |
| Carnivora | Canidae | *Aenocyon_dirus* | extinct | LACM_unspec |
| Carnivora | Canidae | *Aenocyon_dirus* | extinct | LACM_unspec |
| Carnivora | Canidae | *Aenocyon_dirus* | extinct | LACM_unspec |
| Carnivora | Canidae | *Aenocyon_dirus* | extinct | LACM_unspec |
| Carnivora | Canidae | *Aenocyon_dirus* | extinct | LACM_unspec |
| Carnivora | Canidae | *Aenocyon_dirus* | extinct | LACM_unspec |
| Carnivora | Canidae | *Aenocyon_dirus* | extinct | LACM_unspec |
| Carnivora | Canidae | *Aenocyon_dirus* | extinct | LACM_unspec |
| Carnivora | Canidae | *Aenocyon_dirus* | extinct | LACM_unspec |
| Carnivora | Canidae | *Aenocyon_dirus* | extinct | LACM_unspec |
| Carnivora | Canidae | *Aenocyon_dirus* | extinct | LACM_unspec |
| Carnivora | Canidae | *Aenocyon_dirus* | extinct | LACM_unspec |
| Carnivora | Canidae | *Aenocyon_dirus* | extinct | LACM_unspec |
| Carnivora | Canidae | *Aenocyon_dirus* | extinct | LACM_unspec |
| Carnivora | Canidae | *Aenocyon_dirus* | extinct | LACM_unspec |
| Carnivora | Canidae | *Aenocyon_dirus* | extinct | LACM_unspec |
| Carnivora | Canidae | *Aenocyon_dirus* | extinct | LACM_unspec |
| Carnivora | Canidae | *Aenocyon_dirus* | extinct | LACM_unspec |
| Carnivora | Canidae | *Aenocyon_dirus* | extinct | LACM_unspec |
| Carnivora | Canidae | *Aenocyon_dirus* | extinct | LACM_unspec |
| Carnivora | Canidae | *Aenocyon_dirus* | extinct | LACM_unspec |
| Carnivora | Canidae | *Aenocyon_dirus* | extinct | LACM_unspec |
| Carnivora | Canidae | *Aenocyon_dirus* | extinct | LACM_unspec |
| Carnivora | Canidae | *Aenocyon_dirus* | extinct | LACM_unspec |
| Carnivora | Canidae | *Aenocyon_dirus* | extinct | LACM_unspec |
| Carnivora | Canidae | *Aenocyon_dirus* | extinct | LACM_unspec |
| Carnivora | Canidae | *Aenocyon_dirus* | extinct | LACM_unspec |
| Carnivora | Canidae | *Aenocyon_dirus* | extinct | LACM_unspec |
| Carnivora | Canidae | *Aenocyon_dirus* | extinct | LACM_unspec |
| Carnivora | Canidae | *Aenocyon_dirus* | extinct | LACM_unspec |
| Carnivora | Canidae | *Aenocyon_dirus* | extinct | LACM_unspec |
| Carnivora | Canidae | *Aenocyon_dirus* | extinct | LACM_unspec |
| Carnivora | Canidae | *Aenocyon_dirus* | extinct | LACM_unspec |
| Carnivora | Canidae | *Aenocyon_dirus* | extinct | LACM_unspec |
| Carnivora | Canidae | *Aenocyon_dirus* | extinct | LACM_unspec |
| Carnivora | Canidae | *Aenocyon_dirus* | extinct | LACM_unspec |
| Carnivora | Canidae | *Aenocyon_dirus* | extinct | LACM_unspec |
| Carnivora | Canidae | *Aenocyon_dirus* | extinct | LACM_unspec |
| Carnivora | Canidae | *Aenocyon_dirus* | extinct | LACM_unspec |
| Carnivora | Canidae | *Aenocyon_dirus* | extinct | LACM_unspec |
| Carnivora | Canidae | *Aenocyon_dirus* | extinct | LACM_unspec |
| Carnivora | Canidae | *Aenocyon_dirus* | extinct | LACM_unspec |
| Carnivora | Canidae | *Aenocyon_dirus* | extinct | LACM_unspec |
| Carnivora | Canidae | *Aenocyon_dirus* | extinct | LACM_unspec |
| Carnivora | Canidae | *Aenocyon_dirus* | extinct | LACM_unspec |
| Carnivora | Canidae | *Aenocyon_dirus* | extinct | LACM_unspec |
| Carnivora | Canidae | *Aenocyon_dirus* | extinct | LACM_unspec |
| Carnivora | Canidae | *Aenocyon_dirus* | extinct | LACM_unspec |
| Carnivora | Canidae | *Aenocyon_dirus* | extinct | LACM_unspec |
| Carnivora | Canidae | *Aenocyon_dirus* | extinct | LACM_unspec |
| Carnivora | Canidae | *Aenocyon_dirus* | extinct | LACM_unspec |
| Carnivora | Canidae | *Aenocyon_dirus* | extinct | LACM_unspec |
| Carnivora | Canidae | *Aenocyon_dirus* | extinct | LACM_unspec |
| Carnivora | Canidae | *Aenocyon_dirus* | extinct | LACM_unspec |
| Carnivora | Canidae | *Aenocyon_dirus* | extinct | LACM_unspec |
| Carnivora | Canidae | *Aenocyon_dirus* | extinct | LACM_unspec |
| Carnivora | Canidae | *Aenocyon_dirus* | extinct | LACM_unspec |
| Carnivora | Canidae | *Aenocyon_dirus* | extinct | LACM_unspec |
| Carnivora | Canidae | *Aenocyon_dirus* | extinct | LACM_unspec |
| Carnivora | Canidae | *Aenocyon_dirus* | extinct | LACM_unspec |
| Carnivora | Canidae | *Aenocyon_dirus* | extinct | LACM_unspec |
| Carnivora | Canidae | *Aenocyon_dirus* | extinct | LACM_unspec |
| Carnivora | Canidae | *Aenocyon_dirus* | extinct | LACM_unspec |
| Carnivora | Canidae | *Aenocyon_dirus* | extinct | LACM_unspec |
| Carnivora | Canidae | *Aenocyon_dirus* | extinct | LACM_unspec |
| Carnivora | Canidae | *Aenocyon_dirus* | extinct | LACM_unspec |
| Carnivora | Canidae | *Aenocyon_dirus* | extinct | LACM_unspec |
| Carnivora | Canidae | *Aenocyon_dirus* | extinct | LACM_unspec |
| Carnivora | Canidae | *Aenocyon_dirus* | extinct | LACM_unspec |
| Carnivora | Canidae | *Aenocyon_dirus* | extinct | LACM_unspec |
| Carnivora | Canidae | *Aenocyon_dirus* | extinct | LACM_unspec |
| Carnivora | Canidae | *Aenocyon_dirus* | extinct | LACM_unspec |
| Carnivora | Canidae | *Aenocyon_dirus* | extinct | LACM_unspec |
| Carnivora | Canidae | *Aenocyon_dirus* | extinct | LACM_unspec |
| Carnivora | Canidae | *Aenocyon_dirus* | extinct | LACM_unspec |
| Carnivora | Canidae | *Aenocyon_dirus* | extinct | LACM_unspec |
| Carnivora | Canidae | *Aenocyon_dirus* | extinct | LACM_unspec |
| Carnivora | Canidae | *Aenocyon_dirus* | extinct | LACM_unspec |
| Carnivora | Canidae | *Aenocyon_dirus* | extinct | LACM_unspec |
| Carnivora | Canidae | *Aenocyon_dirus* | extinct | LACM_unspec |
| Carnivora | Canidae | *Aenocyon_dirus* | extinct | LACM_unspec |
| Carnivora | Canidae | *Aenocyon_dirus* | extinct | LACM_unspec |
| Carnivora | Canidae | *Aenocyon_dirus* | extinct | LACM_unspec |
| Carnivora | Canidae | *Aenocyon_dirus* | extinct | LACM_unspec |
| Carnivora | Canidae | *Aenocyon_dirus* | extinct | LACM_unspec |
| Carnivora | Canidae | *Aenocyon_dirus* | extinct | LACM_unspec |
| Carnivora | Canidae | *Aenocyon_dirus* | extinct | LACM_unspec |
| Carnivora | Canidae | *Aenocyon_dirus* | extinct | LACM_unspec |
| Carnivora | Canidae | *Aenocyon_dirus* | extinct | LACM_unspec |
| Carnivora | Canidae | *Aenocyon_dirus* | extinct | LACM_unspec |
| Carnivora | Canidae | *Aenocyon_dirus* | extinct | LACMcompile |
| Carnivora | Canidae | *Canis_edwardii* | extinct | AMNH63100 |
| Carnivora | Canidae | *Enhydrocyon_stenocephalus* | extinct | JODA6222 |
| Carnivora | Canidae | *Epicyon_haydeni* | extinct | UCMP29638 |
| Carnivora | Canidae | *Epicyon_saevus* | extinct | AMNH8305 |
| Carnivora | Canidae | *Eucyon_davisi* | extinct | UO3241 |
| Carnivora | Canidae | *Hesperocyon_gregarius* | extinct | FMNH-UC1405 |
| Carnivora | Canidae | *Hesperocyon_gregarius* | extinct | FMNH-UC1417 |
| Carnivora | Canidae | *Hesperocyon_gregarius* | extinct | FMNH-UC495 |
| Carnivora | Canidae | *Hesperocyon_gregarius* | extinct | UNSM133290 |
| Carnivora | Canidae | *Hesperocyon_gregarius* | extinct | UNSM25701 |
| Carnivora | Canidae | *Hesperocyon_gregarius* | extinct | YPM010068 |
| Carnivora | Canidae | *Leptocyon_vafer* | extinct | UNSM51949 |
| Carnivora | Canidae | *Leptocyon_vafer* | extinct | UNSM51950 |
| Carnivora | Canidae | *Mesocyon_brachyops* | extinct | UO4351 |
| Carnivora | Canidae | *Mesocyon_coryphaeus* | extinct | USNM552 |
| Carnivora | Canidae | *Mesocyon_coryphaeus* | extinct | YPM011163 |
| Carnivora | Canidae | *Mesocyon_coryphaeus* | extinct |  |
| Carnivora | Canidae | *Paraenhydrocyon_josephi* | extinct | JODA761 |
| Carnivora | Canidae | *Paraenhydrocyon_josephi* | extinct | YPM012702 |
| Carnivora | Canidae | *Phlaocyon_multicuspus* | extinct |  |
| Carnivora | Canidae | *Protocyon_troglodytes* | extinct | UF27889 |
| Carnivora | Canidae | *Sunkahetanka_geringensis* | extinct | YPM013602 |
| Carnivora | Canidae | *Tephrocyon_rurestris* | extinct | UO24191 |
| Carnivora | Ursidae | *Arctodus_pristinus* | extinct | UF322971 |
| Carnivora | Ursidae | *Arctodus_pristinus* | extinct | UF40089 |
| Carnivora | Ursidae | *Arctodus_simus* | extinct | USNM521336 |
| Carnivora | Ursidae | *Arctodus_simus* | extinct | YPM055748 |
| Carnivora | Ursidae | *Plithocyon_ursinus* | extinct | AMNH21101 |
| Carnivora | Ursidae | *Plithocyon_ursinus* | extinct | YPM012915 |
| Carnivora | Ursidae | *Tremarctos_floridanus* | extinct | UF133299 |
| Carnivora | 12Phocidae | *Pteronarctos_goedertae* | extinct | UO35536 |
| Carnivora | 12Phocidae | *Pteronarctos_piersoni* | extinct | UO35535 |
| Carnivora | 14Ailuridae | *Simocyon_primigenius* | extinct | YPM011649 |
| Carnivora | Procyonidae | *Cyonasua_brevirostris* | extinct | FMNH-P14342 |
| Carnivora | Procyonidae | *Cyonasua_brevirostris* | extinct | FMNH-P14537 |
| Carnivora | Mustelidae | *Enhydrictis_ardea* | extinct | YPM031529 |
| Carnivora | Mustelidae | *Enhydritherium_terraenovae* | extinct | UF100000 |
| Carnivora | Mustelidae | *Satherium_ingens* | extinct | UO11837 |
| Carnivora | Mustelidae | *Megalictis_ferox* | extinct | FMNH12154 |
| Carnivora | Mustelidae | *Promartes_olcotti* | extinct | FMNH14055 |
| Carnivora | Mustelidae | *Promartes_olcotti* | extinct | FMNH15178 |
| Carnivora | Mustelidae | *Satherium_piscinarium* | extinct | USNM23266 |
| Carnivora | Mustelidae | *Sthenictis_dolichops* | extinct | AMNH25235 |
| Carnivora | Mustelidae | *Taxidea_mexicana* | extinct | LACM_CIT3532 |
| Carnivora | Mustelidae | *Trigonictis_cookii* | extinct | UO16352 |
| Carnivora | Mustelidae | *Trigonictis_macrodon* | extinct | UF234500 |
| Carnivora | Mustelidae | *Trigonictis_macrodon* | extinct | UO16351 |
| Carnivora | Mustelidae | *Zodiolestes_daimonelixensis* | extinct | FMNH12032 |
| Carnivora | Amphicyonidae | *Amphicyon_ingens* | extinct | AMNH |
| Carnivora | Amphicyonidae | *Crassidia_intermedia* | extinct | UF_unspec |
| Carnivora | Amphicyonidae | *Crassidia_intermedia* | extinct | UF_unspec |
| Carnivora | Amphicyonidae | *Crassidia_intermedia* | extinct | UF_unspec |
| Carnivora | Amphicyonidae | *Crassidia_intermedia* | extinct | UF_unspec |
| Carnivora | Amphicyonidae | *Crassidia_intermedia* | extinct | UF_unspec |
| Carnivora | Amphicyonidae | *Crassidia_intermedia* | extinct | UF_unspec |
| Carnivora | Amphicyonidae | *Crassidia_intermedia* | extinct | UF_unspec |
| Carnivora | Amphicyonidae | *Crassidia_intermedia* | extinct | UF_unspec |
| Carnivora | Amphicyonidae | *Crassidia_intermedia* | extinct | UF_unspec |
| Carnivora | Amphicyonidae | *Crassidia_intermedia* | extinct | UF_unspec |
| Carnivora | Amphicyonidae | *Crassidia_intermedia* | extinct | UF_unspec |
| Carnivora | Amphicyonidae | *Crassidia_intermedia* | extinct | UF_unspec |
| Carnivora | Amphicyonidae | *Crassidia_intermedia* | extinct | UF162716 |
| Carnivora | Amphicyonidae | *Crassidia_intermedia* | extinct | UF201000 |
| Carnivora | Amphicyonidae | *Crassidia_intermedia* | extinct | UF211501 |
| Carnivora | Amphicyonidae | *Crassidia_intermedia* | extinct | UF254770 |
| Carnivora | Amphicyonidae | *Crassidia_intermedia* | extinct | UF260014 |
| Carnivora | Amphicyonidae | *Crassidia_intermedia* | extinct | UF299539 |
| Carnivora | Amphicyonidae | *Crassidia_intermedia* | extinct | UF6501 |
| Carnivora | Amphicyonidae | *Brachyrhynchocyon_dodgei* | extinct | USNM17847 |
| Carnivora | Amphicyonidae | *Brachyrhynchocyon_dodgei* | extinct | YPM024612 |
| Carnivora | Amphicyonidae | *Cynelos_sinapius* | extinct | AMNH9356 |
| Carnivora | Amphicyonidae | *Daphoenus_hartshornianus* | extinct | UF207948 |
| Carnivora | Amphicyonidae | *Daphoenus_hartshornianus* | extinct | UNSM25785 |
| Carnivora | Amphicyonidae | *Daphoenus_hartshornianus* | extinct | YPM011421 |
| Carnivora | Amphicyonidae | *Daphoenus_vetus* | extinct | FMNH-P12021 |
| Carnivora | Amphicyonidae | *Daphoenus_vetus* | extinct | YPM013580 |
| Carnivora | Amphicyonidae | *Daphoenus_vetus* | extinct | YPM013792 |
| Carnivora | Amphicyonidae | *Daphoenus_vetus* | extinct | USNM17842 |
| Carnivora | Amphicyonidae | *Temnocyon_altigenis* | extinct | YPM010065 |
| Carnivora | Amphicyonidae | *Temnocyon_fingeruti* | extinct | UO280161 |
| Carnivora | Barbourofelidae | *Barbourofelis_loveorum* | extinct | UF_unspec |
| Carnivora | Barbourofelidae | *Barbourofelis_loveorum* | extinct | UF_unspec |
| Carnivora | Barbourofelidae | *Barbourofelis_loveorum* | extinct | UF_unspec |
| Carnivora | Barbourofelidae | *Barbourofelis_loveorum* | extinct | UF_unspec |
| Carnivora | Barbourofelidae | *Barbourofelis_loveorum* | extinct | UF_unspec |
| Carnivora | Barbourofelidae | *Barbourofelis_loveorum* | extinct | UF_unspec |
| Carnivora | Barbourofelidae | *Barbourofelis_loveorum* | extinct | UF_unspec |
| Carnivora | Barbourofelidae | *Barbourofelis_loveorum* | extinct | UF_unspec |
| Carnivora | Barbourofelidae | *Barbourofelis_loveorum* | extinct | UF_unspec |
| Carnivora | Barbourofelidae | *Barbourofelis_loveorum* | extinct | UF_unspec |
| Carnivora | Barbourofelidae | *Barbourofelis_loveorum* | extinct | UF_unspec |
| Carnivora | Barbourofelidae | *Barbourofelis_loveorum* | extinct | UF_unspec |
| Carnivora | Barbourofelidae | *Barbourofelis_loveorum* | extinct | UF_unspec |
| Carnivora | Barbourofelidae | *Barbourofelis_loveorum* | extinct | UF_unspec |
| Carnivora | Barbourofelidae | *Barbourofelis_loveorum* | extinct | UF_unspec |
| Carnivora | Barbourofelidae | *Barbourofelis_loveorum* | extinct | UF_unspec |
| Carnivora | Barbourofelidae | *Barbourofelis_loveorum* | extinct | UF_unspec |
| Carnivora | Barbourofelidae | *Barbourofelis_loveorum* | extinct | UF_unspec |
| Carnivora | Barbourofelidae | *Barbourofelis_loveorum* | extinct | UF_unspec |
| Carnivora | Barbourofelidae | *Barbourofelis_loveorum* | extinct | UF_unspec |
| Carnivora | Barbourofelidae | *Barbourofelis_loveorum* | extinct | UF_unspec |
| Carnivora | Barbourofelidae | *Barbourofelis_loveorum* | extinct | UF_unspec |
| Carnivora | Barbourofelidae | *Barbourofelis_loveorum* | extinct | UF_unspec |
| Carnivora | Barbourofelidae | *Barbourofelis_loveorum* | extinct | UF_unspec |
| Carnivora | Barbourofelidae | *Barbourofelis_loveorum* | extinct | UF_unspec |
| Carnivora | Barbourofelidae | *Barbourofelis_loveorum* | extinct | UF_unspec |
| Carnivora | Barbourofelidae | *Barbourofelis_loveorum* | extinct | UF_unspec |
| Carnivora | Barbourofelidae | *Barbourofelis_loveorum* | extinct | UF_unspec |
| Carnivora | Barbourofelidae | *Barbourofelis_loveorum* | extinct | UF_unspec |
| Carnivora | Barbourofelidae | *Barbourofelis_loveorum* | extinct | UF_unspec |
| Carnivora | Barbourofelidae | *Barbourofelis_loveorum* | extinct | UF_unspec |
| Carnivora | Barbourofelidae | *Barbourofelis_loveorum* | extinct | UF_unspec |
| Carnivora | Barbourofelidae | *Barbourofelis_loveorum* | extinct | UF_unspec |
| Carnivora | Barbourofelidae | *Barbourofelis_loveorum* | extinct | UF_unspec |
| Carnivora | Barbourofelidae | *Barbourofelis_loveorum* | extinct | UF24429 |
| Carnivora | Barbourofelidae | *Barbourofelis_loveorum* | extinct | UF24447 |
| Carnivora | Barbourofelidae | *Barbourofelis_loveorum* | extinct | UF24458 |
| Carnivora | Barbourofelidae | *Barbourofelis_loveorum* | extinct | UF30000 |
| Carnivora | Barbourofelidae | *Barbourofelis_loveorum* | extinct | UF37000 |
| Carnivora | Barbourofelidae | *Barbourofelis_loveorum* | extinct | UF466168 |
| Carnivora | Barbourofelidae | *Barbourofelis_loveorum* | extinct | UF466174 |
| Carnivora | Desmatophocidae | *Allodesmus_kelloggi* | extinct | AMNH32763 |
| Carnivora | Desmatophocidae | *Allodesmus_kernensis* | extinct | SDNHM143027 |
| Carnivora | Nimravidae | *Dinictis_felina* | extinct | AMNH38805 |
| Carnivora | Nimravidae | *Dinictis_felina* | extinct | UF207947 |
| Carnivora | Nimravidae | *Dinictis_felina* | extinct | USNM18219 |
| Carnivora | Nimravidae | *Dinictis_felina* | extinct | UWBM90773 |
| Carnivora | Nimravidae | *Dinictis_felina* | extinct | YPM010035 |
| Carnivora | Nimravidae | *Dinictis_felina* | extinct | YPM010972 |
| Carnivora | Nimravidae | *Dinictis_felina* | extinct | YPM056858 |
| Carnivora | Nimravidae | *Hoplophoneus_cerebralis* | extinct | JODA7047 |
| Carnivora | Nimravidae | *Hoplophoneus_mentalis* | extinct | YPM013593 |
| Carnivora | Nimravidae | *Hoplophoneus_primaevus* | extinct | AMNH655 |
| Carnivora | Nimravidae | *Hoplophoneus_primaevus* | extinct | UWBM87094 |
| Carnivora | Nimravidae | *Hoplophoneus_sicarius* | extinct | FMNH-PM9467 |
| Carnivora | Nimravidae | *Nimravus_brachyops* | extinct | JODA1312 |
| Carnivora | Nimravidae | *Nimravus_brachyops* | extinct | JODA17778 |
| Carnivora | Nimravidae | *Nimravus_brachyops* | extinct | USNM3957 |
| Carnivora | Nimravidae | *Nimravus_brachyops* | extinct | YPM010517 |
| Carnivora | Semantoridae | *Potamotherium_valletoni* | extinct | AMNH_unspec |
| Carnivora | Semantoridae | *Potamotherium_valletoni* | extinct | AMNH_unspec |
| Carnivora | Semantoridae | *Potamotherium_valletoni* | extinct | AMNH_unspec |
| Carnivora | Semantoridae | *Potamotherium_valletoni* | extinct | AMNH_unspec |
| Carnivora | Semantoridae | *Potamotherium_valletoni* | extinct | AMNH_unspec |
| Carnivora | Semantoridae | *Potamotherium_valletoni* | extinct | AMNH_unspec |
| Carnivora | Semantoridae | *Potamotherium_valletoni* | extinct | AMNH_unspec |
| Carnivora | Semantoridae | *Potamotherium_valletoni* | extinct | AMNH_unspec |
| Carnivora | Semantoridae | *Potamotherium_valletoni* | extinct | AMNH_unspec |
| Carnivora | Semantoridae | *Potamotherium_valletoni* | extinct | AMNH_unspec |
| Carnivora | Semantoridae | *Potamotherium_valletoni* | extinct | AMNH_unspec |
| Carnivora | Semantoridae | *Potamotherium_valletoni* | extinct | AMNH_unspec |
| Carnivora | Semantoridae | *Potamotherium_valletoni* | extinct | AMNH_unspec |
| Carnivora | Semantoridae | *Potamotherium_valletoni* | extinct | AMNH_unspec |
| Carnivora | Semantoridae | *Potamotherium_valletoni* | extinct | AMNH_unspec |
| Carnivora | Semantoridae | *Potamotherium_valletoni* | extinct | AMNH_unspec |
| Carnivora | Semantoridae | *Potamotherium_valletoni* | extinct | AMNH_unspec |
| Carnivora | Semantoridae | *Potamotherium_valletoni* | extinct | AMNH_unspec |
| Carnivora | Semantoridae | *Potamotherium_valletoni* | extinct | AMNH_unspec |
| Carnivora | Semantoridae | *Potamotherium_valletoni* | extinct | AMNH_unspec |
| Carnivora | Semantoridae | *Potamotherium_valletoni* | extinct | AMNH_unspec |
| Carnivora | Semantoridae | *Potamotherium_valletoni* | extinct | AMNH_unspec |
| Carnivora | Semantoridae | *Potamotherium_valletoni* | extinct | AMNH_unspec |
| Carnivora | Semantoridae | *Potamotherium_valletoni* | extinct | AMNH_unspec |
| Carnivora | Semantoridae | *Potamotherium_valletoni* | extinct | AMNH_unspec |
| Carnivora | Semantoridae | *Potamotherium_valletoni* | extinct | AMNH_unspec |
| Carnivora | Semantoridae | *Potamotherium_valletoni* | extinct | AMNH_unspec |
| Carnivora | Semantoridae | *Potamotherium_valletoni* | extinct | AMNH_unspec |
| Carnivora | Semantoridae | *Potamotherium_valletoni* | extinct | AMNH_unspec |
| Carnivora | Semantoridae | *Potamotherium_valletoni* | extinct | AMNH_unspec |
| Carnivora | Semantoridae | *Potamotherium_valletoni* | extinct | AMNH_unspec |
| Carnivora | Semantoridae | *Potamotherium_valletoni* | extinct | AMNH_unspec |
| Carnivora | Semantoridae | *Potamotherium_valletoni* | extinct | AMNH22520 |
| Carnivora | Semantoridae | *Potamotherium_valletoni* | extinct | AMNHcomp |
| Carnivora | Subparictidae | *Eoarctos_vorax* | extinct | USNM637259 |
| Carnivoramorpha | Miacidae | *Miacis_washakius* | extinct | FMNH-PM3869 |
| Carnivoramorpha | Miacidae | *Miocyon_vallisrubrae* | extinct | TxVP_unspec |
| Carnivoramorpha | Miacidae | *Miocyon_vallisrubrae* | extinct | TxVP40165-4 |
| Carnivoramorpha | Miacidae | *Neovulpavus_mccarrolli* | extinct | FMNH3593 |
| Carnivoramorpha | Miacidae | *Tapocyon_robustus* | extinct | SDNHM36000 |
| Carnivoramorpha | Miacidae | *Vulpavus_hargeri* | extinct | YPM011839 |
| Carnivoramorpha |  | *Vulpavus_profectus* | extinct | USNM538 |
| Pan-Carnivora | Hyaenodontidae | *Arfia_shoshoniensis* | extinct | YPM016141 |
| Pan-Carnivora | Hyaenodontidae | *Hyaenodon_crucians* | extinct | YPM010076 |
| Pan-Carnivora | Hyaenodontidae | *Hyaenodon_crucians* | extinct | YPM012553 |
| Pan-Carnivora | Hyaenodontidae | *Hyaenodon_horridus* | extinct | UNSM1-15-8-33SP |
| Pan-Carnivora | Hyaenodontidae | *Hyaenodon_horridus* | extinct | YPM010010 |
| Pan-Carnivora | Hyaenodontidae | *Hyaenodon_horridus* | extinct | YPM010074 |
| Pan-Carnivora | Hyaenodontidae | *Limnocyon_potens* | extinct | AMNH13138 |
| Pan-Carnivora | Hyaenodontidae | *Limnocyon_verus* | extinct | AMNH12155 |
| Pan-Carnivora | Hyaenodontidae | *Limnocyon_verus* | extinct | YPM011796 |
| Pan-Carnivora | Hyaenodontidae | *Sinopa_major* | extinct | USNM5341 |
| Pan-Carnivora | Hyaenodontidae | *Tritemnodon_agilis* | extinct | YPM010073 |
| Pan-Carnivora | Oxyaenidae | *Oxyaena_forcipata* | extinct | YPM014551 |
| Pan-Carnivora | Oxyaenidae | *Palaeonictis_occidentalis* | extinct | FMNH-216 |

**Table S2. Trait loadings from PCAs of skeletal phenome and each component**. Trait abbreviations are defined in Fig. S1. All traits are size-corrected.

| A. Skeletal phenome | | PC1 | PC2 | PC3 | PC4 | PC5 | PC6 | PC7 |
| --- | --- | --- | --- | --- | --- | --- | --- | --- |
|  | Proportion of Variance | 30% | 18% | 11% | 6% | 5% | 4% | 3% |
|  | lnCBL | 0.06 | 0.05 | 0.02 | 0.13 | 0.08 | 0.05 | 0.11 |
|  | lnPOC | 0.20 | -0.12 | -0.12 | 0.13 | -0.24 | 0.20 | -0.09 |
|  | lnZB | 0.04 | -0.02 | -0.04 | 0.07 | 0.01 | 0.11 | 0.01 |
|  | lnMB | -0.01 | 0.12 | -0.21 | 0.13 | -0.07 | 0.07 | -0.03 |
|  | lnCVH | 0.05 | 0.04 | -0.02 | 0.10 | -0.01 | 0.10 | 0.01 |
|  | lnPW | 0.13 | -0.06 | -0.04 | 0.04 | 0.04 | 0.01 | -0.07 |
|  | lnBCL | 0.08 | 0.09 | 0.01 | 0.02 | 0.01 | 0.05 | 0.03 |
|  | lnMAT | 0.05 | 0.04 | -0.04 | 0.14 | 0.15 | 0.15 | 0.07 |
|  | lnMAM | 0.01 | 0.00 | 0.06 | 0.02 | 0.21 | 0.00 | 0.39 |
|  | lnMAM2 | 0.01 | 0.05 | 0.06 | 0.03 | 0.11 | 0.19 | 0.09 |
|  | lnMOL | 0.02 | 0.05 | 0.07 | 0.13 | 0.15 | 0.09 | 0.28 |
|  | lnCOL | 0.07 | 0.03 | 0.11 | 0.12 | 0.14 | 0.05 | 0.21 |
|  | lnMW | 0.07 | 0.05 | -0.06 | 0.02 | 0.01 | 0.02 | 0.00 |
|  | lnAMW | 0.07 | -0.04 | -0.06 | 0.02 | 0.02 | 0.06 | -0.04 |
|  | lnscap_L | -0.05 | -0.06 | 0.14 | -0.05 | -0.03 | 0.06 | -0.03 |
|  | lnscap_W | -0.17 | 0.03 | 0.13 | -0.06 | -0.12 | 0.17 | -0.02 |
|  | lnhum_L | 0.09 | -0.14 | 0.08 | -0.02 | -0.01 | 0.15 | 0.00 |
|  | lnhum_D | -0.16 | -0.02 | 0.13 | -0.04 | -0.02 | 0.19 | -0.02 |
|  | lnhum_prox | -0.10 | 0.00 | 0.05 | -0.01 | -0.07 | 0.15 | -0.06 |
|  | lnhum_dist | -0.13 | -0.02 | -0.05 | 0.01 | 0.00 | 0.21 | -0.09 |
|  | lnul_L | 0.06 | -0.13 | 0.18 | -0.04 | -0.04 | 0.13 | 0.03 |
|  | lnul_D | -0.22 | -0.04 | -0.05 | -0.13 | 0.06 | 0.29 | -0.21 |
|  | lnul_OL | -0.14 | 0.05 | 0.11 | -0.06 | 0.10 | 0.21 | -0.09 |
|  | lnrad_L | 0.07 | -0.15 | 0.24 | -0.04 | -0.07 | 0.13 | 0.08 |
|  | lnrad_D | -0.24 | -0.01 | 0.27 | -0.11 | -0.10 | 0.22 | 0.06 |
|  | lnMC3L | 0.18 | -0.10 | 0.26 | -0.05 | -0.17 | -0.03 | 0.02 |
|  | lnMC3W | -0.07 | -0.12 | 0.11 | -0.05 | -0.01 | 0.02 | -0.17 |
|  | lnpel_L | -0.02 | -0.06 | -0.02 | -0.10 | -0.01 | -0.04 | 0.05 |
|  | lnfem_L | 0.17 | -0.27 | 0.05 | 0.00 | -0.03 | 0.14 | -0.01 |
|  | lnfem_D | -0.13 | -0.10 | 0.02 | -0.23 | 0.57 | -0.12 | 0.03 |
|  | lnfem_EB | -0.06 | -0.05 | -0.02 | -0.06 | -0.03 | 0.04 | 0.05 |
|  | lnfem_GT | -0.08 | -0.21 | -0.07 | 0.09 | 0.12 | 0.21 | 0.37 |
|  | lntib_L | 0.13 | -0.04 | 0.10 | -0.13 | -0.06 | 0.02 | 0.15 |
|  | lntib_D | -0.04 | -0.09 | 0.13 | -0.14 | -0.08 | 0.01 | 0.07 |
|  | lntib_prox | -0.04 | -0.08 | 0.00 | -0.07 | -0.06 | 0.06 | 0.01 |
|  | lntib_dist | -0.05 | -0.07 | -0.01 | -0.05 | -0.10 | 0.07 | -0.08 |
|  | lnfib_L | 0.14 | -0.05 | 0.10 | -0.13 | -0.04 | 0.01 | 0.15 |
|  | lncal_L | 0.03 | -0.09 | 0.08 | -0.10 | -0.01 | 0.02 | 0.03 |
|  | lnMT3L | 0.26 | -0.07 | 0.17 | -0.18 | -0.10 | -0.18 | 0.02 |
|  | lnMT3D | 0.03 | -0.14 | 0.07 | -0.18 | -0.07 | -0.12 | -0.14 |
|  | lnC3_VW | -0.06 | 0.11 | -0.03 | 0.07 | -0.04 | -0.01 | -0.11 |
|  | lnC3_VH | 0.01 | 0.15 | 0.09 | 0.06 | -0.02 | -0.01 | -0.05 |
|  | lnC3_CL | 0.03 | 0.17 | 0.26 | 0.02 | 0.01 | -0.07 | 0.03 |
|  | lnC3_CW | -0.08 | 0.03 | 0.00 | 0.01 | -0.08 | 0.05 | -0.04 |
|  | lnC3_CH | -0.07 | 0.16 | 0.11 | 0.06 | -0.05 | -0.10 | 0.03 |
|  | lnC3_ZL | 0.04 | 0.16 | 0.21 | 0.06 | 0.02 | -0.07 | 0.00 |
|  | lnC3_NSH | 0.07 | 0.12 | 0.07 | 0.08 | 0.00 | 0.07 | -0.10 |
|  | lnC5_VW | -0.11 | 0.05 | -0.08 | 0.04 | -0.02 | 0.07 | -0.12 |
|  | lnC5_VH | 0.00 | 0.11 | 0.14 | 0.07 | 0.03 | -0.04 | -0.11 |
|  | lnC5_CL | -0.02 | 0.22 | 0.19 | 0.05 | 0.05 | -0.06 | 0.04 |
|  | lnC5_CW | -0.10 | 0.05 | -0.05 | 0.06 | -0.04 | 0.04 | 0.00 |
|  | lnC5_CH | -0.07 | 0.14 | 0.18 | 0.04 | -0.07 | -0.10 | 0.06 |
|  | lnC5_ZL | 0.00 | 0.19 | 0.16 | 0.08 | -0.01 | -0.02 | 0.01 |
|  | lnC5_NSH | 0.03 | 0.08 | 0.11 | 0.07 | 0.16 | 0.00 | -0.23 |
|  | lnT1_VW | -0.10 | 0.06 | -0.08 | 0.01 | 0.00 | 0.03 | 0.03 |
|  | lnT1_VH | -0.02 | 0.00 | 0.16 | 0.02 | 0.03 | -0.04 | -0.11 |
|  | lnT1_CL | -0.02 | 0.19 | 0.08 | -0.04 | -0.02 | 0.00 | -0.02 |
|  | lnT1_CW | -0.08 | -0.01 | 0.02 | 0.12 | -0.07 | 0.00 | -0.01 |
|  | lnT1_CH | -0.18 | 0.07 | 0.11 | 0.03 | -0.02 | -0.06 | 0.04 |
|  | lnT1_ZL | -0.03 | 0.20 | 0.09 | -0.02 | -0.02 | 0.01 | -0.02 |
|  | lnT1_NSH | 0.01 | -0.03 | 0.18 | 0.03 | 0.06 | -0.03 | -0.16 |
|  | lnmidT_VW | -0.07 | 0.04 | -0.08 | -0.01 | -0.06 | -0.02 | 0.01 |
|  | lnmidT_VH | -0.07 | -0.10 | 0.10 | -0.03 | 0.02 | -0.04 | -0.12 |
|  | lnmidT_CL | -0.01 | 0.18 | -0.03 | -0.10 | -0.01 | 0.01 | -0.03 |
|  | lnmidT_CW | -0.09 | 0.04 | -0.01 | -0.04 | -0.13 | -0.03 | 0.03 |
|  | lnmidT_CH | -0.18 | 0.00 | 0.09 | -0.02 | -0.05 | 0.00 | 0.04 |
|  | lnmidT_ZL | -0.02 | 0.13 | -0.06 | -0.08 | 0.00 | -0.01 | -0.01 |
|  | lnmidT_NSH | -0.03 | -0.15 | 0.11 | -0.03 | 0.15 | -0.06 | -0.21 |
|  | lntranT_VW | -0.05 | 0.03 | -0.08 | 0.02 | -0.13 | -0.04 | 0.05 |
|  | lntranT_VH | -0.13 | -0.07 | 0.01 | 0.10 | 0.16 | -0.10 | -0.05 |
|  | lntranT_CL | 0.00 | 0.16 | -0.04 | -0.17 | -0.04 | 0.00 | 0.02 |
|  | lntranT_CW | -0.08 | 0.03 | -0.02 | 0.06 | -0.12 | -0.06 | 0.04 |
|  | lntranT_CH | -0.19 | -0.04 | 0.02 | -0.06 | -0.01 | -0.01 | 0.06 |
|  | lntranT_ZL | -0.06 | 0.12 | -0.06 | -0.05 | -0.03 | -0.02 | 0.05 |
|  | lntranT_NSH | -0.09 | -0.08 | 0.01 | 0.18 | 0.22 | -0.15 | -0.10 |
|  | lnlastT_VW | -0.06 | 0.01 | -0.10 | 0.03 | -0.04 | -0.03 | 0.02 |
|  | lnlastT_VH | -0.14 | -0.08 | 0.02 | 0.02 | -0.03 | -0.08 | 0.04 |
|  | lnlastT_CL | 0.04 | 0.14 | -0.05 | -0.18 | -0.05 | -0.01 | 0.07 |
|  | lnlastT_CW | -0.08 | -0.01 | -0.06 | 0.02 | -0.08 | -0.03 | 0.02 |
|  | lnlastT_CH | -0.18 | -0.01 | -0.03 | -0.07 | -0.06 | 0.03 | 0.10 |
|  | lnlastT_ZL | 0.01 | 0.08 | -0.06 | -0.14 | -0.02 | -0.02 | 0.04 |
|  | lnlastT_NSH | -0.11 | -0.12 | 0.04 | 0.07 | -0.02 | -0.14 | 0.01 |
|  | lnL1_VW | -0.09 | -0.02 | 0.01 | 0.06 | -0.12 | -0.21 | 0.15 |
|  | lnL1_VH | -0.13 | -0.08 | 0.01 | 0.03 | -0.07 | -0.12 | 0.04 |
|  | lnL1_CL | 0.06 | 0.11 | -0.05 | -0.21 | -0.05 | -0.03 | 0.06 |
|  | lnL1_CW | -0.07 | -0.04 | -0.08 | 0.03 | -0.09 | -0.02 | 0.02 |
|  | lnL1_CH | -0.18 | 0.00 | -0.03 | -0.09 | -0.02 | 0.01 | 0.10 |
|  | lnL1_ZL | 0.03 | 0.08 | -0.06 | -0.17 | -0.02 | -0.01 | 0.02 |
|  | lnL1_NSH | -0.10 | -0.12 | 0.03 | 0.11 | -0.13 | -0.18 | 0.01 |
|  | lnmidL_VW | -0.06 | -0.08 | 0.03 | 0.08 | -0.16 | -0.20 | 0.09 |
|  | lnmidL_VH | -0.08 | -0.10 | 0.01 | -0.03 | -0.04 | -0.13 | 0.02 |
|  | lnmidL_CL | 0.09 | 0.09 | -0.03 | -0.27 | 0.00 | -0.01 | 0.05 |
|  | lnmidL_CW | -0.05 | -0.04 | -0.08 | 0.02 | -0.09 | -0.01 | 0.03 |
|  | lnmidL_CH | -0.19 | -0.03 | -0.02 | -0.08 | -0.08 | 0.01 | 0.11 |
|  | lnmidL_ZL | 0.07 | 0.08 | -0.06 | -0.21 | -0.01 | 0.03 | 0.01 |
|  | lnmidL_NSH | -0.03 | -0.15 | 0.02 | -0.01 | 0.01 | -0.19 | -0.04 |
|  | lnlastL_VW | 0.00 | -0.09 | -0.02 | 0.08 | -0.08 | -0.11 | 0.11 |
|  | lnlastL_VH | -0.09 | -0.07 | -0.06 | -0.09 | 0.03 | -0.09 | -0.01 |
|  | lnlastL_CL | 0.05 | 0.07 | -0.09 | -0.27 | 0.06 | -0.01 | -0.01 |
|  | lnlastL_CW | -0.03 | -0.06 | -0.07 | 0.02 | -0.07 | -0.04 | 0.06 |
|  | lnlastL_CH | -0.16 | 0.00 | -0.03 | -0.14 | -0.02 | -0.01 | 0.11 |
|  | lnlastL_ZL | 0.04 | 0.07 | -0.11 | -0.21 | 0.08 | 0.01 | -0.02 |
|  | lnlastL_NSH | -0.05 | -0.12 | -0.08 | -0.10 | 0.14 | -0.14 | -0.09 |
| B. Cranium | | PC1 | PC2 | PC3 | PC4 | PC5 | PC6 | PC7 |
|  | Proportion of Variance | 50% | 20% | 13% | 8% | 5% | 2% | 2% |
|  | lnCBL | 0.13 | 0.14 | 0.48 | 0.18 | 0.30 | 0.73 | 0.27 |
|  | lnPOC | 0.86 | -0.17 | -0.39 | 0.27 | 0.02 | 0.05 | 0.04 |
|  | lnZB | 0.17 | 0.08 | 0.10 | -0.25 | 0.21 | 0.20 | -0.90 |
|  | lnMB | 0.16 | 0.87 | -0.21 | -0.29 | -0.27 | 0.08 | 0.12 |
|  | lnCVH | 0.18 | 0.25 | 0.23 | -0.07 | 0.72 | -0.56 | 0.14 |
|  | lnPW | 0.35 | -0.30 | 0.42 | -0.70 | -0.27 | -0.07 | 0.20 |
|  | lnBCL | 0.18 | 0.20 | 0.58 | 0.50 | -0.45 | -0.32 | -0.20 |
| C. Mandible | | PC1 | PC2 | PC3 | PC4 | PC5 | PC6 | PC7 |
|  | Proportion of Variance | 47% | 21% | 14% | 8% | 6% | 3% | 3% |
|  | lnMAT | 0.41 | 0.47 | 0.34 | 0.59 | 0.36 | 0.16 | 0.06 |
|  | lnMAM | 0.53 | -0.81 | 0.13 | 0.19 | 0.04 | -0.05 | -0.01 |
|  | lnMAM2 | 0.33 | 0.21 | -0.28 | 0.29 | -0.77 | -0.07 | -0.30 |
|  | lnMOL | 0.47 | 0.20 | -0.31 | -0.31 | 0.01 | -0.24 | 0.70 |
|  | lnCOL | 0.45 | 0.15 | -0.08 | -0.57 | 0.23 | 0.31 | -0.54 |
|  | lnMW | 0.07 | 0.13 | 0.34 | -0.13 | 0.09 | -0.87 | -0.29 |
|  | lnAMW | 0.06 | 0.05 | 0.76 | -0.31 | -0.46 | 0.24 | 0.23 |
| D. Forelimb | | PC1 | PC2 | PC3 | PC4 | PC5 | PC6 | PC7 |
|  | Proportion of Variance | 49% | 27% | 7% | 5% | 3% | 3% | 2% |
|  | lnscap_L | 0.13 | 0.20 | 0.01 | 0.08 | 0.07 | 0.05 | 0.21 |
|  | lnscap_W | 0.35 | 0.08 | 0.20 | 0.15 | -0.30 | -0.58 | 0.51 |
|  | lnhum_L | -0.10 | 0.29 | -0.32 | 0.31 | -0.11 | 0.08 | 0.00 |
|  | lnhum_D | 0.33 | 0.12 | 0.04 | -0.01 | -0.10 | 0.17 | -0.05 |
|  | lnhum_prox | 0.22 | 0.05 | 0.04 | -0.07 | 0.01 | 0.14 | -0.46 |
|  | lnhum_dist | 0.24 | -0.08 | -0.26 | 0.11 | -0.09 | 0.02 | -0.19 |
|  | lnul_L | -0.04 | 0.40 | -0.09 | 0.30 | 0.08 | 0.14 | 0.02 |
|  | lnul_D | 0.44 | -0.12 | -0.58 | 0.14 | -0.06 | -0.31 | -0.28 |
|  | lnul_OL | 0.31 | 0.00 | 0.13 | 0.16 | 0.89 | -0.10 | 0.07 |
|  | lnrad_L | -0.06 | 0.50 | -0.03 | 0.37 | -0.04 | 0.19 | 0.09 |
|  | lnrad_D | 0.51 | 0.28 | 0.44 | -0.21 | -0.21 | 0.24 | -0.18 |
|  | lnMC3L | -0.25 | 0.55 | 0.02 | -0.36 | 0.10 | -0.58 | -0.36 |
|  | lnMC3W | 0.15 | 0.21 | -0.48 | -0.64 | 0.10 | 0.23 | 0.44 |
| E. Hindlimb | | PC1 | PC2 | PC3 | PC4 | PC5 | PC6 | PC7 |
|  | Proportion of Variance | 37% | 24% | 14% | 7% | 5% | 4% | 3% |
|  | lnpel_L | 0.03 | 0.14 | 0.04 | 0.64 | 0.70 | 0.23 | 0.10 |
|  | lnfem_L | 0.48 | 0.35 | -0.25 | -0.02 | 0.19 | -0.55 | -0.45 |
|  | lnfem_D | -0.26 | 0.51 | 0.76 | -0.23 | 0.13 | -0.08 | -0.13 |
|  | lnfem_EB | -0.06 | 0.15 | -0.03 | 0.19 | -0.11 | 0.08 | -0.05 |
|  | lnfem_GT | -0.10 | 0.66 | -0.50 | -0.33 | 0.05 | 0.30 | 0.32 |
|  | lntib_L | 0.33 | 0.01 | 0.08 | -0.08 | -0.04 | 0.39 | -0.26 |
|  | lntib_D | 0.07 | 0.19 | 0.08 | 0.32 | -0.47 | 0.31 | -0.09 |
|  | lntib_prox | 0.00 | 0.16 | -0.03 | 0.24 | -0.16 | 0.02 | -0.10 |
|  | lntib_dist | -0.02 | 0.12 | -0.08 | 0.27 | -0.25 | -0.10 | -0.06 |
|  | lnfib_L | 0.34 | 0.01 | 0.10 | -0.13 | 0.02 | 0.38 | -0.24 |
|  | lncal_L | 0.16 | 0.14 | 0.07 | 0.16 | -0.22 | 0.09 | -0.18 |
|  | lnMT3L | 0.63 | -0.07 | 0.24 | -0.18 | 0.07 | 0.07 | 0.48 |
|  | lnMT3D | 0.19 | 0.21 | 0.16 | 0.28 | -0.28 | -0.35 | 0.51 |
| F. Third cervical | | PC1 | PC2 | PC3 | PC4 | PC5 | PC6 | PC7 |
|  | Proportion of Variance | 62% | 17% | 12% | 5% | 3% | 1% | 0% |
|  | lnC3_VW | 0.16 | 0.54 | 0.43 | 0.50 | 0.43 | 0.23 | 0.02 |
|  | lnC3_VH | 0.37 | 0.01 | 0.33 | -0.31 | -0.01 | -0.12 | -0.80 |
|  | lnC3_CL | 0.57 | -0.17 | -0.38 | 0.20 | -0.21 | 0.65 | -0.07 |
|  | lnC3_CW | 0.05 | 0.47 | 0.06 | 0.24 | -0.82 | -0.23 | 0.00 |
|  | lnC3_CH | 0.39 | 0.52 | -0.19 | -0.63 | 0.13 | -0.05 | 0.35 |
|  | lnC3_ZL | 0.51 | -0.19 | -0.24 | 0.38 | 0.21 | -0.68 | 0.08 |
|  | lnC3_NSH | 0.32 | -0.39 | 0.68 | -0.12 | -0.20 | 0.03 | 0.46 |
| G. Fifth cervical | | PC1 | PC2 | PC3 | PC4 | PC5 | PC6 | PC7 |
|  | Proportion of Variance | 61% | 19% | 11% | 5% | 2% | 2% | 1% |
|  | lnC5_VW | 0.06 | 0.61 | 0.49 | 0.03 | 0.50 | 0.35 | 0.11 |
|  | lnC5_VH | 0.34 | -0.19 | 0.35 | -0.32 | 0.12 | -0.10 | -0.77 |
|  | lnC5_CL | 0.59 | 0.00 | -0.23 | 0.46 | -0.23 | 0.56 | -0.14 |
|  | lnC5_CW | 0.09 | 0.58 | 0.19 | 0.22 | -0.57 | -0.47 | -0.13 |
|  | lnC5_CH | 0.44 | 0.26 | -0.28 | -0.74 | -0.14 | 0.04 | 0.29 |
|  | lnC5_ZL | 0.49 | -0.04 | -0.19 | 0.29 | 0.52 | -0.58 | 0.18 |
|  | lnC5_NSH | 0.28 | -0.43 | 0.66 | 0.01 | -0.25 | -0.02 | 0.48 |
| H. First thoracic | | PC1 | PC2 | PC3 | PC4 | PC5 | PC6 | PC7 |
|  | Proportion of Variance | 50% | 23% | 19% | 3% | 3% | 1% | 0% |
|  | lnT1_VW | 0.28 | 0.43 | 0.02 | 0.57 | 0.64 | 0.03 | 0.01 |
|  | lnT1_VH | 0.19 | -0.48 | 0.29 | 0.10 | 0.12 | 0.07 | 0.78 |
|  | lnT1_CL | 0.43 | -0.17 | -0.51 | -0.11 | 0.08 | -0.71 | 0.05 |
|  | lnT1_CW | 0.24 | 0.22 | 0.35 | -0.77 | 0.41 | 0.00 | -0.02 |
|  | lnT1_CH | 0.66 | 0.19 | 0.41 | 0.15 | -0.56 | -0.04 | -0.13 |
|  | lnT1_ZL | 0.45 | -0.19 | -0.51 | -0.12 | 0.03 | 0.69 | -0.08 |
|  | lnT1_NSH | 0.07 | -0.66 | 0.32 | 0.15 | 0.28 | -0.05 | -0.60 |
| I. Middle thoracic | | PC1 | PC2 | PC3 | PC4 | PC5 | PC6 | PC7 |
|  | Proportion of Variance | 43% | 37% | 11% | 5% | 3% | 1% | 1% |
|  | lnmidT_VW | 0.02 | 0.35 | 0.31 | 0.67 | 0.57 | 0.05 | 0.04 |
|  | lnmidT_VH | -0.51 | 0.11 | -0.22 | 0.07 | -0.07 | 0.06 | 0.82 |
|  | lnmidT_CL | 0.40 | 0.32 | -0.59 | 0.06 | -0.02 | 0.62 | -0.01 |
|  | lnmidT_CW | -0.03 | 0.45 | 0.29 | 0.30 | -0.78 | 0.01 | -0.09 |
|  | lnmidT_CH | -0.38 | 0.67 | 0.04 | -0.54 | 0.24 | -0.01 | -0.24 |
|  | lnmidT_ZL | 0.29 | 0.28 | -0.47 | 0.10 | 0.01 | -0.78 | 0.06 |
|  | lnmidT_NSH | -0.60 | -0.17 | -0.45 | 0.39 | -0.04 | 0.00 | -0.51 |
| J. Diaphragmatic | | PC1 | PC2 | PC3 | PC4 | PC5 | PC6 | PC7 |
|  | Proportion of Variance | 51% | 29% | 10% | 7% | 2% | 2% | 0% |
|  | lntranT_VW | 0.06 | 0.30 | 0.00 | 0.65 | 0.56 | 0.41 | 0.08 |
|  | lntranT_VH | 0.58 | -0.06 | 0.13 | -0.13 | 0.00 | 0.01 | 0.79 |
|  | lntranT_CL | -0.20 | 0.48 | 0.53 | -0.29 | 0.42 | -0.44 | 0.06 |
|  | lntranT_CW | 0.17 | 0.31 | -0.03 | 0.59 | -0.44 | -0.58 | 0.00 |
|  | lntranT_CH | 0.49 | 0.46 | -0.57 | -0.33 | 0.18 | -0.06 | -0.28 |
|  | lntranT_ZL | 0.03 | 0.53 | 0.34 | -0.13 | -0.53 | 0.55 | -0.07 |
|  | lntranT_NSH | 0.59 | -0.31 | 0.51 | 0.07 | 0.09 | -0.02 | -0.53 |
| K. Last thoracic | | PC1 | PC2 | PC3 | PC4 | PC5 | PC6 | PC7 |
|  | Proportion of Variance | 60% | 23% | 8% | 5% | 2% | 1% | 0% |
|  | lnlastT_VW | 0.18 | 0.27 | 0.83 | 0.19 | 0.41 | 0.02 | 0.00 |
|  | lnlastT_VH | 0.51 | 0.01 | -0.21 | 0.20 | 0.10 | 0.05 | -0.80 |
|  | lnlastT_CL | -0.28 | 0.62 | -0.22 | 0.30 | -0.02 | 0.63 | 0.00 |
|  | lnlastT_CW | 0.27 | 0.21 | 0.31 | 0.01 | -0.89 | -0.02 | -0.01 |
|  | lnlastT_CH | 0.54 | 0.46 | -0.22 | -0.58 | 0.18 | -0.01 | 0.28 |
|  | lnlastT_ZL | -0.14 | 0.47 | -0.20 | 0.35 | 0.02 | -0.77 | 0.01 |
|  | lnlastT_NSH | 0.50 | -0.24 | -0.18 | 0.61 | 0.03 | 0.09 | 0.53 |
| L. First lumbar | | PC1 | PC2 | PC3 | PC4 | PC5 | PC6 | PC7 |
|  | Proportion of Variance | 60% | 20% | 10% | 7% | 3% | 1% | 0% |
|  | lnL1_VW | 0.40 | 0.18 | 0.61 | 0.65 | 0.04 | 0.09 | 0.00 |
|  | lnL1_VH | 0.48 | 0.03 | 0.00 | -0.30 | 0.17 | -0.08 | -0.80 |
|  | lnL1_CL | -0.27 | 0.58 | 0.38 | -0.27 | 0.08 | -0.60 | 0.03 |
|  | lnL1_CW | 0.23 | 0.13 | -0.03 | -0.08 | -0.96 | -0.09 | -0.02 |
|  | lnL1_CH | 0.45 | 0.58 | -0.58 | 0.10 | 0.19 | -0.01 | 0.30 |
|  | lnL1_ZL | -0.18 | 0.46 | 0.20 | -0.31 | -0.04 | 0.78 | -0.05 |
|  | lnL1_NSH | 0.50 | -0.27 | 0.33 | -0.55 | 0.11 | 0.01 | 0.51 |
| M. Middle lumbar | | PC1 | PC2 | PC3 | PC4 | PC5 | PC6 | PC7 |
|  | Proportion of Variance | 54% | 18% | 14% | 9% | 3% | 1% | 0% |
|  | lnmidL_VW | 0.39 | 0.06 | 0.24 | 0.86 | 0.21 | 0.07 | 0.04 |
|  | lnmidL_VH | 0.36 | -0.02 | 0.37 | -0.25 | 0.08 | -0.08 | -0.81 |
|  | lnmidL_CL | -0.45 | -0.35 | 0.53 | 0.10 | 0.09 | -0.61 | 0.09 |
|  | lnmidL_CW | 0.21 | -0.19 | 0.05 | 0.15 | -0.94 | -0.11 | -0.01 |
|  | lnmidL_CH | 0.48 | -0.77 | -0.12 | -0.20 | 0.22 | 0.02 | 0.26 |
|  | lnmidL_ZL | -0.37 | -0.33 | 0.37 | 0.05 | -0.07 | 0.78 | -0.09 |
|  | lnmidL_NSH | 0.32 | 0.37 | 0.61 | -0.36 | -0.04 | 0.09 | 0.51 |
| N. Last lumbar | | PC1 | PC2 | PC3 | PC4 | PC5 | PC6 | PC7 |
|  | Proportion of Variance | 39% | 29% | 16% | 10% | 4% | 1% | 1% |
|  | lnlastL_VW | 0.22 | 0.37 | 0.52 | 0.66 | 0.33 | 0.03 | 0.07 |
|  | lnlastL_VH | 0.54 | -0.05 | 0.00 | -0.12 | 0.08 | 0.07 | -0.82 |
|  | lnlastL_CL | -0.01 | -0.68 | 0.21 | 0.25 | 0.03 | -0.65 | -0.05 |
|  | lnlastL_CW | 0.20 | 0.09 | 0.04 | 0.31 | -0.92 | -0.01 | 0.00 |
|  | lnlastL_CH | 0.50 | -0.09 | -0.71 | 0.33 | 0.18 | -0.03 | 0.30 |
|  | lnlastL_ZL | -0.02 | -0.61 | 0.14 | 0.20 | 0.00 | 0.75 | 0.05 |
|  | lnlastL_NSH | 0.60 | -0.08 | 0.40 | -0.50 | -0.02 | -0.02 | 0.47 |

**Table S3. Comparisons of the best fitting evolutionary models in simulated datasets of skeletal phenome and size.** Simulated datasets were generated using the parameter estimates of the best-fit model(s) in the empirical dataset for skeletal phenome and skeletal size. Rows in boldface type represent the best-fit model as indicated by ΔAICc < 2.

|  | Model | AICc | ΔAICc | AICcW |
| --- | --- | --- | --- | --- |
| Skeletal phenome: Simulations under OUBMi_EOT_ | | | | |
|  | BM1 | -8073.10 | 114.56 | 0.00 |
|  | trend | -8050.57 | 137.09 | 0.00 |
|  | EB | -8065.71 | 121.95 | 0.00 |
|  | OU1 | -7926.25 | 261.41 | 0.00 |
|  | BMBM_EOT_ | -8113.20 | 74.47 | 0.00 |
|  | OUOU_EOT_ | -7930.12 | 257.55 | 0.00 |
|  | OUBM_EOT_ | -8052.58 | 135.08 | 0.00 |
|  | **OUBMi_EOT_** | **-8187.66** | **0.00** | **>0.99** |
|  | BMBM_MMCT_ | -8166.78 | 20.89 | 0.00 |
|  | OUOU_MMCT_ | -7925.05 | 262.61 | 0.00 |
|  | OUBM_MMCT_ | -8031.74 | 155.92 | 0.00 |
|  | OUBMi_MMCT_ | -8096.72 | 90.94 | 0.00 |
|  | sOU2 | -7931.60 | 256.07 | 0.00 |
|  | sOU3 | -7919.53 | 268.13 | 0.00 |
|  | cOU3 | -7955.28 | 232.39 | 0.00 |
|  | cOU4 | -7942.74 | 244.92 | 0.00 |
| Skeletal phenome: Simulations under OUBMi_MMCT_ | | | | |
|  | BM1 | -8139.69 | 58.28 | 0.00 |
|  | trend | -8114.30 | 83.67 | 0.00 |
|  | EB | -8133.64 | 64.32 | 0.00 |
|  | OU1 | -8094.24 | 103.72 | 0.00 |
|  | BMBM_EOT_ | -8147.70 | 50.26 | 0.00 |
|  | OUOU_EOT_ | -7978.14 | 219.82 | 0.00 |
|  | OUBM_EOT_ | -8049.55 | 148.41 | 0.00 |
|  | OUBMi_EOT_ | -8112.99 | 84.97 | 0.00 |
|  | **BMBM_MMCT_** | **-8197.96** | **0.00** | **0.73** |
|  | OUOU_MMCT_ | -7960.50 | 237.46 | 0.00 |
|  | OUBM_MMCT_ | -8025.59 | 172.37 | 0.00 |
|  | **OUBMi_MMCT_** | **-8196.00** | **1.96** | **0.27** |
|  | sOU2 | -7970.78 | 227.18 | 0.00 |
|  | sOU3 | -7974.81 | 223.15 | 0.00 |
|  | cOU3 | -7971.27 | 226.69 | 0.00 |
|  | cOU4 | -7975.61 | 222.35 | 0.00 |
| Skeletal size: Simulations under OUBMi_MMCT_ | | | | |
|  | BM1 | -1050.46 | 4.23 | 0.03 |
|  | trend | -1048.36 | 6.33 | 0.01 |
|  | EB | -1048.40 | 6.29 | 0.01 |
|  | **OU1** | **-1054.69** | **0.00** | **0.21** |
|  | BMBM_EOT_ | -1048.40 | 6.29 | 0.01 |
|  | **OUOU_EOT_** | **-1053.89** | **0.79** | **0.14** |
|  | OUBM_EOT_ | -1048.48 | 6.21 | 0.01 |
|  | OUBMi_EOT_ | -1049.60 | 5.08 | 0.02 |
|  | BMBM_MMCT_ | -1048.42 | 6.27 | 0.01 |
|  | **OUOU_MMCT_** | **-1052.98** | **1.71** | **0.09** |
|  | OUBM_MMCT_ | -1052.08 | 2.61 | 0.06 |
|  | OUBMi_MMCT_ | -1051.77 | 2.91 | 0.05 |
|  | **sOU2** | **-1053.07** | **1.61** | **0.09** |
|  | sOU3 | -1051.59 | 3.10 | 0.05 |
|  | **cOU3** | **-1054.09** | **0.59** | **0.16** |
|  | cOU4 | -1052.60 | 2.09 | 0.07 |

**Table S4. Phylogenetic signal of skeletal components.**

| Skeletal component | Pagel’s λ | P-value | Blomberg’s K | P-value |
| --- | --- | --- | --- | --- |
| cranium | 0.94 | 0.001 | 0.38 | 0.001 |
| mandible | 1.00 | 0.001 | 0.45 | 0.001 |
| forelimb | 0.44 | 0.001 | 0.36 | 0.001 |
| hindlimb | 0.44 | 0.001 | 0.36 | 0.001 |
| third cervical | 1.00 | 0.001 | 0.42 | 0.001 |
| fifth cervical | 0.97 | 0.001 | 0.39 | 0.001 |
| first thoracic | 1.00 | 0.001 | 0.42 | 0.001 |
| middle thoracic | 1.00 | 0.001 | 0.42 | 0.001 |
| diaphragmatic | 0.97 | 0.001 | 0.39 | 0.001 |
| last thoracic | 0.97 | 0.001 | 0.39 | 0.001 |
| first lumbar | 0.97 | 0.001 | 0.39 | 0.001 |
| middle lumbar | 0.97 | 0.001 | 0.39 | 0.001 |
| last lumbar | 0.97 | 0.001 | 0.39 | 0.001 |

**Table S5. Comparisons of the best-fitting evolutionary models in each of the skeletal components.** Rows in boldface type represent the best supported model as indicated by ΔAICc < 2.

| Skeletal component | Model | AICc | ΔAICc | AICcW |
| --- | --- | --- | --- | --- |
| cranium | BM1 | -1261.44 | 13.67 | 0.00 |
| (PCs 1-5: 95.8% of variance) | trend | -1269.25 | 5.86 | 0.02 |
|  | EB | -1248.77 | 26.34 | 0.00 |
|  | OU1 | -1254.84 | 20.27 | 0.00 |
|  | **BMBM_EOT_** | **-1275.11** | **0.00** | **0.37** |
|  | OUOU_EOT_ | -1266.64 | 8.48 | 0.01 |
|  | **OUBM_EOT_** | **-1273.68** | **1.43** | **0.18** |
|  | OUBMi_EOT_ | -1271.22 | 3.89 | 0.05 |
|  | BMBM_MMCT_ | -1269.64 | 5.47 | 0.02 |
|  | OUOU_MMCT_ | -1269.79 | 5.33 | 0.03 |
|  | OUBM_MMCT_ | -1270.62 | 4.49 | 0.04 |
|  | **OUBMi_MMCT_** | **-1274.53** | **0.59** | **0.28** |
|  | sOU2 | -1252.98 | 22.13 | 0.00 |
|  | sOU3 | -1250.34 | 24.77 | 0.00 |
|  | cOU3 | -1244.84 | 30.28 | 0.00 |
|  | cOU4 | -1242.04 | 33.07 | 0.00 |
| mandible | BM1 | -1415.08 | 24.97 | 0.00 |
| (PCs 1-5: 94.7% of variance) | trend | -1420.77 | 19.28 | 0.00 |
|  | EB | -1407.71 | 32.34 | 0.00 |
|  | OU1 | -1424.13 | 15.93 | 0.00 |
|  | BMBM_EOT_ | -1412.70 | 27.35 | 0.00 |
|  | **OUOU_EOT_** | **-1440.05** | **0.00** | **0.54** |
|  | OUBM_EOT_ | -1418.68 | 21.37 | 0.00 |
|  | OUBMi_EOT_ | -1423.66 | 16.40 | 0.00 |
|  | BMBM_MMCT_ | -1432.48 | 7.57 | 0.01 |
|  | OUOU_MMCT_ | -1425.60 | 14.45 | 0.00 |
|  | OUBM_MMCT_ | -1430.67 | 9.39 | 0.00 |
|  | **OUBMi_MMCT_** | **-1438.74** | **1.32** | **0.28** |
|  | sOU2 | -1421.30 | 18.75 | 0.00 |
|  | sOU3 | -1437.59 | 2.47 | 0.16 |
|  | cOU3 | -1413.82 | 26.24 | 0.00 |
|  | cOU4 | -1414.49 | 25.57 | 0.00 |
| forelimb | BM1 | -1251.64 | 26.79 | 0.00 |
| (PCs 1-7: 95.8% of variance) | trend | -1245.17 | 33.26 | 0.00 |
|  | EB | -1230.38 | 48.05 | 0.00 |
|  | OU1 | -1265.18 | 13.25 | 0.00 |
|  | BMBM_EOT_ | -1261.49 | 16.94 | 0.00 |
|  | OUOU_EOT_ | -1255.21 | 23.22 | 0.00 |
|  | OUBM_EOT_ | -1263.97 | 14.46 | 0.00 |
|  | OUBMi_EOT_ | -1259.19 | 19.24 | 0.00 |
|  | BMBM_MMCT_ | -1272.29 | 6.15 | 0.04 |
|  | **OUOU_MMCT_** | **-1278.43** | **0.00** | **0.84** |
|  | OUBM_MMCT_ | -1267.05 | 11.38 | 0.00 |
|  | OUBMi_MMCT_ | -1274.39 | 4.05 | 0.11 |
|  | sOU2 | -1242.45 | 35.98 | 0.00 |
|  | sOU3 | -1240.83 | 37.60 | 0.00 |
|  | cOU3 | -1237.59 | 40.84 | 0.00 |
|  | cOU4 | -1223.66 | 54.77 | 0.00 |
| hindlimb | BM1 | -709.63 | 241.05 | 0.00 |
| (PCs 1-7: 95.1% of variance) | trend | -713.87 | 236.81 | 0.00 |
|  | EB | -713.79 | 236.90 | 0.00 |
|  | OU1 | -810.14 | 140.54 | 0.00 |
|  | BMBM_EOT_ | -864.59 | 86.09 | 0.00 |
|  | OUOU_EOT_ | -829.62 | 121.07 | 0.00 |
|  | OUBM_EOT_ | -741.06 | 209.62 | 0.00 |
|  | OUBMi_EOT_ | -848.70 | 101.98 | 0.00 |
|  | BMBM_MMCT_ | -928.83 | 21.85 | 0.00 |
|  | OUOU_MMCT_ | -842.11 | 108.57 | 0.00 |
|  | OUBM_MMCT_ | -754.35 | 196.34 | 0.00 |
|  | **OUBMi_MMCT_** | **-950.68** | **0.00** | **1.00** |
|  | sOU2 | -811.43 | 139.25 | 0.00 |
|  | sOU3 | -801.04 | 149.64 | 0.00 |
|  | cOU3 | -799.59 | 151.09 | 0.00 |
|  | cOU4 | -788.50 | 162.19 | 0.00 |
| third cervical vertebrae | BM1 | -416.65 | 25.87 | 0.00 |
| (PCs 1-3: 90.9% of variance) | trend | -417.34 | 25.18 | 0.00 |
|  | EB | -414.58 | 27.94 | 0.00 |
|  | OU1 | -428.38 | 14.14 | 0.00 |
|  | **BMBM_EOT_** | **-440.71** | **1.81** | **0.28** |
|  | OUOU_EOT_ | -428.10 | 14.43 | 0.00 |
|  | **OUBM_EOT_** | **-442.52** | **0.00** | **0.69** |
|  | OUBMi_EOT_ | -436.32 | 6.20 | 0.03 |
|  | BMBM_MMCT_ | -416.73 | 25.79 | 0.00 |
|  | OUOU_MMCT_ | -430.28 | 12.24 | 0.00 |
|  | OUBM_MMCT_ | -425.03 | 17.49 | 0.00 |
|  | OUBMi_MMCT_ | -425.00 | 17.52 | 0.00 |
|  | sOU2 | -423.57 | 18.95 | 0.00 |
|  | sOU3 | -420.20 | 22.32 | 0.00 |
|  | cOU3 | -417.26 | 25.26 | 0.00 |
|  | cOU4 | -413.88 | 28.65 | 0.00 |
| fifth cervical vertebrae | BM1 | -851.20 | 47.37 | 0.00 |
| (PCs 1-4: 95.9% of variance) | trend | -852.01 | 46.56 | 0.00 |
|  | EB | -848.54 | 50.03 | 0.00 |
|  | OU1 | -873.52 | 25.05 | 0.00 |
|  | BMBM_EOT_ | -884.43 | 14.15 | 0.00 |
|  | OUOU_EOT_ | -872.71 | 25.86 | 0.00 |
|  | OUBM_EOT_ | -883.74 | 14.84 | 0.00 |
|  | OUBMi_EOT_ | -874.21 | 24.37 | 0.00 |
|  | BMBM_MMCT_ | -884.20 | 14.38 | 0.00 |
|  | OUOU_MMCT_ | -872.05 | 26.52 | 0.00 |
|  | **OUBM_MMCT_** | **-898.47** | **0.10** | **0.49** |
|  | **OUBMi_MMCT_** | **-898.57** | **0.00** | **0.51** |
|  | sOU2 | -872.13 | 26.44 | 0.00 |
|  | sOU3 | -859.32 | 39.26 | 0.00 |
|  | cOU3 | -859.85 | 38.72 | 0.00 |
|  | cOU4 | -850.85 | 47.72 | 0.00 |
| first thoracic vertebrae | BM1 | -589.31 | 24.57 | 0.00 |
| (PCs 1-3: 92.4% of variance) | trend | -586.16 | 27.72 | 0.00 |
|  | EB | -588.48 | 25.40 | 0.00 |
|  | OU1 | -568.63 | 45.25 | 0.00 |
|  | **BMBM_EOT_** | **-613.88** | **0.00** | **0.81** |
|  | OUOU_EOT_ | -581.86 | 32.03 | 0.00 |
|  | OUBM_EOT_ | -594.24 | 19.64 | 0.00 |
|  | OUBMi_EOT_ | -609.06 | 4.82 | 0.07 |
|  | BMBM_MMCT_ | -597.97 | 15.91 | 0.00 |
|  | OUOU_MMCT_ | -580.94 | 32.94 | 0.00 |
|  | OUBM_MMCT_ | -586.35 | 27.53 | 0.00 |
|  | OUBMi_MMCT_ | -610.00 | 3.88 | 0.12 |
|  | sOU2 | -568.42 | 45.46 | 0.00 |
|  | sOU3 | -562.12 | 51.76 | 0.00 |
|  | cOU3 | -562.25 | 51.63 | 0.00 |
|  | cOU4 | -556.68 | 57.20 | 0.00 |
| middle thoracic vertebrae | BM1 | -624.34 | 23.14 | 0.00 |
| (PCs 1-3: 89.9% of variance) | trend | -623.56 | 23.93 | 0.00 |
|  | EB | -622.26 | 25.22 | 0.00 |
|  | OU1 | -608.43 | 39.06 | 0.00 |
|  | **BMBM_EOT_** | **-646.01** | **1.48** | **0.30** |
|  | OUOU_EOT_ | -615.92 | 31.56 | 0.00 |
|  | OUBM_EOT_ | -633.82 | 13.67 | 0.00 |
|  | **OUBMi_EOT_** | **-647.48** | **0.00** | **0.63** |
|  | BMBM_MMCT_ | -635.07 | 12.41 | 0.00 |
|  | OUOU_MMCT_ | -616.86 | 30.63 | 0.00 |
|  | OUBM_MMCT_ | -624.25 | 23.23 | 0.00 |
|  | OUBMi_MMCT_ | -642.95 | 4.53 | 0.07 |
|  | sOU2 | -612.25 | 35.24 | 0.00 |
|  | sOU3 | -607.40 | 40.09 | 0.00 |
|  | cOU3 | -606.00 | 41.48 | 0.00 |
|  | cOU4 | -601.19 | 46.29 | 0.00 |
| diaphragmatic vertebrae | BM1 | -906.03 | 56.31 | 0.00 |
| (PCs 1-4: 96.3% of variance) | trend | -908.20 | 54.14 | 0.00 |
|  | EB | -896.03 | 66.32 | 0.00 |
|  | OU1 | -939.27 | 23.07 | 0.00 |
|  | BMBM_EOT_ | -933.32 | 29.03 | 0.00 |
|  | OUOU_EOT_ | -923.89 | 38.45 | 0.00 |
|  | OUBM_EOT_ | -935.17 | 27.17 | 0.00 |
|  | OUBMi_EOT_ | -946.80 | 15.54 | 0.00 |
|  | BMBM_MMCT_ | -956.30 | 6.05 | 0.05 |
|  | OUOU_MMCT_ | -944.59 | 17.75 | 0.00 |
|  | OUBM_MMCT_ | -945.65 | 16.70 | 0.00 |
|  | **OUBMi_MMCT_** | **-962.34** | **0.00** | **0.95** |
|  | sOU2 | -939.91 | 22.43 | 0.00 |
|  | sOU3 | -939.10 | 23.24 | 0.00 |
|  | cOU3 | -931.33 | 31.01 | 0.00 |
|  | cOU4 | -929.30 | 33.05 | 0.00 |
| last thoracic vertebrae | BM1 | -1161.77 | 40.01 | 0.00 |
| (PCs 1-4: 96.8% of variance) | trend | -1159.90 | 41.88 | 0.00 |
|  | EB | -1147.73 | 54.04 | 0.00 |
|  | OU1 | -1173.04 | 28.73 | 0.00 |
|  | BMBM_EOT_ | -1169.26 | 32.51 | 0.00 |
|  | OUOU_EOT_ | -1170.37 | 31.41 | 0.00 |
|  | OUBM_EOT_ | -1171.52 | 30.26 | 0.00 |
|  | OUBMi_EOT_ | -1169.21 | 32.57 | 0.00 |
|  | BMBM_MMCT_ | -1180.73 | 21.05 | 0.00 |
|  | OUOU_MMCT_ | -1168.72 | 33.06 | 0.00 |
|  | OUBM_MMCT_ | -1184.68 | 17.10 | 0.00 |
|  | **OUBMi_MMCT_** | **-1201.78** | **0.00** | **1.00** |
|  | sOU2 | -1169.95 | 31.82 | 0.00 |
|  | sOU3 | -1164.97 | 36.80 | 0.00 |
|  | cOU3 | -1162.34 | 39.44 | 0.00 |
|  | cOU4 | -1156.60 | 45.18 | 0.00 |
| first lumbar vertebrae | BM1 | -965.81 | 59.19 | 0.00 |
| (PCs 1-4: 96.4% of variance) | trend | -965.53 | 59.47 | 0.00 |
|  | EB | -963.73 | 61.27 | 0.00 |
|  | OU1 | -978.05 | 46.96 | 0.00 |
|  | BMBM_EOT_ | -996.10 | 28.90 | 0.00 |
|  | OUOU_EOT_ | -975.81 | 49.20 | 0.00 |
|  | OUBM_EOT_ | -984.26 | 40.75 | 0.00 |
|  | OUBMi_EOT_ | -988.33 | 36.67 | 0.00 |
|  | BMBM_MMCT_ | -993.24 | 31.77 | 0.00 |
|  | OUOU_MMCT_ | -979.71 | 45.29 | 0.00 |
|  | OUBM_MMCT_ | -993.78 | 31.22 | 0.00 |
|  | **OUBMi_MMCT_** | **-1025.00** | **0.00** | **1.00** |
|  | sOU2 | -977.82 | 47.18 | 0.00 |
|  | sOU3 | -982.25 | 42.75 | 0.00 |
|  | cOU3 | -971.68 | 53.32 | 0.00 |
|  | cOU4 | -961.88 | 63.12 | 0.00 |
| middle lumbar vertebrae | BM1 | -869.68 | 16.89 | 0.00 |
| (PCs 1-4: 95.6% of variance) | trend | -870.41 | 16.16 | 0.00 |
|  | EB | -863.31 | 23.26 | 0.00 |
|  | OU1 | -878.83 | 7.74 | 0.02 |
|  | BMBM_EOT_ | -872.89 | 13.69 | 0.00 |
|  | OUOU_EOT_ | -882.73 | 3.84 | 0.11 |
|  | OUBM_EOT_ | -875.23 | 11.35 | 0.00 |
|  | OUBMi_EOT_ | -874.24 | 12.33 | 0.00 |
|  | BMBM_MMCT_ | -863.63 | 22.94 | 0.00 |
|  | OUOU_MMCT_ | -882.83 | 3.74 | 0.12 |
|  | OUBM_MMCT_ | -873.52 | 13.05 | 0.00 |
|  | **OUBMi_MMCT_** | **-886.57** | **0.00** | **0.75** |
|  | sOU2 | -876.36 | 10.21 | 0.00 |
|  | sOU3 | -869.87 | 16.71 | 0.00 |
|  | cOU3 | -868.95 | 17.62 | 0.00 |
|  | cOU4 | -862.23 | 24.35 | 0.00 |
| last lumbar vertebrae | BM1 | -692.02 | 82.52 | 0.00 |
| (PCs 1-4: 94.9% of variance) | trend | -692.30 | 82.24 | 0.00 |
|  | EB | -687.96 | 86.58 | 0.00 |
|  | **OU1** | **-774.42** | **0.12** | **0.36** |
|  | BMBM_EOT_ | -718.99 | 55.55 | 0.00 |
|  | **OUOU_EOT_** | **-774.54** | **0.00** | **0.39** |
|  | OUBM_EOT_ | -708.20 | 66.34 | 0.00 |
|  | OUBMi_EOT_ | -735.16 | 39.38 | 0.00 |
|  | BMBM_MMCT_ | -730.45 | 44.09 | 0.00 |
|  | **OUOU_MMCT_** | **-773.38** | **1.16** | **0.22** |
|  | OUBM_MMCT_ | -726.64 | 47.90 | 0.00 |
|  | OUBMi_MMCT_ | -753.90 | 20.64 | 0.00 |
|  | sOU2 | -768.08 | 6.46 | 0.02 |
|  | sOU3 | -768.13 | 6.41 | 0.02 |
|  | cOU3 | -760.51 | 14.03 | 0.00 |
|  | cOU4 | -761.07 | 13.47 | 0.00 |

**Table S6. Parameter estimates from best** **fitting evolutionary models of skeletal components.** Mean parameters were calculated from all PC axes used in each model. θ = theta; σ^2^ = evolutionary rate; α = strength of selection; SV = stationary variances (σ^2^/2α); half-life = phylogenetic half-life (ln(2)/α). In all models, the “mean θ” and “mean σ^2^” columns shows the mean θ and σ^2^ values, respectively, from before the EOT or MMCT, and the “parameter” column shows the mean θ or σ^2^ values from after the EOT/MMCT.

| skeletal component | best model | mean θ | mean σ^2^ | mean α | SV | halflife | parameter |
| --- | --- | --- | --- | --- | --- | --- | --- |
| cranium | **BMBM_EOT_** | -0.022 | <0.001 | NA | NA | NA | σ^2^_post-EOT_ = 0.001 |
| mandible | **OUOU_EOT_** | 0.251 | 0.001 | 0.027 | 0.044 | 25.300 | θ_post-EOT_ = -0.358 |
| forelimb | **OUOU_MMCT_** | -0.084 | 0.002 | 0.034 | 0.069 | 20.519 | θ_post-MMCT_ = 1.538 |
| hindlimb | **OUBMi_MMCT_** | -0.002 | 0.007 | 0.094 | 0.062 | 7.381 | σ^2^_post-MMCT_ = 0.002 |
| third cervical | **OUBM_EOT_** | 0.056 | 0.002 | 1.161 | 0.001 | 0.597 | σ^2^_post-EOT_ = 0.001 |
| fifth cervical | **OUBMi_MMCT_** | 0.071 | 0.002 | 0.098 | 0.013 | 7.079 | σ^2^_post-MMCT_ = 0.001 |
| first thoracic | **BMBM_EOT_** | -0.022 | <0.001 | NA | NA | NA | σ^2^_post-EOT_ = 0.001 |
| middle thoracic | **OUBMi_EOT_** | 0.004 | 0.004 | 0.118 | 0.015 | 5.853 | σ^2^_post-EOT_ < 0.001 |
| diaphragmatic | **OUBMi_MMCT_** | 0.074 | 0.002 | 0.055 | 0.043 | 12.504 | σ^2^_post-MMCT_ = 0.001 |
| last thoracic | **OUBMi_MMCT_** | 0.029 | 0.002 | 0.094 | 0.021 | 7.352 | σ^2^_post-MMCT_ < 0.001 |
| first lumbar | **OUBMi_MMCT_** | 0.056 | 0.003 | 0.124 | 0.024 | 5.607 | σ^2^_post-MMCT_ < 0.001 |
| middle lumbar | **OUBMi_MMCT_** | -0.011 | 0.004 | 0.089 | 0.048 | 7.745 | σ^2^_post-MMCT_ < 0.001 |
| last lumbar | **OUOU_EOT_** | -0.412 | 0.013 | 0.432 | 0.006 | 1.600 | θ_post-EOT_ = -3.34 |

**Table S7. Comparisons of the best fitting evolutionary models in simulated datasets of skeletal components.** Simulated data were generated using parameter estimates of the best-fit model in the empirical dataset of each skeletal component. Rows in boldface type represent the best supported model as indicated by ΔAICc < 2.

| Skeletal component | Model | AICc | ΔAICc | AICcW |
| --- | --- | --- | --- | --- |
| cranium | **BM1** | **-7943.85** | **0.00** | **0.95** |
| (PCs 1-5: 95.8% of variance) | trend | -7912.66 | 31.19 | 0.00 |
|  | EB | -7937.96 | 5.89 | 0.05 |
|  | OU1 | -7549.43 | 394.42 | 0.00 |
|  | BMBM_EOT_ | -7912.09 | 31.77 | 0.00 |
|  | OUOU_EOT_ | -7548.59 | 395.26 | 0.00 |
|  | OUBM_EOT_ | -7837.82 | 106.03 | 0.00 |
|  | OUBMi_EOT_ | -7802.18 | 141.67 | 0.00 |
|  | BMBM_MMCT_ | -7912.40 | 31.45 | 0.00 |
|  | OUOU_MMCT_ | -7546.25 | 397.60 | 0.00 |
|  | OUBM_MMCT_ | -7778.95 | 164.90 | 0.00 |
|  | OUBMi_MMCT_ | -7762.92 | 180.93 | 0.00 |
|  | sOU2 | -7552.20 | 391.65 | 0.00 |
|  | sOU3 | -7550.06 | 393.79 | 0.00 |
|  | cOU3 | -7548.56 | 395.29 | 0.00 |
|  | cOU4 | -7546.74 | 397.11 | 0.00 |
| mandible | BM1 | -6668.65 | 496.35 | 0.00 |
| (PCs 1-5: 94.7% of variance) | trend | -6544.95 | 620.05 | 0.00 |
|  | EB | -6867.86 | 297.13 | 0.00 |
|  | OU1 | -6444.89 | 720.11 | 0.00 |
|  | BMBM_EOT_ | -6636.88 | 528.11 | 0.00 |
|  | **OUOU_EOT_** | **-7164.99** | **0.00** | **1.00** |
|  | OUBM_EOT_ | -6316.79 | 848.20 | 0.00 |
|  | OUBMi_EOT_ | -6284.02 | 880.98 | 0.00 |
|  | BMBM_MMCT_ | -6636.88 | 528.11 | 0.00 |
|  | OUOU_MMCT_ | -6758.49 | 406.50 | 0.00 |
|  | OUBM_MMCT_ | -5941.18 | 1223.82 | 0.00 |
|  | OUBMi_MMCT_ | -5908.40 | 1256.59 | 0.00 |
|  | sOU2 | -6747.34 | 417.65 | 0.00 |
|  | sOU3 | -6762.94 | 402.05 | 0.00 |
|  | cOU3 | -6777.11 | 387.88 | 0.00 |
|  | cOU4 | -6776.91 | 388.08 | 0.00 |
| forelimb | BM1 | -5670.75 | 590.85 | 0.00 |
| (PCs 1-7: 95.8% of variance) | trend | -5629.16 | 632.44 | 0.00 |
|  | EB | -5704.55 | 557.04 | 0.00 |
|  | OU1 | -5562.22 | 699.37 | 0.00 |
|  | BMBM_EOT_ | -5709.74 | 551.85 | 0.00 |
|  | OUOU_EOT_ | -5789.87 | 471.72 | 0.00 |
|  | OUBM_EOT_ | -5545.92 | 715.67 | 0.00 |
|  | OUBMi_EOT_ | -5677.58 | 584.02 | 0.00 |
|  | BMBM_MMCT_ | -5870.37 | 391.23 | 0.00 |
|  | **OUOU_MMCT_** | **-6261.59** | **0.00** | **1.00** |
|  | OUBM_MMCT_ | -5475.40 | 786.19 | 0.00 |
|  | OUBMi_MMCT_ | -5833.46 | 428.13 | 0.00 |
|  | sOU2 | -5682.51 | 579.08 | 0.00 |
|  | sOU3 | -5686.85 | 574.74 | 0.00 |
|  | cOU3 | -5677.44 | 584.15 | 0.00 |
|  | cOU4 | -5687.28 | 574.32 | 0.00 |
| hindlimb | BM1 | -6910.66 | 97.87 | 0.00 |
| (PCs 1-7: 95.1% of variance) | trend | -6895.98 | 112.55 | 0.00 |
|  | EB | -6920.79 | 87.74 | 0.00 |
|  | OU1 | -6741.39 | 267.15 | 0.00 |
|  | BMBM_EOT_ | -6878.90 | 129.64 | 0.00 |
|  | OUOU_EOT_ | -6749.68 | 258.86 | 0.00 |
|  | OUBM_EOT_ | -6857.03 | 151.51 | 0.00 |
|  | OUBMi_EOT_ | -6821.42 | 187.11 | 0.00 |
|  | BMBM_MMCT_ | -6879.62 | 128.92 | 0.00 |
|  | OUOU_MMCT_ | -6747.47 | 261.07 | 0.00 |
|  | OUBM_MMCT_ | -6809.82 | 198.71 | 0.00 |
|  | **OUBMi_MMCT_** | **-7008.53** | **0.00** | **1.00** |
|  | sOU2 | -6747.54 | 260.99 | 0.00 |
|  | sOU3 | -6745.85 | 262.69 | 0.00 |
|  | cOU3 | -6741.43 | 267.11 | 0.00 |
|  | cOU4 | -6736.29 | 272.25 | 0.00 |
| third cervical vertebrae | BM1 | -4316.87 | 20.64 | 0.00 |
| (PCs 1-3: 90.9% of variance) | trend | -4310.16 | 27.34 | 0.00 |
|  | EB | -4314.80 | 22.71 | 0.00 |
|  | OU1 | -4250.82 | 86.68 | 0.00 |
|  | BMBM_EOT_ | -4306.05 | 31.46 | 0.00 |
|  | OUOU_EOT_ | -4247.44 | 90.07 | 0.00 |
|  | **OUBM_EOT_** | **-4337.50** | **0.00** | **0.99** |
|  | OUBMi_EOT_ | -4326.45 | 11.05 | 0.00 |
|  | BMBM_MMCT_ | -4306.36 | 31.14 | 0.00 |
|  | OUOU_MMCT_ | -4248.42 | 89.08 | 0.00 |
|  | OUBM_MMCT_ | -4324.93 | 12.57 | 0.00 |
|  | OUBMi_MMCT_ | -4316.15 | 21.35 | 0.00 |
|  | sOU2 | -4245.62 | 91.88 | 0.00 |
|  | sOU3 | -4240.54 | 96.97 | 0.00 |
|  | cOU3 | -4251.60 | 85.91 | 0.00 |
|  | cOU4 | -4246.05 | 91.45 | 0.00 |
| fifth cervical vertebrae | BM1 | -6196.14 | 18.85 | 0.00 |
| (PCs 1-4: 95.9% of variance) | trend | -6186.05 | 28.94 | 0.00 |
|  | EB | -6192.09 | 22.90 | 0.00 |
|  | OU1 | -6167.69 | 47.29 | 0.00 |
|  | BMBM_EOT_ | -6176.12 | 38.87 | 0.00 |
|  | OUOU_EOT_ | -6162.78 | 52.21 | 0.00 |
|  | OUBM_EOT_ | -6202.17 | 12.82 | 0.00 |
|  | OUBMi_EOT_ | -6180.44 | 34.54 | 0.00 |
|  | BMBM_MMCT_ | -6177.20 | 37.78 | 0.00 |
|  | OUOU_MMCT_ | -6164.46 | 50.53 | 0.00 |
|  | **OUBM_MMCT_** | **-6214.99** | **0.00** | **1.00** |
|  | OUBMi_MMCT_ | -6202.84 | 12.15 | 0.00 |
|  | sOU2 | -6163.01 | 51.98 | 0.00 |
|  | sOU3 | -6155.57 | 59.42 | 0.00 |
|  | cOU3 | -6159.04 | 55.95 | 0.00 |
|  | cOU4 | -6151.90 | 63.09 | 0.00 |
| first thoracic vertebrae | **BM1** | -4547.65 | 0.00 | 0.53 |
| (PCs 1-3: 92.4% of variance) | trend | -4540.12 | 7.53 | 0.01 |
|  | **EB** | -4547.34 | 0.31 | 0.46 |
|  | OU1 | -4415.84 | 131.81 | 0.00 |
|  | BMBM_EOT_ | -4535.13 | 12.52 | 0.00 |
|  | OUOU_EOT_ | -4417.96 | 129.69 | 0.00 |
|  | OUBM_EOT_ | -4508.66 | 39.00 | 0.00 |
|  | OUBMi_EOT_ | -4494.62 | 53.03 | 0.00 |
|  | BMBM_MMCT_ | -4535.13 | 12.52 | 0.00 |
|  | OUOU_MMCT_ | -4415.29 | 132.37 | 0.00 |
|  | OUBM_MMCT_ | -4463.63 | 84.02 | 0.00 |
|  | OUBMi_MMCT_ | -4476.49 | 71.16 | 0.00 |
|  | sOU2 | -4413.06 | 134.59 | 0.00 |
|  | sOU3 | -4412.60 | 135.06 | 0.00 |
|  | cOU3 | -4418.03 | 129.63 | 0.00 |
|  | cOU4 | -4424.20 | 123.46 | 0.00 |
| middle thoracic vertebrae | **BM1** | **-4623.47** | **0.00** | **0.56** |
| (PCs 1-3: 89.9% of variance) | trend | -4616.37 | 7.10 | 0.02 |
|  | **EB** | **-4622.92** | **0.55** | **0.42** |
|  | OU1 | -4521.19 | 102.28 | 0.00 |
|  | BMBM_EOT_ | -4611.92 | 11.55 | 0.00 |
|  | OUOU_EOT_ | -4519.62 | 103.85 | 0.00 |
|  | OUBM_EOT_ | -4614.03 | 9.44 | 0.00 |
|  | OUBMi_EOT_ | -4602.11 | 21.36 | 0.00 |
|  | BMBM_MMCT_ | -4611.36 | 12.11 | 0.00 |
|  | OUOU_MMCT_ | -4519.53 | 103.94 | 0.00 |
|  | OUBM_MMCT_ | -4584.48 | 38.99 | 0.00 |
|  | OUBMi_MMCT_ | -4573.11 | 50.36 | 0.00 |
|  | sOU2 | -4516.56 | 106.91 | 0.00 |
|  | sOU3 | -4513.81 | 109.66 | 0.00 |
|  | cOU3 | -4520.16 | 103.31 | 0.00 |
|  | cOU4 | -4516.80 | 106.67 | 0.00 |
| diaphragmatic vertebrae | BM1 | -6306.99 | 165.10 | 0.00 |
| (PCs 1-4: 96.3% of variance) | trend | -6295.85 | 176.25 | 0.00 |
|  | EB | -6312.88 | 159.21 | 0.00 |
|  | OU1 | -6076.20 | 395.89 | 0.00 |
|  | BMBM_EOT_ | -6286.29 | 185.80 | 0.00 |
|  | OUOU_EOT_ | -6069.73 | 402.36 | 0.00 |
|  | OUBM_EOT_ | -6252.80 | 219.30 | 0.00 |
|  | OUBMi_EOT_ | -6234.09 | 238.00 | 0.00 |
|  | **BMBM_MMCT_** | **-6470.11** | **1.99** | **0.27** |
|  | OUOU_MMCT_ | -6071.98 | 400.12 | 0.00 |
|  | OUBM_MMCT_ | -6212.64 | 259.46 | 0.00 |
|  | **OUBMi_MMCT_** | **-6472.09** | **0.00** | **0.73** |
|  | sOU2 | -6069.17 | 402.92 | 0.00 |
|  | sOU3 | -6064.49 | 407.61 | 0.00 |
|  | cOU3 | -6066.79 | 405.30 | 0.00 |
|  | cOU4 | -6063.01 | 409.08 | 0.00 |
| last thoracic vertebrae | BM1 | -6569.08 | 4.87 | 0.08 |
| (PCs 1-4: 96.8% of variance) | trend | -6556.52 | 17.43 | 0.00 |
|  | **EB** | **-6573.95** | **0.00** | **0.92** |
|  | OU1 | -6443.05 | 130.90 | 0.00 |
|  | BMBM_EOT_ | -6548.06 | 25.88 | 0.00 |
|  | OUOU_EOT_ | -6444.90 | 129.04 | 0.00 |
|  | OUBM_EOT_ | -6526.92 | 47.02 | 0.00 |
|  | OUBMi_EOT_ | -6503.53 | 70.41 | 0.00 |
|  | BMBM_MMCT_ | -6555.39 | 18.55 | 0.00 |
|  | OUOU_MMCT_ | -6446.84 | 127.11 | 0.00 |
|  | OUBM_MMCT_ | -6515.38 | 58.56 | 0.00 |
|  | OUBMi_MMCT_ | -6518.37 | 55.57 | 0.00 |
|  | sOU2 | -6442.13 | 131.81 | 0.00 |
|  | sOU3 | -6440.80 | 133.15 | 0.00 |
|  | cOU3 | -6440.33 | 133.61 | 0.00 |
|  | cOU4 | -6438.01 | 135.93 | 0.00 |
| first lumbar vertebrae | **BM1** | **-6279.85** | **0.00** | **0.82** |
| (PCs 1-4: 96.4% of variance) | trend | -6268.06 | 11.79 | 0.00 |
|  | EB | -6276.78 | 3.07 | 0.18 |
|  | OU1 | -6195.78 | 84.07 | 0.00 |
|  | BMBM_EOT_ | -6258.83 | 21.02 | 0.00 |
|  | OUOU_EOT_ | -6193.28 | 86.56 | 0.00 |
|  | OUBM_EOT_ | -6245.54 | 34.30 | 0.00 |
|  | OUBMi_EOT_ | -6223.97 | 55.88 | 0.00 |
|  | BMBM_MMCT_ | -6258.83 | 21.02 | 0.00 |
|  | OUOU_MMCT_ | -6189.01 | 90.84 | 0.00 |
|  | OUBM_MMCT_ | -6231.77 | 48.07 | 0.00 |
|  | OUBMi_MMCT_ | -6210.20 | 69.65 | 0.00 |
|  | sOU2 | -6198.44 | 81.41 | 0.00 |
|  | sOU3 | -6190.91 | 88.94 | 0.00 |
|  | cOU3 | -6201.74 | 78.11 | 0.00 |
|  | cOU4 | -6193.83 | 86.01 | 0.00 |
| middle lumbar vertebrae | **BM1** | **-6185.00** | **0.00** | **0.90** |
| (PCs 1-4: 95.6% of variance) | trend | -6175.31 | 9.69 | 0.01 |
|  | EB | -6180.41 | 4.59 | 0.09 |
|  | OU1 | -6025.59 | 159.41 | 0.00 |
|  | BMBM_EOT_ | -6163.98 | 21.02 | 0.00 |
|  | OUOU_EOT_ | -6020.93 | 164.07 | 0.00 |
|  | OUBM_EOT_ | -6169.86 | 15.14 | 0.00 |
|  | OUBMi_EOT_ | -6148.29 | 36.71 | 0.00 |
|  | BMBM_MMCT_ | -6163.98 | 21.02 | 0.00 |
|  | OUOU_MMCT_ | -6021.48 | 163.52 | 0.00 |
|  | OUBM_MMCT_ | -6143.25 | 41.76 | 0.00 |
|  | OUBMi_MMCT_ | -6121.68 | 63.33 | 0.00 |
|  | sOU2 | -6019.08 | 165.92 | 0.00 |
|  | sOU3 | -6011.59 | 173.41 | 0.00 |
|  | cOU3 | -6014.73 | 170.27 | 0.00 |
|  | cOU4 | -6020.83 | 164.17 | 0.00 |
| last lumbar vertebrae | BM1 | -5213.77 | 443.54 | 0.00 |
| (PCs 1-4: 94.9% of variance) | trend | -5136.61 | 520.70 | 0.00 |
|  | EB | -5222.39 | 434.92 | 0.00 |
|  | OU1 | -5133.97 | 523.34 | 0.00 |
|  | BMBM_EOT_ | -5192.75 | 464.56 | 0.00 |
|  | OUOU_EOT_ | -5654.55 | 2.76 | 0.19 |
|  | OUBM_EOT_ | -5116.92 | 540.39 | 0.00 |
|  | OUBMi_EOT_ | -5152.79 | 504.52 | 0.00 |
|  | BMBM_MMCT_ | -5278.24 | 379.07 | 0.00 |
|  | OUOU_MMCT_ | -5414.53 | 242.78 | 0.00 |
|  | OUBM_MMCT_ | -5027.05 | 630.26 | 0.00 |
|  | OUBMi_MMCT_ | -5227.23 | 430.08 | 0.00 |
|  | **sOU2** | **-5657.31** | **0.00** | **0.75** |
|  | sOU3 | -5652.41 | 4.91 | 0.06 |
|  | cOU3 | -5416.64 | 240.67 | 0.00 |
|  | cOU4 | -5408.55 | 248.77 | 0.00 |

**Table S8. Extinct species that were added to the phylogeny**. PBDB = Paleobiology Database.

| **Species** | **Phylogenetic position** | **Age Range** |
| --- | --- | --- |
| *Crassidia intermedia* | Basal caniform (sister to *Daphoenus*) (1) | 18 Ma (Florida Museum) |
| *Corumictis wolsani* | Stem to Mustelidae (sister to *Promartes*) (2) | 30.8-18.5 Ma (PBDB) |
| *Cyonasua brevirostris* | Sister to genus *Nasua* (3) | 7.24-3 Ma (PBDB) |
| *Eoarctos vorax* | Sister to Ursidae (4) | 32.0–31.4 Ma (4) |
| *Puma pardoides* | Sister to *Puma concolor* (5) | L Plio to E Pleist. (~3-2 Ma) (5) |
| *Satherium ingens* | Sister to genus *Pteronura* (6) | 4.9-1.8 Ma (PBDB) |
| *Neovulpavus mccarrolli* | Stem carnivoramorphan (7) | Early Uintan (Ui1b) NALMA (46.2 - 39.7 Ma) (7) |
| *Promartes olcotti* | Stem to Mustelidae (sister to *Corumictis*) (8) | 26.3-18.5 Ma (PBDB) |

**Table S9.** **Comparisons of the best-fitting evolutionary models for skeletal phenome based on the first seven phylogenetic PC axes.** Rows in boldface type represent the best-fit model as indicated by the lowest ΔAICc score. ΔAICc = AICc minus the minimum AICc between models.

| Model | AICc | ΔAICc | AICcW |
| --- | --- | --- | --- |
| **BM1** | 1039.57 | 112.43 | 0.00 |
| trend | 1045.26 | 118.12 | 0.00 |
| EB | 1054.35 | 127.21 | 0.00 |
| OU1 | 974.04 | 46.90 | 0.00 |
| BMBM_EOT_ | 955.58 | 28.45 | 0.00 |
| OUOU_EOT_ | 969.59 | 42.46 | 0.00 |
| OUBM_EOT_ | 1002.30 | 75.17 | 0.00 |
| **OUBMi_EOT_** | **927.14** | **0.00** | **0.74** |
| BMBM_MMCT_ | 946.88 | 19.74 | 0.00 |
| OUOU_MMCT_ | 964.29 | 37.15 | 0.00 |
| OUBM_MMCT_ | 975.73 | 48.59 | 0.00 |
| OUBMi_MMCT_ | 929.19 | 2.06 | 0.26 |
| sOU2 | 980.79 | 53.66 | 0.00 |
| sOU3 | 989.35 | 62.21 | 0.00 |
| cOU3 | 993.05 | 65.91 | 0.00 |
| cOU4 | 1001.19 | 74.06 | 0.00 |
